# Genome folding principles revealed in condensin-depleted mitotic chromosomes

**DOI:** 10.1101/2023.11.09.566494

**Authors:** Han Zhao, Yinzhi Lin, En Lin, Fuhai Liu, Lirong Shu, Dannan Jing, Baiyue Wang, Manzhu Wang, Fengnian Shan, Lin Zhang, Jessica C. Lam, Susannah C. Midla, Belinda M. Giardine, Cheryl A. Keller, Ross C. Hardison, Gerd A. Blobel, Haoyue Zhang

**Author notes:** Correspondence (G.A.B); (H.Z.).

## Abstract

During mitosis, condensin activity interferes with interphase chromatin structures. Here, we generated condensin-free mitotic chromosomes to investigate genome folding principles. Co- depletion of condensin I and II, but neither alone, triggered mitotic chromosome compartmentalization in ways that differ from interphase. Two distinct euchromatic compartments, indistinguishable in interphase, rapidly emerged upon condensin loss with different interaction preferences and dependence on H3K27ac. Constitutive heterochromatin gradually self-aggregated and co-compartmentalized with the facultative heterochromatin, contrasting with their separation during interphase. While topologically associating domains (TADs) and CTCF/cohesin mediated structural loops remained undetectable, cis-regulatory element contacts became apparent, providing an explanation for their quick re-establishment during mitotic exit. HP1 proteins, which are thought to partition constitutive heterochromatin, were absent from mitotic chromosomes, suggesting, surprisingly, that constitutive heterochromatin can self-aggregate without HP1. Indeed, in cells traversing from M- to G1-phase in the combined absence of HP1α, HP1Π and HP1γ, re-established constitutive heterochromatin compartments normally. In sum, “clean-slate” condensin-deficient mitotic chromosomes illuminate mechanisms of genome compartmentalization not revealed in interphase cells.

## INTRODUCTION

The mammalian genome is highly organized in 3D which includes formation of compartments, topologically associating domains (TADs), and chromatin loops. While the compartmentalization of the active (compartment A) and repressive genome (compartment B) is linked to transcriptional activity and histone modifications^1^, the mechanisms driving the aggregation and separation of compartments remain unclear. At finer scales, there are two major categories of chromatin loops: structural loops which are mediated by CTCF/cohesin driven loop extrusion, and loops between cis-regulatory elements (CREs) such as enhancers or promoters ^2^. It has been demonstrated that CRE loops can form independently of CTCF and cohesin ^2,3^. However, forces that mediate CRE loop formation are still unclear.

All architectural features are highly dynamic throughout the cell cycle ^2,4-6^. Upon entering mitosis, A/B compartments, TADs and chromatin loops are disrupted with concomitant loading of the condensin ring complexes ^7^. Despite these dramatic changes, weak domain-like (triangles along the diagonal) structures remain detectable on mitotic chromosomes ^2^. The mechanisms underlying these residual mitotic folding patterns are unknown. In condensin-deficient cells, the disappearance of A/B compartments and TADs is delayed during mitotic entry ^7,8^. Morphologically, depletion of condensin abrogates the cylindrical shape of mitotic chromosomes and triggers intermingling of individual chromatids ^8^. The total volume of mitotic chromosomes is mostly unchanged upon condensin loss as assessed by three-dimensional electron microscopy ^8^. These and other observations suggest that in addition to condensins, other forces may shape the conformation of mitotic chromosomes ^7^.

Mitotic cells differ from those in interphase in several aspects: (1) the nuclear envelope breaks down and the nuclear lamina meshwork is disassembled, allowing the mixture of nucleoplasm and cytoplasm; (2) transcription factors, co-regulators, chromatin modifiers and other chromatin binding partners are largely evicted from the chromosomes ^9^; (3) transcription ceases globally ^2,10^ and (4) cohesin is unloaded from the chromosome arms ^2^. Many of the features that are lost in mitosis have been proposed to modulate genome architecture in interphase cells. Therefore, investigating the conformation of condensin-deficient mitotic chromosomes, which are devoid of the above influences, may reveal the roles of cellular activities (e.g., transcription) and proteins in chromosome structure regulation, and provide insights into folding principles, such as compartmentalization of the genome, that may be hidden in interphase cells.

Here, we exploit a system in which a major force of mitotic chromosome organization has been removed by acute inactivation of condensin. We find that despite lacking transcription and association with numerous chromatin binding partners, mitotic chromosomes maintain the capacity to undergo intricate compartmentalization in ways that can be obscured in interphase chromatin. Unexpectedly, HP1 proteins are dispensable for constitutive heterochromatin compartment formation not only in condensin depleted mitotic cells but also during re-entry into G1-phase. Taken together, our results highlight the power of studying mitotic chromatin and cell cycle transition states to reveal forces driving genome compartmentalization.

## RESULTS

### Acute depletion of condensin in mitotic chromosomes

To characterize the intrinsic drivers of chromatin folding, we sought to acutely inactivate both type I and II condensin in mitotic cells by targeting their shared subunit SMC2 ^7,11^. Using CRISPR/Cas9 directed genome editing, we inserted a minimal auxin-inducible degron (mAID) homozygously at the *Smc2* locus in G1E-ER4 cells, a well characterized murine erythroblast line (Fig. 1a) ^2,12^. The resulting G1E-ER4:SMC2-mAID-mCherry cells displayed normal chromosome condensation and correct localization of the SMC2 fusion protein (Extended Data Fig. 1a). Thus, SMC2 protein function was not measurably perturbed by its modification. Incubation of cells with auxin depleted SMC2 within 1 hour (Extended Data Fig. 1b). Prolonged SMC2 depletion dramatically shifted cell cycle distribution and arrested cell growth (Extended Data Fig. 1b, c), confirming a critical role of condensin for mitotic progression ^7^. The rapidity of SMC2 degradation enabled removal of SMC2 specifically during mitosis. We enriched prometaphase cells by nocodazole induced cell cycle arrest, followed by auxin treatment for 0.5h, 1h, 4h or 8h (Fig. 1b). Light microscopy revealed a progressive loss of the “rod-like” shape of mitotic chromosomes, along with extensive entanglements among individual chromatids, confirming the essential role of condensin in the maintenance of mitotic chromosome morphology (Extended Data Fig. 1d, e). Hence, mitotic chromosomes undergo major restructuring after being depleted of condensin ring complexes.

**Figure 1.**
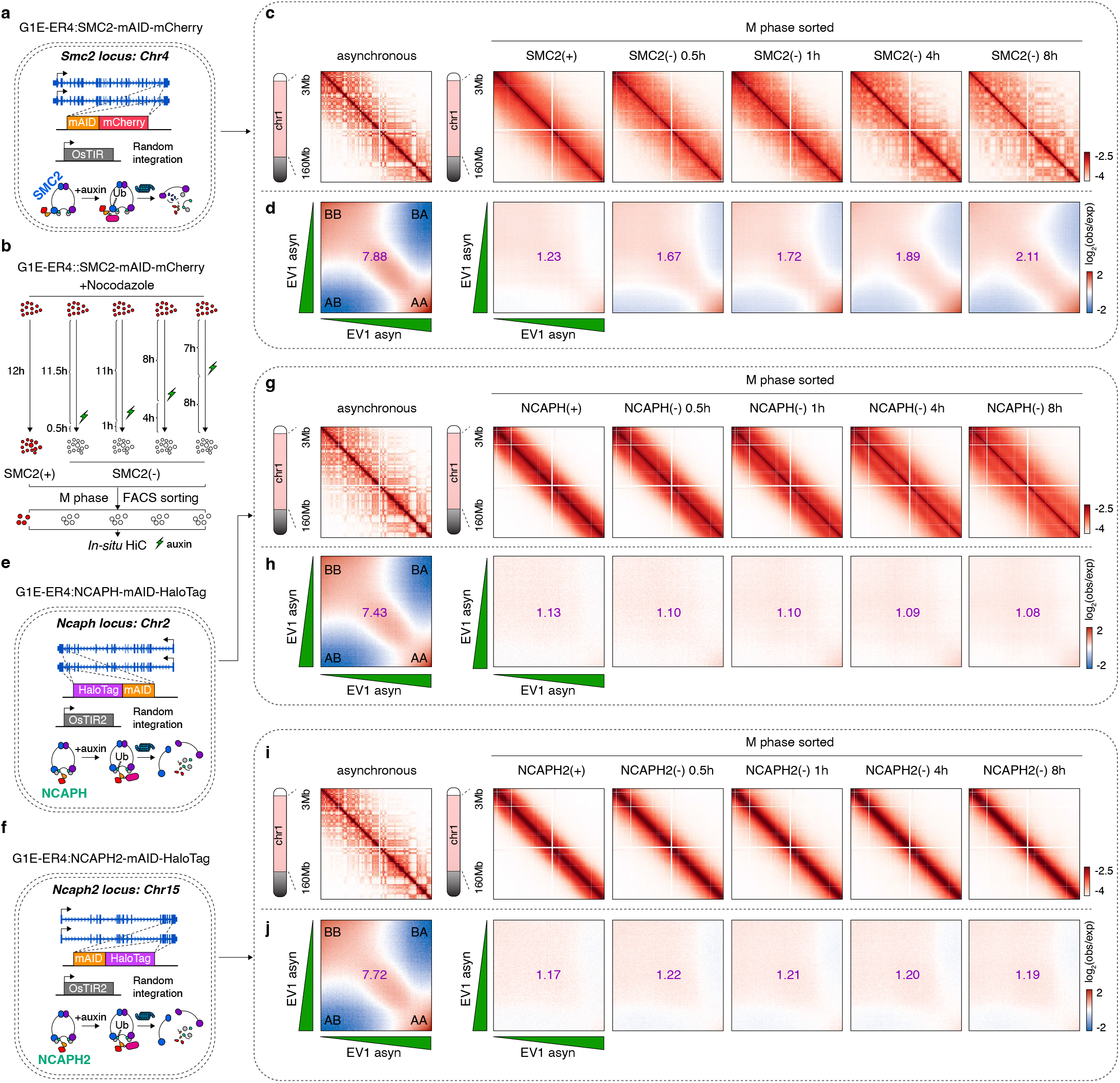
Progressive compartmentalization of the condensin-depleted mitotic chromosomes. **a**, Schematic illustration showing the homozygous insertion of mAID-mCherry tag to the C terminus of endogenous SMC2 protein. **b**, Experimental design, showing the strategy of prometaphase arrest and auxin treatment. **c**, KR balanced Hi-C contact matrices showing the global re-configuration of mitotic chromosomes after condensin removal. Hi-C maps of asynchronous cells were shown as control. Bin size: 100kb. **d**, Saddle-plots showing the progressive compartmentalization of mitotic chromosomes after condensin removal. Compartmental strength is shown for each sample. **e**, Schematic illustration showing the homozygous insertion of mAID-Halotag to the C terminus of endogenous NCAPH protein. **f**, Schematic illustration showing the homozygous insertion of mAID-Halotag to the C terminus of endogenous NCAPH2 protein. **g**, KR balanced Hi-C contact matrices of mitotic chromosomes after condensin I removal. Bin size: 100kb. **h**, Saddle-plots showing lack of compartmentalization during mitosis after condensin I removal. **i**, KR balanced Hi-C contact matrices of mitotic chromosomes after condensin II removal. Bin size: 100kb. **j**, Saddle-plots showing lack of compartmentalization during mitosis after condensin II removal.

### TADs and structural loops failed to reform in the condensin-depleted mitotic chromosomes

To characterize the folding patterns of condensin-depleted mitotic chromosomes, we purified nocodazole arrested prometaphase cells before and after auxin treatment by fluorescence activated cell sorting (FACS) on the basis of pMPM2 (mitotic-specific marker) and surveyed their chromatin architecture using *in-situ* Hi-C (Fig. 1b; Extended Data Fig. 2a; Supplementary table1)^13^. Asynchronously growing cells were processed as “interphase” control. Collectively, *in-situ* Hi-C yielded ∼2 billion valid interaction pairs, with high concordance among biological replicates (Extended Data Fig. 2b). In the absence of condensin, *trans*-chromosomal interactions were progressively gained, corroborating the gradually increased entanglements of individual chromatids (Extended Data Fig. 2c, d).

The triangular structures along the diagonal of Hi-C contact matrices can represent TADs formed by cohesin mediated loop extrusion or compartmental domains. Since cohesin and to varying degrees CTCF are evicted from mitotic chromosomes ^2,4,14^, we first examined which if any CTCF/cohesin dependent structures are observable in the condensin-depleted mitotic chromosomes. Using the “Arrowhead” tool in combination with local minima of insulation score profiles, we identified 2,914 domains in asynchronous cells (Extended Data Fig. 3a; Supplementary table2). Remarkably, 1,247 domains were called in condensin-deficient mitotic cells. However, these domains are mostly distinct from the domains called in interphase, with only 316 domains overlapping been the two conditions (Extended Data Fig. 3b).

Arrested loop extrusion by cohesin at CTCF sites, leaves signature “corner dots” at the summits of the domain triangles, and a substantial fraction (∼34.6%) of domains present in interphase chromatin displayed such dots (Extended Data Fig. 3c-e). By contrast, domains detected in condensin depleted mitotic cells rarely exhibited corner-dots (Extended Data Fig. 3f-h), suggesting they were formed by mechanisms independent of loop extrusion. We speculate that the residual corner-dot signals may reflect micro-compartmental interactions instead of CTCF/cohesin mediated loops. In agreement with these findings, the CTCF/cohesin mediated structural loops were undetectable in condensin-depleted mitotic chromosomes as indicated by aggregated peak analysis (APA) (Extended Data Fig. 3i). Finally, ChIP-seq experiments revealed a complete loss of cohesin positioning in mitotic cells without condensin (Extended Data Fig. 3j), similar to our previous findings in condensin replete mitotic cells ^2^. Together, these results indicate that condensin deficient mitotic chromosomes have the ability to form domain-like structures, but these are distinct from CTCF/cohesin dependent TADs and structural loops.

### Mitotic chromosomes maintain the ability to compartmentalize in the absence of condensin

We next explored the compartmentalization of mitotic chromosomes after condensin loss. Consistent with prior reports, compartments were almost entirely missing in control mitotic cells (Fig. 1c). Remarkably, a checkerboard pattern reflective of compartmentalization started to emerge as soon as 30min after auxin treatment, as assessed by visual inspection of individual chromosomes and genome-wide saddle plots (Fig. 1c, d). Mitotic compartments strengthened and expanded progressively over time (Fig. 1c, d; Extended Data Fig. 4a, b). Despite being attenuated, the patterns of compartmentalization during mitosis were remarkably similar to those in interphase (Fig. 1c). Importantly, *in-silico* simulation demonstrated that even a 20% contribution of interphase signals failed to reproduce the mitotic compartments, suggesting that compartmentalization was not caused by the admixture of interphase cells (Extended Data Fig. 4c). To rule out the possibility that mitotic compartments were nocodazole-induced artifacts, we incubated asynchronous cells with auxin for 4 hours to degrade SMC2 and purified mitotic cells through FACS. Mitotic cells without auxin treatment were sorted in parallel as control (Extended Data Fig. 5a, b). Hi-C experiments uncovered conventional “structure-less” mitotic chromosomes in untreated control cells (Extended Data Fig. 5c). Importantly, in auxin treated cells, mitotic chromosomes reproducibly exhibited clear plaid-like compartmentalization patterns (Extended Data Fig. 5c). Compartment strengths were comparable to cells that had undergone nocodazole-induced mitotic arrest as evidenced by highly correlated eigenvector1 values (Extended Data Fig. 5d). Interestingly, loss of condensin in interphase cells increased *trans*-interactions among peri-centromeric regions of different chromosomes (data not shown), supporting a prior report ^15^. However, intra-chromosomal compartments in condensin depleted interphase cells remained unchanged (Extended Data Fig. 5e). In sum, a major re-configuration of mitotic chromosome architecture takes place upon loss of condensin, gradually re-building compartmentalization patterns that resemble those in interphase cells. The ability to re-form compartments for mitotic chromosomes may partially explain the maintenance of chromosome compaction in the absence of condensin ^8^.

### Condensin I and II both suppress mitotic compartments

We next asked which type of condensin counteracts mitotic compartments. We generated G1E-ER4 cell lines with mAID endogenously tagged to the condensin I specific kleisin subunit NCAPH or the condensin II specific kleisin subunit NCAPH2 (Fig. 1e, f; Extended Data Fig. 6a- c). In contrast to cells lacking both condensins, cells devoid of either condensin I or II maintained the rod-shaped morphology of individual mitotic chromosomes and displayed normal cell cycle distribution (Extended Data Fig. 6d-f). In line with this observation, inactivating either condensin I or II for up to 8 hours did not increase *trans*-chromosomal interactions (Extended Data Fig. 7a- d).

After 1 hour of NCAPH deletion (condensin II active), the Hi-C contact matrices and contact probability decay curves (*P(s)* curve) of mitotic chromosomes started to display a parallel secondary diagonal at ∼10-20Mb of genomic separations, representing periodically increased interaction frequencies along the entire chromosome (Fig. 1g; Extended Data Fig. 7e). Upon extended NCAPH loss (∼8h), the secondary diagonal progressively migrated to larger genomic distances (∼40Mb) (Fig. 1g; Extended Data Fig. 7e), implying constant extrusion activity by condensin II to promote longitudinal shortening of mitotic chromosomes. In NCAPH2 depleted (condensin I active) mitotic cells, we observed a gradual reduction of contacts at ∼10-30Mb genomic separations (Fig. 1i; Extended Data Fig. 7f). Chromatin spread analysis revealed long and “curly” mitotic chromosomes resembling those in prophase cells (Extended Data Fig. 7g). Unlike deleting SMC2, removal of either NCAPH or NCAPH2 for up to 8 hours failed to induce the plaid-like pattern of compartments (Fig. 1g, i). Saddle plots revealed no marked changes in compartment strengths compared to controls (Fig. 1h, j). Taken together, our data indicate that condensin I and II govern different aspects of mitotic chromosome morphogenesis, supporting a prior study^7^. Importantly, both complexes can suppress mitotic compartmentalization.

### Intricate patterns of compartmentalization in condensin-depleted mitotic chromosomes

Comparing mitotic vs. interphase compartmentalization patterns may inform principles of chromatin organization that are overlooked in interphase cells. We segmented the genome into 25kb bins and performed eigenvector decomposition for the nocodazole arrested condensin-deficient mitotic chromosomes (4h auxin) and the asynchronous control cells. In asynchronous cells, ∼41.3% of the genome displayed positive EV1 values. Yet, this percentage dropped to ∼27.7% during mitosis, implying broad compartment switches (Fig. 2a). Accordingly, we detected small B-like regions emerging within conventional A-type compartments in condensin-depleted mitotic chromosomes, indicating a more fragmented pattern of compartmental interactions than in interphase chromatin (Fig. 2b).

**Figure 2.**
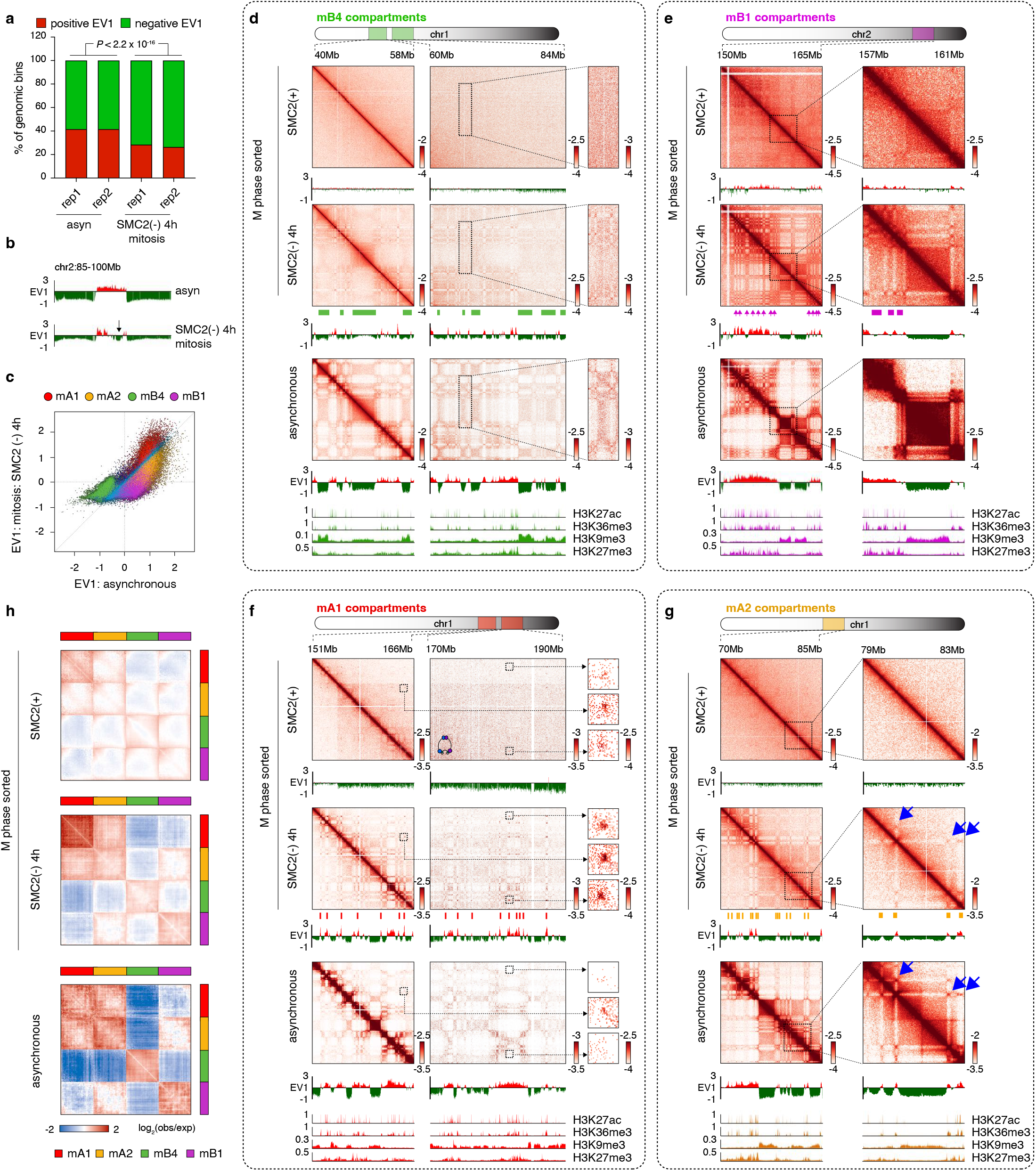
Intricate patterns of mitotic compartments. **a**, Bar graph showing the composition of 25kb genomic bins with positive or negative EV1 values in condensin-deficient mitotic and asynchronous control cells. *P* values were calculated using a two-sided Fisher’s exact test. **b**, Browser tracks showing the EV1 values of asynchronous and condensin-depleted mitotic cells (4h). Arrow indicates the compartment switch. **c**, Scatter plot showing the EV1 values of 25kb genomic bins in asynchronous control cells (*x*-axis) against condensin-deficient mitotic cells (*y*-axis). Bins were color coded based on their compartment assignment. **d**, KR balanced Hi-C contact matrices showing a representative (chr1:40-58Mb vs. chr1:60-84Mb) homotypic mB4 interactions in condensin-deficient (4h) mitotic cells. Bin size: 25kb (10kb for enlarged view). mB4 compartments were indicated by green bars. Browser tracks of EV1 values for each condition as well as H3K27ac, H3K36me3, H3K9me3 and H3K27me3 from asynchronously growing cells were shown. **e**, Similar to (**d**), showing representative mB1- type compartments and highlighting a conversion of interaction patterns between mB1 and mB4 compartments from asynchronous to condensin-deficient mitotic cells. mB1 compartments were indicated by purple bars. **f**, KR balanced Hi-C contact matrices showing a representative (chr1:151-166Mb vs. chr1:170-190Mb) strengthening of the mA1 compartmental interactions in condensin-deficient (4h) mitotic cells. Bin size: 25kb (10kb for enlarged view). mA1 compartment was indicated by red bars. **g**, Similar to (**f**), showing representative examples (chr1:70-85Mb) of mA2 compartmental interactions. mA2 compartment was indicated by yellow bars. Homotypic interactions of mA2 was highlighted by blue arrows. **h**, Attraction-repulsion plots showing the homotypic and heterotypic interactions among four different compartments: mA1, mA2, mB4 and mB1. Plots for control mitotic, condensin-deficient (4h) mitotic and asynchronous cells were shown.

Using unsupervised *k*-means clustering, we partitioned the genome into three groups. Group I (∼49.5%) and II (∼33%) displayed increased and decreased EV1 values, respectively, indicating that mitosis-specific compartmentalization differ from that in interphase (Extended Data Fig. 8a). Group III (∼17.5%) showed no apparent change. Notably, groups I and II were each decorated with both active and repressive histone marks during interphase, implying that they may be further sub-categorized (Extended Data Fig. 8b-h). Combining chromatin associated features and the EV1 clustering results, we identified four types of compartments, mA1, mA2, mB1 and mB4, (m for mitosis) with distinct interaction preferences in condensin-depleted mitotic chromosomes (Extended Data Fig. 8i; Supplementary table3). Hence, condensin depletion in mitotic chromatin uncovered novel patterns of chromatin compartmentalization.

### Compartmentalization of the repressed genome in condensin-depleted mitotic cells

In the inactive chromatin, we identified two types of compartments: mB1 and mB4 (Fig. 2c; Extended Data Fig. 8i). The latter was termed as mB4 because it was epigenetically similar to sub-compartment B4 previously identified in HCT116 and GM12878 cells ^16,17^. mB4 occupied ∼38.4% of the genome and was enriched for the constitutive heterochromatin mark H3K9me3 during interphase (Extended Data Fig. 8i, 9, 10a). Hi-C contact maps indicate that in condensin deficient mitotic cells, mB4-mB4 aggregation achieved ∼30% of that found in interphase cells (Fig. 2d; Extended Data Fig. 9). Whether even higher mB4-mB4 aggregation were attainable if condensin was removed for a longer duration or whether qualitative differences in the chromatin limit further aggregation remains an open question. Nevertheless, it is remarkable that constitutive heterochromatin was able to self-aggregate during mitosis, despite at reduced levels, given that the nuclear lamina is disassembled and the H3K9me3 reader proteins such as HP1 are displaced from chromosomes (see below).

The mB1 compartment represents facultative heterochromatin marked by H3K27me3, accounting for ∼20.9% of the genome (Extended Data Fig. 8i, 9, 10a). In interphase cells, mB1 was frequently (∼84.7%) separated from constitutive heterochromatin, displaying positive EV1 values and categorized as the conventional A compartments (Fig. 2c, e). Strikingly, ∼61% of mB1 compartment displayed a flip of EV1 direction during mitosis, far exceeding the rest of the genome, suggesting a major switch of interaction preferences (Extended Data Fig. 10b). Indeed, in the condensin-depleted mitotic chromosomes, mB1 displayed mild attraction to mB4 at the regions surrounding centromeres and telomeres, as reflected by relatively high interaction frequencies (Fig. 2e; Extended Data Fig. 9, 10c-e). Together, our data demonstrate that facultative heterochromatin and constitutive heterochromatin have the ability to co-compartmentalize during mitosis.

### Compartmentalization of the active genome in condensin-depleted mitotic cells

Two active mitotic compartments (mA1 and mA2) emerged from our analysis, both of which were decorated by active histone marks, but fell into group I and II regions, respectively (Fig. 2c; Extended Data Fig. 8b-f, i). In the case of mA1, we observed increased EV1 values and a striking gain of homotypic interactions in the condensin-deficient mitotic cells (Fig. 2c, f; Extended Data Fig. 9). It is noteworthy that these interactions were so pronounced that they were even detectable in the condensin replete control mitotic cells (Extended Data Fig. 11a). Thus, the aforementioned residual structures in normal mitotic cells may originate from strong mA1 homotypic interactions. In contrast, the mA2 compartment showed lower EV1 values and a dramatic reduction of homotypic interactions during mitosis when compared to interphase cells (Fig. 2c, g). In interphase cells, mA2 was less enriched with active histone marks and displayed lower transcription activity (Extended Data Fig. 11b). To our surprise, despite the dramatic difference in the condensin-deficient mitotic cells, during interphase, EV1 values for mA1 and mA2 were very similar rendering them virtually indistinguishable (Extended Data Fig. 11c, d). To better understand the spatial arrangements of mA1 and mA2, we generated attraction-repulsion plots by re-ordering the rows and columns of the 25kb binned Hi-C contact matrices such that genomic bins belonging to the same type of compartment were grouped together. We found that mA1 and mA2 were clearly segregated during mitosis in the absence of condensin, but virtually inseparable in interphase cells, showing similarly strong intra- and inter-compartmental interactions (Fig. 2h). As expected, mA1 repulses both mB4 and mB1 compartments (Fig. 2h). Intriguingly, mA2 is excluded from mB4 but mildly attracted to mB1, despite their classic euchromatic epigenetic signatures, suggesting a qualitative difference from mA1 (Fig. 2h). It is worth mentioning that these specific patterns of euchromatin and heterochromatin compartmentalization were also observed in the “asyn-sorted” condensin-free mitotic (but not interphase) cells (Extended Data Fig. 5f-j). In sum, using condensin-depleted mitotic chromosomes as an architectural platform, we unveiled a new complexity of active chromatin organization and discovered two types of active compartments.

### Distinct compartmentalization in extrusion-free mitotic and interphase chromatin

Cohesin-driven loop extrusion interferes with chromatin compartmentalization in interphase cells ^18,19^. This raises the question whether extrusion-free chromatin in condensin-deficient mitotic cells and in cohesin depleted interphase cells compartmentalize similarly. Exploration of this question may shed light on the roles of cellular processes, such as gene expression and transcription factor occupancy on chromatin compartmentalization, given their differences between mitotic and interphase cells. To address this, we generated Hi-C datasets in G1E-ER4:SMC3-mAID-mCherry cells, which contained a mAID-mCherry tag homozygously fused to the C-terminus of endogenous SMC3, a cohesin core subunit (Extended Data Fig. 12a-c). As expected, depletion of cohesin through auxin treatment for 4 hours in interphase cells triggered a widespread disruption of domain boundaries and structural loops (Extended Data Fig. 12d-g). Eigenvector decomposition uncovered increments of EV1 values for both mA1 and mA2 compartments, suggesting a general gain of euchromatin self-aggregation upon cohesin loss (Extended Data Fig. 12h). Therefore, the inhibitory effect of cohesin on euchromatin compartmentalization did not distinguish mA1 from mA2 in interphase cells. Visual examination of individual Hi-C contact maps and genome-wide comparisons of compartment strengths for the “extrusion-free” interphase and mitotic genome revealed that the latter displayed slightly stronger mA1 but dramatically attenuated mA2 homotypic interactions (Extended Data Fig. 12i-k), suggesting that mA2 (but not mA1) compartmentalization was impaired during mitosis. In addition, the attraction between mB1 and mB4 compartments observed at telomere-proximal regions in the condensin-deficient mitotic chromosomes was missing in interphase chromatin free of cohesin (Extended Data Fig. 12l). Our data demonstrate that when released from condensin, mitotic chromosomes, which presumably lack PolII binding, are capable of strong euchromatin self-aggregation, highlighting that compartmentalization of mA1 active chromatin can be independent of transcription as well as the numerous proteins that are evicted from mitotic chromatin.

### Active and inactive chromatin compartmentalize with distinct dynamics

Dissecting the kinetics of mitotic chromatin compartmentalization may reveal forces that partition the genome. Our time course mitotic Hi-C datasets showed that despite globally weaker compartmentalization at early time points, the homotypic mA1 interactions started to emerge at only 30min after auxin treatment and approached plateau intensity within just 1h (Fig. 3a, b; Extended Data Fig. 13). By contrast, mB4 compartments were barely detectable after 30 min of condensin loss and only slowly intensified over time (Fig. 3a, b). Interestingly, mA2 and mB1 displayed comparable kinetics, slower than that of mA1, but faster than mB4 (Fig. 3b). The difference in rates of mA1 and mB4 compartment formation indicates that the compartmentalization process of euchromatin and constitutive heterochromatin can be uncoupled.

**Figure 3.**
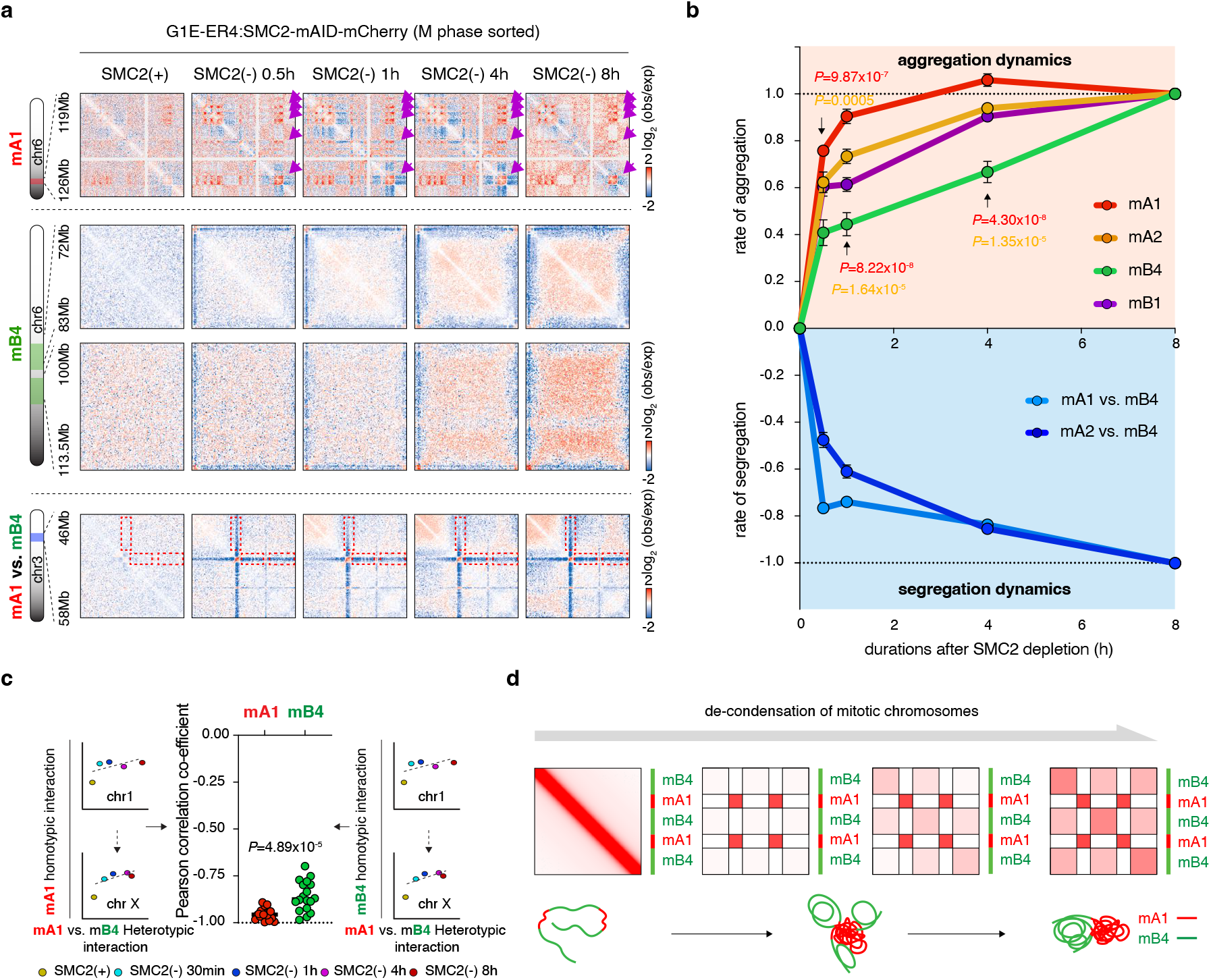
Dynamic attraction and segregation of compartments during mitosis upon condensin loss. **a**, Upper panel: KR balanced Hi-C contact matrices (chr6:119-126Mb) showing rapid emergence of mA1 homotypic interactions in mitosis upon condensin loss. Bin size: 25kb. Middle panel: KR balanced Hi-C contact matrices (chr6:72-83Mb vs. chr6:100-113.5Mb) showing the delay of the formation of mB4 homotypic interactions. Bin size: 100kb. Lower panel: KR balanced Hi-C contact matrices (chr3:46-58Mb) showing rapid segregation of mA1 from B4 compartments in mitosis after condensin loss. Bin size: 25kb. Depletion of interactions was indicated by red dotted boxes. **b**, Line plots showing the differential reformation kinetics for mA1, mA2, mB1 and mB4 as well as the repulsion kinetics between mA1 vs. mB4 and mA2 vs. mB4. Error-bar denotes SEM (n=18 chromosomes). Statistic tests were performed for comparison of the aggregation dynamics between mA1 vs. mB4 compartments (red) and mA2 vs. mB4 compartments (yellow). *P* values were calculated using a two-sided Wilcoxon signed-rank test. **c**, Left and right: schematics showing how correlations are computed between mA1 or mB4 aggregation and mA1 vs mB4 segregation over time. Middle: dot plot showing the Pearson correlation coefficients between mA1 or mB4 compartment strength and mA1 vs. mB4 segregation. Each dot represents an individual chromosome (n=18). *P* values were calculated using a two-sided Wilcoxon signed-rank test. **d**, Schematic showing the fast aggregation of mA1 and rapid exclusion of mA1 from mB4 and delayed self-attraction of mB4, implying that mA1 compartment initiates the formation of checkerboard compartmental interaction pattern in mitotic chromosomes upon condensin loss.

Formation of the checkerboard pattern of compartmental interactions relies on both the self-aggregation of each type of compartment as well as the mutual separation of compartments with opposite states. It is currently unclear whether the separation of two different types of compartments occurs concomitantly as they both aggregate. The dramatic difference between mA1 and mB4 aggregation kinetics in condensin-depleted mitotic chromosomes offered the opportunity to investigate this question. At 30min following condensin loss, we observed a rapid separation of mA1 from mB4 (Fig. 3a, b; Extended Data Fig. 13). Segregation of mA1 from mB4 anti-correlated well with mA1 aggregation but less so with mB4 (Fig. 3c). Our data therefore support a model in which upon condensin loss, the euchromatin compartments quickly self-aggregate and simultaneously exclude constitutive heterochromatin, which in turn aggregates at later time points (Fig. 3d). Our data highlight an important role of active genome shaping chromatin compartments.

### H3K27ac facilitates mA1 compartmentalization during mitosis

We next sought to investigate potential correlation between histone modification and mitotic compartmentalization. We started by profiling via ChIP-seq active chromatin marks, including RNA Polymerase II, H3K27ac, H3K36me3 and H3K4me1 in the condensin-deficient mitotic cells. In the absence of condensin, mitotic chromosomes were completely devoid of RNA PolII, ruling out transcription as the driving force of mitotic compartmentalization (Extended Data Fig. 14a-c). The mA1 compartment was more enriched for H3K27ac, H3K36me3 and H3K4me1 than mA2 (Extended Data Fig. 14c). Interestingly, we observed a more dramatic difference of H3K27ac between mA1 and mA2 than that of the other two marks (Extended Data Fig. 14c), suggesting that differential H3K27ac signal intensity may underline the distinct interaction preferences of mA1 and mA2 during mitosis.

To test whether H3K27ac contributes to euchromatin compartmentalization during mitosis, we ablated mitotic H3K27ac by treating the cells with the selective p300/CBP inhibitor A485 for 3 hours prior to nocodazole arrest and SMC2 degradation (Fig. 4a). ChIP-seq confirmed a substantial reduction of H3K27ac in the condensin-deleted mitotic cells after A485 treatment (Fig. 4b; Extended Data Fig. 15a, b). Hi-C analysis revealed that global mitotic chromatin compartmentalization was not measurably perturbed (Extended Data Fig. 15c). Notably, A485 treatment induced a mild but significant reduction of the EV1 values for mA1 compartments (Fig. 4c). Unsupervised *k-means* clustering uncovered that ∼62% of mA1 was unaffected by A485 treatment (mA1-U, U for “unchanged EV1”) (Fig. 4d, e, g). However, ∼25.7% of mA1 compartment was sensitive to H3K27ac deletion (mA1-D, D for “decreased” EV1), reproducibly displaying decreased EV1 values and reduced compartmental interactions (Fig. 4d, e, f, i; Extended Data Fig. 15d, e). Interestingly, A485 treatment also induced a slight increase of compartmentalization for ∼12.3% of mA1 (mA1-I, I for “increased” EV1) (Fig. 4d, e, h, i; Extended Data Fig. 15d, e). It is worth mentioning that mA1-D compartment displayed higher levels of H3K27ac in the condensin-deficient mitotic chromosome, compared to mA1-I and mA1- U (Fig. 4j). Taken together, these data demonstrate that the H3K27ac mark participates in the compartmentalization of a fraction of mA1 in the condensin-deficient mitotic cells.

**Figure 4.**
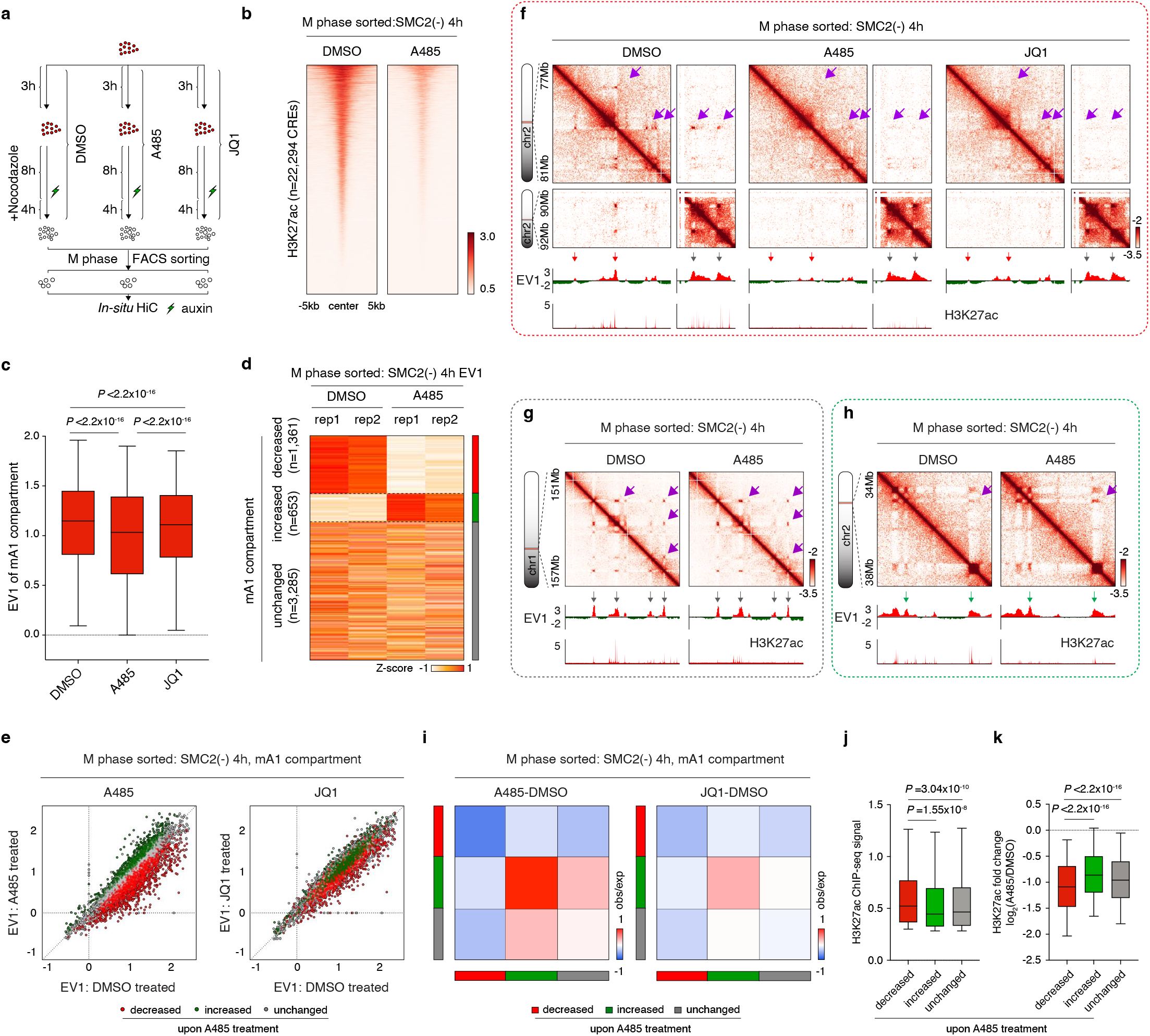
H3K27ac contributes to mA1 compartmentalization in the condensin-deficient mitotic cells. **a**, Schematic showing A485 and JQ1 treatment strategy. **b**, Density heatmaps showing loss of H3K27ac signals in the condensin-deficient (4h) mitotic chromosomes after A485 treatment. **c**, Box-plots showing the EV1 values of each 25kb genomic bins (n=5,299) of mA1 compartment in the condensin-deficient mitotic cells upon DMSO, A485 or JQ1 treatment. For all box plots, central lines denote medians; box limits denote 25th–75th percentile; whiskers denote 5th–95th percentile. *P* values were calculated using a two-sided paired Wilcoxon signed-rank test. **d**, Heatmap showing the clustering result of EV1 values between DMSO and A485 treated samples. **e**, Left panel: Scatter plot showing the EV1 values of mA1 25kb genomic bins in DMSO treated (*x*-axis) against A485 treated (*y*-axis) condensin-deficient (4h) mitotic cells. Right panel: Scatter plot showing impact of JQ1 on mA1 compartment. Bins were color coded based on their response to A485 treatment. **f**, KR balanced Hi-C contact matrices showing representative (chr2:77-81Mb vs. chr2:90-92Mb) mA1-D compartments in the DMSO, A485 or JQ1 treated condensin-deficient mitotic cells. Compartmental interactions were indicated by purple arrows. Bin size: 25kb. Browser tracks of EV1 values for each condition as well as H3K27ac were shown. Red arrows indicate the reduction of EV1 values upon A485 treatment. Gray arrows indicate unaffected regions. **g**, Similar to (**f**), showing representative mA1-U compartmental interactions. **h**, Similar to (**f**), showing representative mA1-I compartmental interactions. **i**, Left panel: Heatmap showing the differential interaction strengths among mA1-D, mA1-I and mA1-U compartments upon A485 treatment compared to DMSO treated controls. Right panel: Heatmap showing the differential interaction strengths among mA1-D, mA1-I and mA1-U compartments upon JQ1 treatment. **j**, Box-plots H3K27ac ChIP-seq signal strength in the condensin-deficient mitotic cells for mA1-D (n=1,361), mA1-I (n=653) and mA1-U (n=3,285) compartments. For all box plots, central lines denote medians; box limits denote 25th–75th percentile; whiskers denote 5th–95th percentile. *P* values were calculated using a two-sided Wilcoxon signed-rank test. **k**, Similar to (**j**) Box-plots showing the log_2_ fold change of H3K27ac signals within the mA1-D (n=1,880), mA1-I (n=861) and mA1-U (n=4,518) compartments.

Since BET proteins such as BRD4 bind to acetylated histones during mitosis, we next asked whether BET protein plays a role in H3K27ac mediated mA1 compartmentalization ^20^. To address this, we treated cells with the BET protein specific inhibitor JQ1 for 3 hours prior to mitotic enrichment and SMC2 depletion (Fig. 4a). We observed significant reduction of mA1 EV1 values in the JQ1 treated cells (Fig. 4c). However, the effect size was milder compared to those treated by A485 (Fig. 4c, e; Extended Data Fig. 15d, f). In agreement with these data, BET protein inhibition also reduced compartmental interactions among mA1-D regions, but to a lesser extent compared to H3K27ac deletion (Fig. 4f, i; Extended Data Fig. 15e, g). Hence, BET proteins contribute to mA1 compartmentalization during mitosis. These results are reminiscent with a previous report that BRD2 compartmentalizes the active genome in interphase ^21^.

### Rapid formation of CRE contacts in condensin-depleted mitotic chromosomes

Contacts among cis-regulatory elements (CREs) may rely on transcription factor binding. Since transcription factors tent to be evicted from mitotic chromosomes, we next asked whether CRE contacts may still emerge during mitosis upon condensin loss. To address this question, we identified 6,825 CRE contacts (see methods) that were present in interphase cells, setting the distances between CRE loop anchor pairs to be less than 300kb to avoid confusion with euchromatic compartmental interactions (Fig. 5a). Based on APA signals, the CRE contacts were nearly undetectable in the control mitotic cells as expected (Fig. 5b, c). Already after 30min of auxin treatment, focal contacts between CRE pairs emerged and plateaued in strength thereafter (Fig. 5b, c). Deleting either type I or type II condensin alone failed to recapitulate this observation, suggesting an inhibitory role of both complexes (Fig. 5d-g). Interestingly, deletion of H3K27ac at CREs by A485 treatment led to a mild decrease of their connectivity in mitotic chromosomes, suggesting that H3K27ac may partially facilitate CRE communication during mitosis (Extended Data Fig. 16a-d). However, JQ1 treatment failed to show this effect, suggesting that BET proteins are dispensable for CRE contacts in this context, reminiscent of a prior report that interphase CRE contacts are unchanged in the absence of BET proteins (Extended Data Fig. 16b-d) ^22^. Taken together, despite the absence of transcription and numerous chromatin associated factors, mitotic chromosomes maintain the capacity to form CRE contacts.

**Figure 5.**
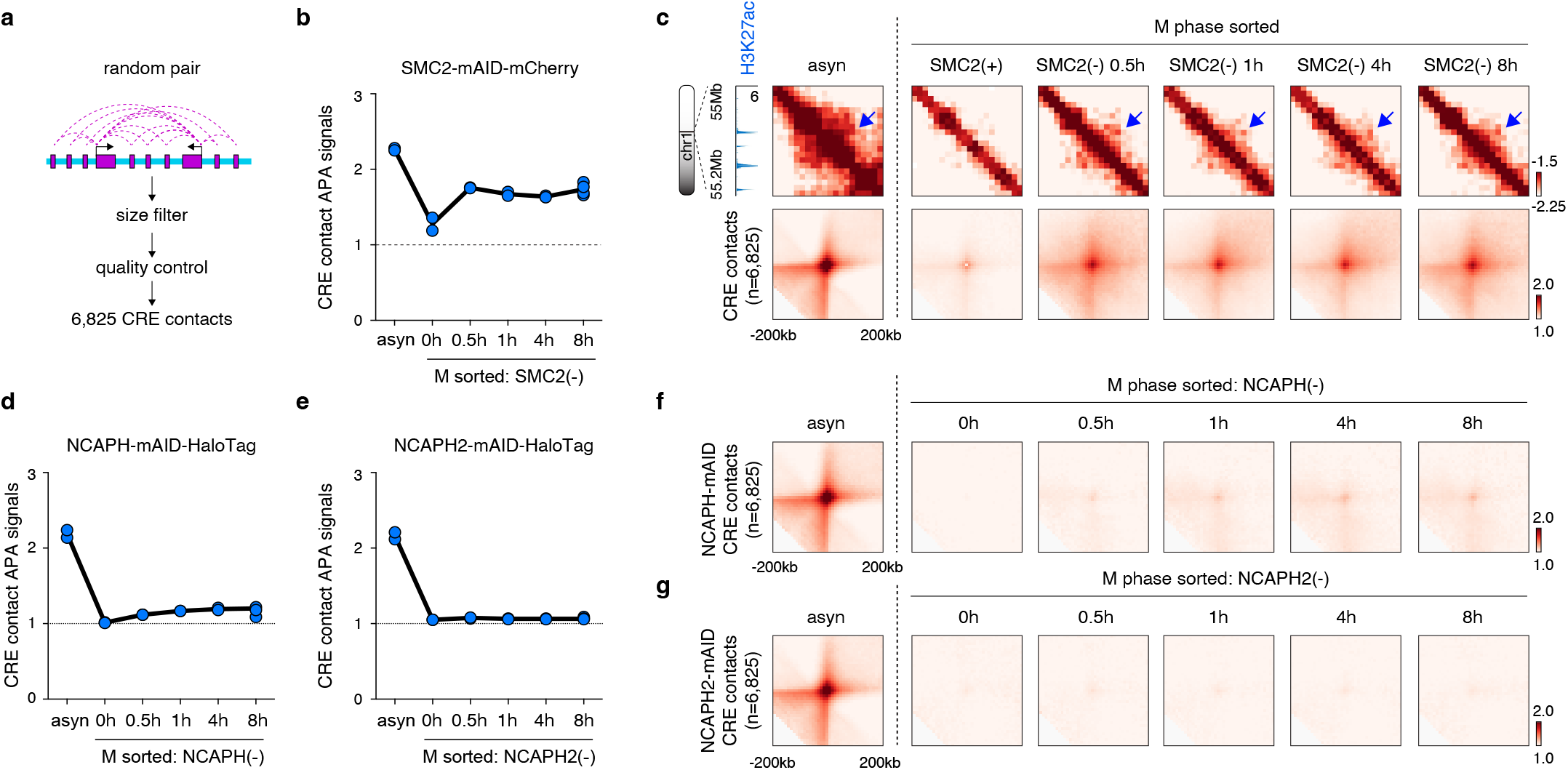
Formation of CRE contacts in mitotic chromosomes upon condensin loss. **a**, Schematic showing stratification of the 6,825 interphase CRE contacts. **b**, Line plot showing the APA signals of CRE contacts in condensin-deficient mitotic cells. **c**, Upper panel: representative KR balanced Hi-C contact matrices (chr1:55-55.2Mb) showing the emergence of focalized signal between CREs in mitotic cells upon condensin loss. Bin size: 10kb. Lower panel: APA plots of CRE contacts in asynchronous and mitotic cells with or without condensin. **d**, Line plot showing the APA signals of CRE contacts in condensin I-deficient mitotic cells. **e**, Line plot showing the APA signals of CRE contacts in condensin II-deficient mitotic cells. **f**, APA plots of CRE contacts in asynchronous and mitotic cells with or without condensin I. **g**, APA plots of CRE contacts in asynchronous and mitotic cells with or without condensin II.

### mA1 compartments and CRE contacts emerge swiftly in the unperturbed post-mitotic cells

During mitotic exit, condensin is gradually unloaded from chromosomes. We next wondered whether structures that were rapidly formed in the condensin-deficient mitotic chromosomes (e.g. mA1 compartments and CRE contacts) also emerge quickly during mitotic exit. Therefore, we assessed the kinetics of euchromatin and heterochromatin compartmentalization in unperturbed cells traversing from mitosis into G1 using our previous Hi-C datasets ^2^. We found that mA1 compartments regained homotypic interactions promptly in ana/telophase cells whereas mB4 was formed more slowly (Extended Data Fig. 17a, b). Similar to what we observed in condensin-deficient mitotic cells, post-mitotic re-formation of the mA2 compartment was slower than that of mA1 but faster than mB4 (Extended Data Fig. 17b). Separation of mA1 from mB4 was less efficient during mitotic exit (Extended Data Fig. 17a-c), potentially due to residual condensin activities in ana/telophase. Thus, euchromatin re-compartmentalizes significantly faster than constitutive heterochromatin after mitosis.

To assess whether the mitotic CRE contacts were quickly assembled after mitosis, we ranked the 6,825 CRE contacts on the basis of their strength during mitosis (Extended Data Fig. 17d). We observed a remarkably efficient enrichment of focal interactions in ana/telophase for the top 10% CRE interactions (Extended Data Fig. 17e, f). By contrast, the bottom 10% contacts displayed no appreciable signals in ana/telophase and were reformed significantly slower (Extended Data Fig. 17e, f). We previously demonstrated that reconnection of CREs after mitosis can be influenced by structural (i.e. CTCF/cohesin driven) loops. We therefore repeated our analysis on the post-mitotic Hi-C datasets of CTCF-deficient cells, which were devoid of structural loops ^3^. Notably, we observed a similar trend of fast intensification of the top 10% CRE contacts, preceding the middle and bottom 10% (Extended Data Fig. 17g).

Taken together, strong mitotic CRE contacts may be independent of transcription and transcription factor binding and are resumed remarkably fast during mitotic exit, providing an explanation for our prior observation that CRE loops are rapidly established after mitosis ^2^. By contrast, weak mitotic CRE interactions reform at a lower rate in new born cells and potentially require post-mitotic recruitment of chromatin binding partners (e.g. transcription factors).

### HP1 proteins are dispensable for the re-establishment of chromatin compartmentalization during the mitosis-to-G1 phase transition

Since most chromatin associated proteins are displaced from mitotic chromosomes, we reasoned that interrogation of the architecture of the condensin-depleted mitotic chromosomes may also inform us about the roles of selected chromatin binding proteins in genome folding. For example, HP1 proteins have been suggested to partition constitutive heterochromatin *in vitro* and *in vivo* ^23-27^. Consistent with these reports, ChIP-seq revealed significant enrichment of HP1 proteins in the constitutive heterochromatin (mB4) regions (Fig. 6a, b). However, during mitosis, all three members of the HP1 family (HP1α, HP1β and HP1γ) were evicted from the chromosome (Fig. 6a-e). Surprisingly, we observed self-aggregation of mB4 in condensin-depleted mitotic chromosomes despite lacking HP1 protein association, raising the question to what extent, if at all, HP1 proteins participate in heterochromatin compartmentalization in general. To address this question, we examined whether HP1 proteins participate in the *de-novo* establishment of compartments during G1 phase entry. We generated a compound homozygous G1E-ER4 cell line carrying mAID tag at the *Cbx5* locus (encoding HP1α) and a FKBP^F36V^ tag at the *Cbx1* locus (encoding HP1β) (Extended Data Fig. 18a). Treatment with 5-Ph-IAA or dTag13 separately or together degraded HP1α, HP1β or both proteins within 4 hours (Extended Data Fig. 18b). Using the nocodazole cell cycle arrest/release approach and FACS sorting, we obtained early- and late-G1 phase cells with four distinct configurations of HP1 proteins: (1) control, (2) HP1α(−), (3) HP1β(−) and (4) HP1α(−);HP1β(−*) (*Extended Data Fig. 18c-e). Depletion of HP1α or/and HP1β caused no noticeable defects in mitotic progression (Extended Data Fig. 18f). *In-situ* Hi-C yielded reproducible results between biological replicates (Extended Data Fig. 19a). Consistent with our previous report ^2^, chromatin compartments emerged in early-G1 phase cells followed by subsequent intensification and expansion from diagonal-proximal regions throughout entire chromosome (Extended Data Fig. 19b). Importantly, visual examination of individual chromosomes as well as genome-wide saddle plots failed to reveal noticeable defects in chromatin compartmentalization in cells lacking HP1α, HP1β or both (Extended Data Fig. 19b, c). The rate of compartment expansion was also unaffected by HP1 loss as illustrated by *P(s)* curves (Extended Data Fig. 19d). Apart from compartments, post-mitotic reformation of structural loops and CRE contacts was also unperturbed in the absence of HP1 proteins (Extended Data Fig. 19e, f).

**Figure 6.**
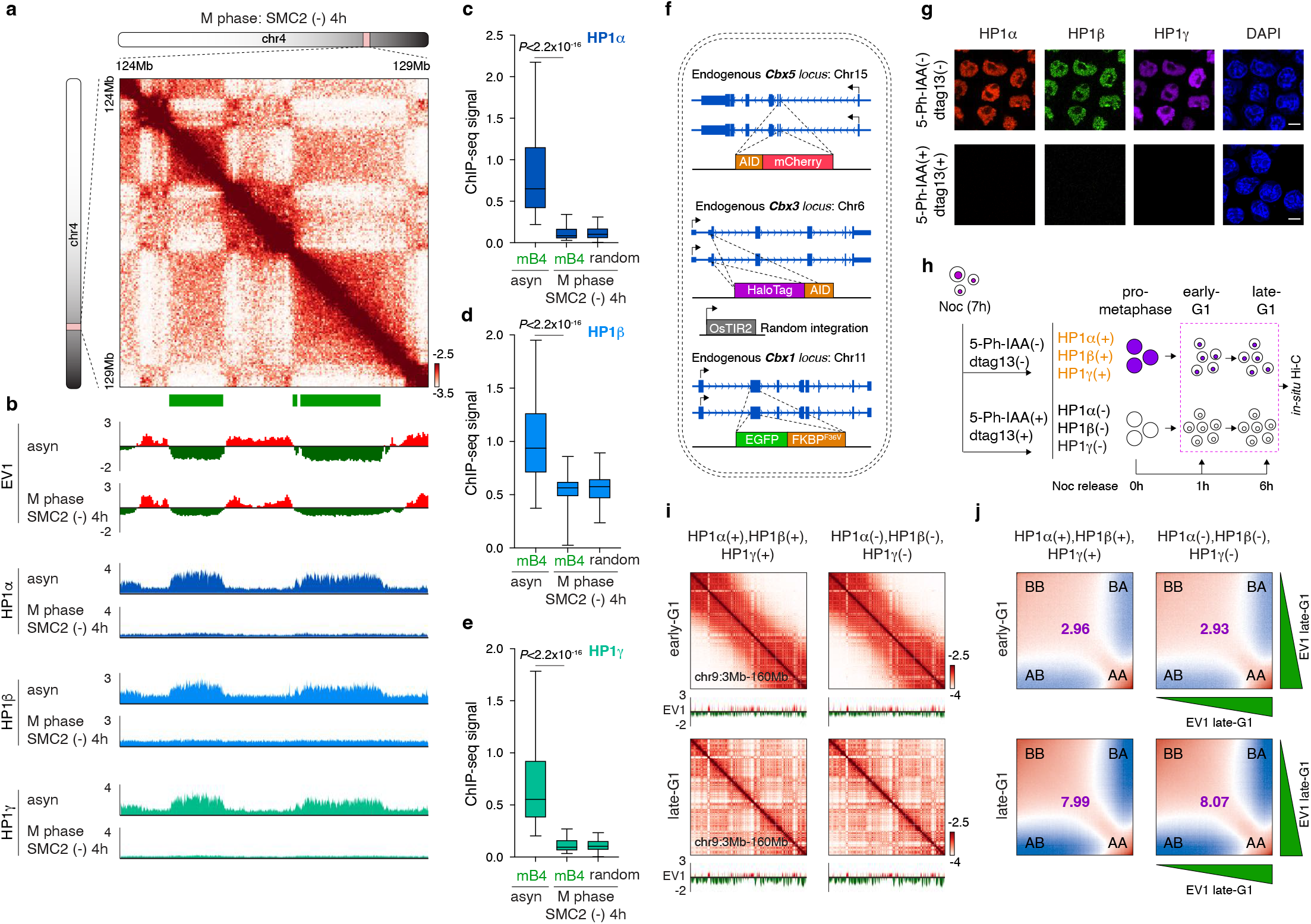
HP1 proteins are dispensable for chromatin re-compartmentalization during mitosis-to-G1 phase transition. **a**, Representative KR balanced Hi-C contact map (chr4:124-129Mb) showing mB4 homotypic interactions. Bin size: 25kb. mB4 compartments are indicated by green bars. **b**, Representative tracks corresponding to the genomic regions in (**a**), showing EV1 values and the ChIP-seq profiles of HP1α, HP1β and HP1γ in asynchronous as well as condensin-deficient (4h) mitotic cells. **c-e**, Box-plots showing the enrichment of indicated HP1 protein in asynchronous and condensin-deficient (4h) mitotic cells for mB4 (n=2,866) and random genomic regions (n=5,000). For all box plots, central lines denote medians; box limits denote 25th–75th percentile; whiskers denote 5th– 95th percentile. *P* values were calculated using a two-sided paired Wilcoxon signed-rank test. **f**, Schematic showing the genome editing strategy to generate the triple inducible degradation cell line targeting HP1α, HP1β and HP1γ simultaneously. **g**, Representative images showing the effect of acute protein degradation for HP1α, HP1β and HP1γ (Halo-646). Nuclei was marked by DAPI staining. Scale bar: 5*μ*m. Two independent experiments were performed. **h**, Schematic illustration showing the strategy of nocodazole based arrest/release in conjunction with 5-Ph-IAA and dTag13 treatment. Early and late-G1 phase cells with or without all HP1 proteins were collected. **i**, KR balanced Hi-C contact matrices (chr9:3-160Mb) showing chromatin the compartmentalization in early- and late-G1 cells. Samples with or without HP1 proteins were shown. Bin size: 100kb. Tracks of corresponding EV1 values were coupled. **j**, Saddle-plots showing the progressive compartmentalization of chromatin from early-G1 to late-G1. Samples with or without HP1 proteins were shown. Compartmental strength for sample were shown for each plot.

HP1γ has been found in euchromatin and thus seemed a less likely candidate for compartment mB4 formation ^28^. However, to rule out the possibility that HP1γ may compensate for the loss of HP1α and HP1β, we generated a cell line in which all three HP1 genes were fused homozygously with inducible degrons (Fig. 6f, g). Early and late-G1 phase cells with or without HP1 proteins were collected as described above (Fig. 6h). Importantly, co-depletion of all HP1 proteins did not alter post-mitotic reformation of compartments, structural loops and CRE contacts (Fig. 6i, j; Extended Data Fig. 20a-c). Taken together, our data demonstrate that all HP1 proteins are dispensable for the compartmentalization of mitotic chromatin as well as for the *de-novo* establishment of Hi-C detectable chromatin structures in newborn nuclei.

## DISCUSSION

It has been proposed that mammalian chromatin compartmentalization cannot be sufficiently described in terms of the original binary separation of A/B compartments ^1,17,29^. Several recent studies performed integrative data analysis using a variety of model systems and explored the relationship between compartmental interactions and chromatin-associated features ^1,29,30^. These efforts yielded multiple types of active and repressive compartments and reached the general conclusion that compartmentalization is highly correlated with chromatin-associated features, especially histone modification profiles. Yet, questions persist regarding the causality between histone modification, “reader proteins”, and gene expression in relation to chromatin compartmentalization. Most prior studies on the subject were performed in interphase cells in which compartmentalization may be influenced by multiple forces, including chromatin binding factors, active transcription, association between chromatin and the nuclear lamina, chromatin-nuclear body interactions, and cohesin mediated loop extrusion. In aggregate, these positive and negative forces may drive or mask some of the fundamental aspects of chromatin compartmentalization. Hence, examination of condensin-depleted mitotic chromosome architecture affords a new angle for the study of chromatin folding principles. Specifically, depletion of condensin in mitotic cells may provide insights into its role in perturbing or promoting chromosome structures and unveil new forces that shape chromosomal compartments. Compared to interphase chromatin, the condensin-deficient mitotic chromosome architecture differs in several key aspects. First, TADs and structural loops remained undetectable in mitotic chromosomes after condensin loss. Second, the condensin-depleted mitotic chromosomes formed two distinct types of euchromatin compartments (mA1 and mA2). The mA1 compartment exhibited an unexpected gain of self-aggregation during mitosis upon condensin loss, whereas contacts among mA2 were weak. Notably, a binary classification of mA1 and mA2 for mitotic euchromatin compartments is only a rough categorization for a series of bins with a continuum of biochemical or biophysical properties. Within the active genome, additional interaction preferences may exist.

Why mA1 and mA2 compartments behave differently is an open question. Gene expression remained completely silent in condensin-deficient mitotic cells, ruling out transcription as the driving forces of mA1 or mA2 compartmentalization. While we observed a higher enrichment of active histone marks in mA1 than mA2 in interphase cells, we did not see a fundamental difference of their epigenetic signatures. In the condensin-deficient mitotic cells, several active histone marks were attenuated, consistent with a prior study showing global histone deacetylation during mitosis^31^. Interestingly, we found a more substantial reduction seen in the mA2 compartment compared to mA1 for H3K27ac (Extended Data Fig. 14d), indicating that loss of H3K27ac may account for the lower mA2 self-aggregation during mitosis. Therefore, the distinct interaction patterns of mA1 and mA2 may originate from mitotic specific reconfiguration of epigenetic profiles. Although mA1 compartment is less sensitive to H3K27ac reduction, it is not completely immune to H3K27ac depletion. Chemically induced H3K27ac removal disrupted a quarter of mA1 compartments (mA1-D) in the condensin-deficient mitotic cells. This percentage is probably an underestimate given that the unchanged mA1-U and enhanced mA1-I compartments displayed insufficient H3K27ac depletion after A485 treatment (Fig. 4k), Together, using the condensin-deficient mitotic chromosomes as a platform, we identified two sub-types of euchromatin with distinct interaction preferences and sensitivities to H3K27ac loss. One caveat of our experiment is that A485 treatment may also reduce acetylation at other histone residuals which may play a functional role.

Whether the histone mark H3K27ac directly influence euchromatin compartmentalization is unclear. BET protein inhibition also reduced mA1-D compartmentalization, but to a lesser extent compared to removal of H3K27ac. Thus, BET proteins may play a role in the H3K27ac mediated mA1 compartmentalization. However, the compartmental strength of mA1-D after A485 treatment is still higher than that of mA2 in the control samples, indicating the existence of additional forces other than H3K27ac that drive mA1 compartmentalization. These data are consistent with the low sensitivity of mA1 compartmentalization to H3K27ac loss. We do not rule out a potential role of additional active histone marks. It is also worth mentioning that RNA has been previously shown to mediate chromatin compartmentalization in interphase cells ^32,33^. Moreover, a variety of RNA species remain associated with chromosomes during mitosis ^34-36^. Thus, chromosome-associated RNAs may contribute to mitotic compartmentalization.

During the mitosis-to-G1 phase transition, condensin is gradually unloaded from the chromatin, mirroring the process of cohesin degradation in mitotic cells. The tendency of mitotic chromosomes to promptly form mA1 compartments and CRE contacts upon condensin loss raised a possibility that these structures are also rapidly formed when cells exit from mitosis. Indeed, we confirmed the fast formation of mA1 compartments as well as strong mitotic CREs in parental cells traversing from mitosis into G1. Our analyses uncovered intrinsic properties of mitotic chromosomes other than post-mitotic recruitment of chromatin binding partners as the driving force of the efficient re-establishment of euchromatin compartmentalization and CRE contacts after mitosis.

Interestingly, the global levels of many histone modification marks are maintained during mitosis as measured by mass spectrometry ^20,37^. Using calibrated ChIP-seq, we found that mitotic chromosomes displayed similar levels of the facultative heterochromatin (H3K27me3) compared to asynchronous controls (Extended Data Fig. 14b). Moreover, facultative heterochromatin (mB1) and constitutive heterochromatin (mB4) can co-compartmentalize around centromeres and telomeres in condensin-depleted mitotic chromosomes but not in interphase chromatin. We speculate that in interphase, additional mechanisms are at play to separate mB4 from mB1, such as binding to “reader” proteins of the heterochromatic marks, or distinct subnuclear distribution of chromatin, such as the association with LADs or nucleolus associated domains (NADs) ^38^.

Several reports suggested a role for HP1 proteins in forming heterochromatin compartments. HP1 proteins have been proposed to be able to form condensates with DNA *in-vitro* and form heterochromatic nuclear foci *in-vivo ^26,39^*. HP1 proteins may aid in the maintenance of the heterochromatin mark H3K9me3 ^40^, and loss of H3K9me3 can eliminate heterochromatin compartmental interactions ^29^. Artificial placement of H3K9me3 by dCas9-KRAB can trigger new compartment formation ^41^. Finally, HP1 depletion was shown to impair chromatin compartments in *Drosophila* embryos ^24^. Considering these studies, it was surprising to find mB4-to-mB4 interactions in condensin-depleted mitotic chromosomes that lack HP1 proteins. This observation motivated us to inquire whether HP1 proteins directly facilitate chromatin compartmentalization in general. Acute depletion of HP1α, HP1β and HP1γ during the mitosis-to-G1 phase transition failed to slow or diminish the re-establishment of any hallmark chromatin structures, including compartments. Our data demonstrate that HP1 proteins are dispenable for proper genome re-folding after mitosis, and they suggest additional mechanism that mediate heterochromatin compartmentalization. These findings are consistent with a previous report that the compaction of heterochromatic foci is independent of HP1 proteins ^39^.

In summary, by leveraging the auxin inducible acute degradation system, we created mitotic chromosomes without condensin-mediated constrains while maintaining the mitotic cellular context. With this strategy, we uncovered detailed patterns of chromatin compartmentalization and novel insights that are unattainable when studying interphase cells.

## ACKNOWLEDGEMENTS

We thank Dr. Leonid Mirny, Dr. Marit Vermunt, Dr. Xiaohua Shen and members of the Zhang and Blobel labs for helpful discussions. We are indebted to Mengyuan Li, Zhiqing Huang and Dechun cheng from the Flow-cytometry Core of Shenzhen Bay Laboratory for technical support on cell sorting. We thank Shixian Huang and Mei Yu from the Bioimaging Core of Shenzhen Bay Laboratory for imaging support. This work was supported by the National Science Foundation of China grant 32100422 to H.Z. and NIH grant U01DK127405 and R01DK058044 to G.A.B.

## AUTHOR CONTRIBUTIONS

G.A.B. and H.Z.* conceived the study and designed experiments. H.Z. and Y.L. created the inducible acute degradation cell line used in this study. H.Z. performed sample preparation, FACS sorting, *in-situ* Hi-C, ChIP-seq and microscopic experiment with help from E.L., L.S., D.J., B.W., M.W., L.Z., F.S., J.L. and S.M. C.K., B.G. and R.H. contributed to sequencing of *in-situ* Hi-C materials. Data analysis were performed by H.Z.* with help from F.L. H.Z.* and G.A.B. wrote the manuscript with inputs from all authors.

## CONTACT FOR REAGENT AND RESOURCE SHARING

Further information and requests for resources and reagents should be directed the lead contact, Gerd A. Blobel and Haoyue Zhang

## METHODS

### Cell culture

The parental G1E-ER4 cell line was a gift from the laboratory of Mitchell Weiss and cultured as previously described. Briefly, G1E-ER4 cells were cultured in IMDM medium supplemented with 15% (v/v) FBS, 2% (v/v) penicillin-streptomycin, 50ng/ml murine stem cell factor, 7.5ng/ml Epogen and 1/100,000 (v/v) 1-Thioglycerol. Cells were maintained below a density of 1 million/ml. The parental G1E-ER4 line and all derivative sublines were periodically tested for mycoplasma. All lines were validated to be mycoplasma negative. All cell lines were cultured in a humidified incubator at 37°C, with 5% CO2.

### CRISPR/Cas9 mediated genome editing

All the inducible acute degradation cell lines listed in this work were generated based on the G1E-ER4 murine erythroblasts using CRISPR/Cas9 based genome editing technology. The edited cell lines include: (1). G1E-ER4:SMC2-mAID-mCherry; (2). G1E-ER4:NCAPH-mAID-HaloTag; (3). G1E-ER4:NCAPH2-mAID-mCherry; (4). G1E-ER4:mCherry-mAID-HP1α/GFP- FKBP^F36V^-HP1β and (5). G1E-ER4:mCherry-mAID-HP1α/GFP-FKBP^F36V^-HP1β*/*HaloTag-mAID-HP1γ. Below is the detailed strategy to generate the G1E-ER4:SMC2-mAID-mCherry cell line. The homology directed repair template was constructed as follows: (1) a 1kb left homology arm fragment that covers part of intron24 and exon 25 (before stop codon) of the *Smc2* gene was PCR amplified from the mouse genomic DNA; (2) A 909bp fragment after the stop codon of the *Smc2* gene was amplified as the right homology arm; (3) a minimal functional mAID tag (aa: 1- 44) and the mCherry coding sequence were separately PCR amplified. These four fragments were assembled together through Gibson assembly (NEB, Cat#E2611L) and cloned into the pCMV-YFP-C1 plasmid to serve as the repair template. A single-guide RNA (sg-RNA) targeting the 3’ end of the *Smc2* gene was designed through the Benchling online tool. Sense and anti-sense sgRNA oligos were annealed together and cloned into the pX458 plasmid which contained a U6 sgRNA expressing cassette and encoded Cas9-P2A-GFP. To enable genome editing, 18*μ*g of sgRNA containing pX458 plasmids and 18*μ*g of the repair template were co-transfected into 5 million parental G1E-ER4 cell lines with a Amaxa 2b Nucleofector with the transfection program G-016. Transfected cells were cultured for 24 hours in antibiotic free culture medium to reduce cell death. Transfected single cells with high mCherry signals were FACS sorted into 96 well plates through a Beckman Coulter Moflo Astrios sorter and further cultured for 7 days until single cell clones were visible. Successfully edited homozygous clones were confirmed by genotyping with primers flanking the left and right homology arm.

Similar strategies were adopted to generate all the other cell lines. To obtain cells with successful insertion of HaloTag, transfected cells were maintained in antibiotics free medium for 24h. Cells were then stained with 5nM Janelia Fluor HaloTag Ligand 646 (GA1120, Promega) for 30min in the incubator and washed for 3 times before single cell FACS sorting. Sequences of all oligos described above to construct repair templates, sgRNAs and genotyping primers to validate edited clones are listed in Supplementary table 4.

### Retroviral infection of murine cells

To enable rapid degradation of mAID tagged endogenous proteins, OsTiR or OsTIR2 were delivered into cells through retroviral infection. Briefly, coding sequence of OsTIR or OsTIR2 was cloned into MigR1 retroviral vector that contained either GFP tag. For virus packaging, 15*μ*g of MigR1-OsTIR-GFP vectors were co-transfected with 15*μ*g of pCL-Eco packaging plasmids into HEK293T cells at 90% confluence through PEI (Polysciences Cat#23966). Virus were collected at 48h and 72h respectively and filtered through 0.45*μ*M filters to remove cell debris. For retroviral infection, 3 million target cells were plated into 6-well plates with 1ml of culture medium and 1ml of viral supernatant, supplemented with 8 mg/ml polybrene and 10mM HEPES buffer (Gibco Cat#15630-106). Cells were spin-infected at 3000 rpm for 1.5h at room temperature. Cells were then washed with PBS and re-suspended in fresh culture medium. 48h later, infected cells were enriched through FACS sorting based on GFP signals by a Beckman Coulter Moflo Astrios sorter.

### Cell cycle analysis of SMC2, NCAPH or NCAPH2 depleted cells

To determine the effect of SMC2 depletion on cell cycle progression, one million G1E- ER4:SMC2-mAID-mCherry cells were treated with 1mM of auxin for 0h, 0.5h, 1h, 2h, 4h and 8h respectively. After drug treatment, cells were fixed with 1% formaldehyde (Sigma Cat#F8775) in 1× PBS for 10min at room temperature and then stained with DAPI (4*μ*g/ml, Sangon Biotech Cat#A606584-0010) in staining buffer (1× PBS containing 2mM EDTA and 0.1% Triton-X 100) for 5min. Cell cycle distribution and depletion of SMC2-mAID-mCherry signal were assessed by a BD LSRFortessa^TM^ cell analyzer. Parent G1E-ER4 cells were used as control. Similar strategy was applied to G1E-ER4:NCAPH-mAID-HaloTag and G1E-ER4:NCAPH2-mAID-HaloTag cells.

### Cell growth curve

To examine the influence of SMC2 depletion on cell growth, ∼10,000 G1E-ER4:SMC2-mAID-mCherry cells were seeded in 6 well plates in 2ml of fresh culture medium. Cells were then treated with 1mM of auxin to degrade SMC2. Cells without any treatment were kept as control. Cell number of each treatment condition was counted every 24 hours for a total of 72-hour duration. Three independent biological replicates were performed.

### Fluorescent confocal imaging

To examine the cellular localization of SMC2-mAID-mCherry fusion protein, asynchronously growing G1E-ER4:SMC2-mAID-mCherry cells were seeded into glass-bottom 384-well plate and imaged with a Dragonfly CR-DFLY-202-2540 spinning disc confocal microscope equipped with 100× objective lense (NA1.45). To examine the cellular localization of NCAPH-mAID-HaloTag protein, asynchronously growing G1E-ER4:NCAPH-mAID-HaloTag cells were stained with 5nM Janelia Fluor HaloTag Ligand 646 for 30min in the incubator, washed three times before imaging.

To examine mitotic chromosome morphology after SMC2 depletion, about 0.5 to 1 of million G1E-ER4:SMC2-mAID-mCherry cells were arrested to prometaphase through nocodazole enrichment. Toward the end of nocodazole treatment, cells were treated with auxin for 0.5h, 1h, 2h, 4h and 8h respectively. After treatment, cells were fixed with 1% formaldehyde for 10min at room temperature. After fixation, cells were stained with pMPM2 antibody (05-368, Milipore; 0.1*μ*l per 10 million cells) to identify mitotic populations. Cells were then re-suspended in 100*μ*l of staining buffer (0.1% Triton-X100 and 2mM EDTA diluted in PBS) containing 1µg/ml DAPI. Cells were then seeded into a glass-bottom 384-well plate and imaged with an Olympus Spin SR super-resolution spinning disc confocal microscope equipped with 100× objective lense (NA1.5). Similar strategies were applied to G1E-ER4:NCAPH-mAID-HaloTag and G1E-ER4:NCAPH2- mAID-HaloTag cells without pMPM2 staining.

### Mitotic chromosome spread assay

To examine the impact of NCAPH2 depletion on sister chromatid resolution, cells were treated with nocodazole for 12h to enrich prometaphase population. Meanwhile, cells were either treated with or without 5-Ph-IAA for 4 hours. After treatment, cells were pelleted and re-suspended in 0.075M KCl and incubated at 37℃ for 15min. Cells were then pelleted at room temperature at 200g for 5min and re-suspended in 5 ml of fresh Carnoy’s Fixative (3:1 ratio of methanol: acetic acid). Fixation was performed twice. Cells were pelleted and re-suspended in 200*μ*l of fresh Carnoy’s Fixative. Drop the cells from 0.5 meter above onto a slide tilted with a 45° angle. The slides chromosome spreads were immediately dried over the flames of an alcohol burner and stained Giemsa solution (Solarbio, Cat#G1015) for 8min, washed by running water and air dried. The slides were then mounted for long term storage and imaging.

### Profiling of LADs

To profile lamina associated domains (LADs) of G1E-ER4 cells, we performed CUT&TAG on asynchronously growing G1E-ER4 cells using lamin B1 antibody (ab16048, abcam) according to the manufacture’s protocol (Vazyme Hyperactive® Universal CUT&Tag Assay Kit for Illumina).

### Drug treatment, sample preparation and cell sorting. For *In-situ* Hi-C experiments

1. To assess the dynamic re-configuration of chromatin architecture after condensin removal during mitosis, G1E-ER4:SMC2-mAID-mCherry cells expressing OsTIR were first arrested to prometaphase with nocodazole (200ng/ml) for 12 hours or 15 hours (for 8h release condition). Toward the end of mitotic arrest, cells were treated with auxin (1mM) for 0 hour (SMC2+), 0.5 hours, 1 hour, 4 hours or 8 hours respectively. Mitotic cells were collected through FACS sorting of pMPM2 positive populations.
2. To assess condensin-deficient mitotic chromatin architecture without nocodazole synchronization, asynchronous G1E-ER4:SMC2-mAID-mCherry cells expressing OsTIR were treated with or without auxin (1mM) for 4 hours. Mitotic cells from asynchronous population were collected through FACS sorting based on pMPM2 staining.
3. To assess the dynamic re-configuration of mitotic chromosome architecture after loss of condensin I or condensin II, G1E-ER4:NCAPH-mAID-HaloTag or G1E-ER4:NCAPH2- mAID-HaloTag cells expressing OsTIR2 were arrested to prometaphase with nocodazole treatment as described above. Cells were treated with 5-Ph-IAA (1*μ*M) for 0 hour, 0.5 hours, 1 hour, 4 hours or 8 hours respectively. Mitotic cells were collected through FACS sorting of pMPM2 positive populations.
4. To assess the role of histone modification H3K27ac and BET protein occupancy on mitotic specific compartmentalization, G1E-ER4:SMC2-mAID-mCherry cells expressing OsTIR were pre-treated with the 20*μ*M of the p300 inhibitor A485 (S8740, Selleck) or 250nM JQ1 (HY-13030, MCE) followed by nocodazole mediated mitotic arrest for 12 hours. Toward the end of mitotic arrest, cells were treated with auxin (1mM) for 4 hours to degrade SMC2. DMSO pre-treated cells were processed in parallel as control. Mitotic cells were collected through FACS sorting of pMPM2 positive populations.
5. To assess the dynamic re-configuration of chromatin architecture through mitosis-to-G1 phase transition with different HP1 protein configurations, G1E-ER4:mCherry-mAID- HP1α;GFP-FKBP^F36V^-HP1β cells were arrested to prometaphase through nocodazole treatment for 8h. During the last 4 hours of nocodazole treatment, cells were treated with one of the following combinations of drugs: (i) 5-Ph-IAA(-)/dTag13(-); (ii) 5-Ph-IAA(+)/dTag13(-); (iii) 5-Ph-IAA(-)/dTag13(+) and (iv) 5-Ph-IAA(+)/dTag13(+). The concentrations of 5-Ph-IAA and dTag13 were 1*μ*M and 200nM respectively. Cells were then allowed to be released from nocodazole and proceed toward the next cell cycle. After 1h and 6h of nocodazole release, early-G1 and late-G1 cells under different HP1 protein configurations were collected. Note, that 5-Ph-IAA or/and dTag13 treatment was maintained during the course of nocodazole release.
6. To address whether triple depletion of HP1α, HP1ϕ3 and HP1γ affect post-mitotic re-compartmentalization, G1E-ER4:mCherry-mAID-HP1α/GFP-FKBP^F36V^-HP1ϕ3*/*HaloTag-mAID-HP1γ cells were arrested to prometaphase through nocodazole treatment for 8h. During the last 4 hours of nocodazole treatment, cells were treated with dTag13 (200nM) and 5-Ph-IAA (1*μ*M) to allow rapid degradation of all three HP1 proteins. Cells were then allowed to be released from nocodazole and proceed toward the next cell cycle. After 1h and 6h of nocodazole release, early-G1 and late-G1 cells were separately collected. Cells without dTag13 or 5-Ph-IAA treatment were collected as controls.

At the end of drug treatment, all the above cells were crosslinked with 2% formaldehyde for 10min at room temperature. Crosslinking was stopped by glycine (Sangon Biotech Cat#A100167) with a final concentration of 1M. Cells were then permeablized by 0.1% Triton X- 100 for 5min. For mitotic cell purification, the following steps were taken: after cell membrane penetration, all cells were stained with anti-pMPM2 antibody (Millipore Cat#05-368, 0.2µl/10million cells) for 50min at room temperature, followed by APC-conjugated F(ab’)2-goat anti-mouse secondary antibody (Thermo Fisher Scientific, Cat#17-4010-82, 2µl/10million cells) for 45min at room temperature. After staining, cells were re-suspended in 1× FACS buffer (1× PBS containing 2mM EDTA and 2% FBS) at a density of 100 million cells per ml and subject to FACS purification. Cells were sorted for mitotic population based on pMPM2 signal (+) and DAPI signals (4N). For newborn G1 cells purification, the following steps were carried out: cells were re-suspended in 1× FACS buffer (1× PBS containing 2mM EDTA and 2% FBS) at a density of 100 million cells per ml and subject to FACS purification. Cells were purified for G1 phase cells based on DAPI (2N), mCherry and GFP signals. FACS purification were performed via a BD FACS AriaIII cell sorter and 200,000 cells were collected for each condition. Sorted cells were pelleted, snap-frozen through liquid nitrogen and stored at -80℃ until further use.

### For ChIP-seq experiments

To compare the level of histone modifications between interphase and SMC2 depleted mitotic cells, G1E-ER4:SMC2-mAID-mCherry cells were arrested to mitosis with nocodazole (200ng/ml) for 12 hours. Toward the end of mitotic arrest, cells were treated with 1mM auxin for 4 hours to degrade SMC2. At the end of drug treatment, cells were harvested in a similar fashion compared to the above samples for Hi-C experiments except that cells were crosslinked with 1% formaldehyde. Protease inhibitor (1:500 Sigma, P8340) and PMSF (Sangon Biotech Cat#A100754, 1:100) were added to all buffers during the entire sample preparation procedure. Mitotic and cells were purified via a BD FACS AriaIII cell sorter based on pMPM2 staining and DAPI signals (4N). Untreated asynchronous cells were processed in parallel and purified through FACS sorter as control. Sorted cells were pelleted, snap-frozen through liquid nitrogen and stored at -80℃ until further use.

### In-situ Hi-C

We optimized our *In-situ* Hi-C protocol to generate high quality Hi-C libraries from relatively small amount of input material. Briefly, FACS purified cells (0.1-0.2 million) were lysed in 1ml cold Cell Lysis Buffer (10mM Tris pH 8, 10mM NaCl, 0.2% NP-40/Igepal) for 10min on ice. Nuclei were pelleted at 4℃ and washed once with cold 1.4 × NEB buffer 3.1 (NEB Cat#B7203S). Nuclei were then re-suspended in 25*μ*l of 1.4 × NEB buffer 3.1 and treated with 0.1% SDS for 10min at 65℃. Nuclei were then treated with 1% Triton X-100 to quench SDS for 1h at 37℃. Chromatin was digested with 25U DpnII restriction enzyme (NEB, Cat#R0543M) at 37℃ overnight with shaking (950rpm on a thermal mixer). After overnight digestion, an additional stroke of 25U DpnII was added into the system to further digest the genome for 4 hours at 37℃. After the 2^nd^ stroke of digestion, nuclei were incubated at 65℃ for 20min to inactivate DpnII. Digested chromatin fragments were then cooled down to room temperature and blunted with dCTP, dGTP, dTTP (Diamond Cat#B110049, B110050, B110051) and Biotin-14-dATP (active motif, Catalog No: 14138) using 6.6U DNA Polymerase I, Large (Klenow) fragment (NEB, Cat#M0210). Chromatin was then ligated *in-situ* with 2000U T4 DNA ligase (NEB, Cat#M0202M) for 4 hours at 16℃ followed by further incubation for 2 hours at room temperature. Chromatin was then reverse-crosslinked overnight at 65℃ in the presence of 1% SDS and proteinase K (G-clone Cat#EZ0970). After reverse-crosslinking, DNase-free RNase A (Vazyme Cat#DE111-01) was added to the system for 30min at 37℃ to remove RNA. DNA was then extracted by phenol-chloroform extraction, precipitated, and dissolved in 130*μ*l of nuclease free water. DNA was sonicated to 200-300bp fragments with a QSonica Q800R3 sonicator (40% amplitude, 15s ON and 15s OFF, 13min sonication) and purified with VAHTS DNA Clean Beads (Vazyme Cat#N411). Biotin-labeled DNA fragments was enriched by incubation with 100µl Dynabeads MyOne Streptavidin C1 beads (Thermal Fisher Scientific, Cat#65002) at room temperature for 15min. Streptavidin C1 beads were then washed twice with 1 × B&W buffer (5mM Tris-HCl, pH 7.5; 0.5mM EDTA; 1M NaCl). Beads were re-suspended in 30*μ*l of 1 × NEB Buffer2 (NEB Cat#7002S) containing 9U of T4 DNA polymerase (NEB, Cat#M0203S) and 40*μ*M dATG/dGTP and incubated at 20℃ for 4 hours to remove unligated fragments. Beads were then washed twice with 1 × B&W buffer and subject to library construction. DNA end repair, dA-tailing and adaptor ligation were constructed on beads, using the AHTS Universal DNA Library Prep Kit for Illumina or MGI (Vazyme, Cat# ND627-01, NDM607-02) based on the manufacture’s protocol. To eluted DNA from streptavidin beads, the library mixture was treated with 0.1% SDS and incubated at 98℃ for 10min. Released DNA was collected from the supernatant and purified with VAHTS DNA Clean Beads. Finally, the library was amplified for 9 cycles on a thermal cycler, using the VAHTS® HiFi Amplification Mix. PCR products were then purified with VAHTS DNA Clean Beads and sequenced on an Illumina NextSeq500 or MGI DNBSEQ-T7 sequencing platform.

### ChIP-seq

Chromatin immunoprecipitation (ChIP) was performed as previously described, using antibodies against H3K27ac (Abcam Cat#ab4729, 3*μ*l/IP), H3K4me1 (Abcam Cat#ab176877, 3*μ*l/IP), H3K27me3 (Active motif Cat#: 39055, 3*μ*l/IP), H3K36me3 (Abcam Cat#ab9050, 3*μ*l/IP), RNA PolII (CST Cat#14958s, 3*μ*l/IP), Rad21(Abcam, Cat#ab992, 3*μ*l/IP), HP1α (Abcam Cat#ab109028, 5*μ*l/IP), HP1β(CST Cat#8676, 5*μ*l/IP) and HP1γ (Abcam Cat#ab217999, 5*μ*l/IP) respectively. For histone modifications, half a million cells were used per IP. For RNA PolII, Rad21 and HP1 proteins, 1 million cells were used. FACS purified cells were re-suspended in 1ml pre-cooled Cell Lysis Buffer supplemented with 1:500 protease inhibitor and 1:100 PMSF for 20min on ice. To control for global change of histone modification that cannot be detected by ChIP-seq, HEK293T cells were used as spike-in controls and added into FACS purified mitotic and asynchronous cells so that both samples contain comparable level of human genome (∼30%). Nuclei were then re-suspended in 1ml Nuclear Lysis Buffer (50mM Tris pH 8, 10mM EDTA, 1% SDS, fresh supplemented with PI and PMSF) for 20min on ice. Chromatin was then fragmented to 200-300bp by sonication (Qsonica Q800R3, 80% amplitude, 20s ON, 40s OFF) for 17min. Samples were centrifuged at 15000g for 10min at 4℃ to remove cell debris. Chromatin in the supernatant was collected and diluted with 4 volumes of IP dilution buffer (20mM Tris pH 8.0, 2mM EDTA, 150mM NaCl, 1% Triton X-100, 0.01% SDS) and then supplemented with protease inhibitor, PMSF and 50*μ*l of A/G agarose beads (Santa Cruz Cat#sc-2003). Preclear of chromatin was performed at 4℃ for 8h with rotation. Precleared chromatin was then incubated with 35µl A/G agarose beads (A:G = 1:1) pre-bound with ChIP antibody at 4℃ overnight. Beads were washed once with IP wash buffer I (20mM Tris pH 8, 2mM EDTA, 50mMNaCl, 1% Triton X- 100, 0.1% SDS), twice with high salt buffer (20mMTris pH 8, 2mM EDTA, 500mMNaCl, 1% Triton X-100, 0.01% SDS), once with IP wash buffer II (10mMTris pH 8, 1mM EDTA, 0.25 M LiCl, 1% NP-40/Igepal, 1% sodium deoxycholate) and twice with TE buffer (10mM Tris pH 8, 1mM EDTA pH 8). All washing steps were performed on ice. Beads were moved to room temperature and DNA was eluted in 200µl fresh Elution Buffer (100mM NaHCO3, 1%SDS). ChIP and input chromatin were reverse crosslinked at 65℃ overnight in the prescence of proteinase K. IP and input samples were supplemented with 10ul of 3M sodium acetate (pH 5.2), and DNA was purified with QIAquick PCR purification kit (QIAGEN Cat#28106). ChIP-seq libraries were constructed using the AHTS Universal DNA Library Prep Kit for MGI (Vazyme, Cat#NDM607-02) according to the manufacture’s protocol. ChIP libraries were sequenced on the MGI DNBSEQ- T7 sequencing platform.

### Quantification and data analysis

### ChIP-seq data processing

ChIP-seq reads were first aligned to human genome assembly hg19 with default parameters. Unmapped reads were then aligned to mouse genome assembly mm9. Human and mouse alignments were then both filtered out for low-quality reads (MAPC <30), PCR duplicates and reads that aligned to mitochondria random contigs and ENCODE blacklisted regions were removed. The final read counts aligned to human hg19 genome was determined by SAMtools (v1.9) and scaling factor was calculated as: 10^6^/hg19_counts. Valid reads that aligned to mouse genome was then normalized using the above scaling factor through Deeptools (v3.1.3).

### Hi-C data preprocessing

*In-situ* Hi-C data from each individual clone or biological replicates were aligned to mouse reference genome mm9 using bowite2 (global parameters: --very-sensitive -L 30 --score-min L, -0.6,-0.2 --end-to-end –reorder; local parameters: --very-sensitive -L 20 -- score-min L,-0.6,-0.2 --end-to-end --reorder). Reads were paired up and PCR duplicates were removed through the Hi-C Pro software (v3.0.0) to generate a valid interaction pair file. The output pair files were then converted to “.hic” files using the hicpro2juicebox utility. HiC files were also converted to “.cool” files using the Hicexplorer (v3.7). For replicate or clone-merged samples, similar steps were taken on reads merged from each clone or biological replicates.

### *Trans-*chromosomal interactions

The percentage of trans chromosomal interactions were acquired from HiC Pro statistic summary. The *cis-* and *trans-* Hi-C contact matrices were generated using cooler software.

### Hi-C matrix correlation measurement

The data quality of Hi-C experiments was assessed by stratum-adjusted correlation coefficient (SCC) as previously suggested by Yang et al ^42^. SCC is calculated for each individual chromosome (chr1-19) using HiCRep on the 100kb binned matrix with a maximum distance of 50Mb.

### General A/B compartment assignment and saddle plotting

General A/B compartments of interphase samples were identified through eigenvector decomposition on the Pearson’s correlation matrix of the observed/expected values of 25kb binned Knight-Ruiz (KR) balanced cis-interaction matrices using the “eigs-cis” function of Cooltools (v0.4.0). Positive and negative EV1 values of each 25kb bin were assigned to A- (active) and B- (inactive) compartments, respectively, based on gene density. To visualize the strength of compartmentalization, saddle plots were generated. Briefly, 25kb binned KR balanced cis observed/expected contact matrix was extracted from each “.hic” file using the DUMP function of juicer_tools (v 1.13.02). For interphase samples, the above matrices were transformed in a way that each column and row of bins were re-ordered based on their corresponding EV1 value in an ascending order from top to bottom and from left to right. For the time course SMC2 depletion experiments, the contact matrices of all mitotic samples (0h, 0.5h, 1h, 4h and 8h of auxin treatment) were subject to the same transformation based on the EV1 values of asynchronous control sample. In this way, bins at the top-left corner of the saddle plot were associated with B-B compartmental interactions, whereas bins at the bottom right corner of the saddle plot were associated with A-A compartmental interactions. Bins at the top-right and lower-left corners were associated with B-A or A-B compartmental interactions. The transformed maps from each individual chromosome were divided into 200 equal sections and averaged together create the genome wide saddle plots. The compartment strength of each individual chromosome was denoted as: (median (top20% AA) + median (top20% BB)) / (median (top20% AB) + median (top20% BA)).

### Simulation of interphase contamination in mitotic samples

To rule out the possibility that the checkerboard compartment structures observed in SMC2(-) mitotic cells was originated from interphase contamination, we performed a digital simulation of chromosome 4, 9 and 11 by sampling different fraction of interphase reads into mitotic samples. Briefly, 100kb binned dense formatted n×n contact matrices with unbalanced raw read counts of SMC2(+) mitotic and late-G1 samples were extracted from “.hic” files using the DUMP utility of Juicer_tools. The matrices from each sample were then normalized to a depth of 20 million reads per chromosome. To simulate 5% interphase contamination, mitotic and late-G1 matrices were multiplied by factors of 0.95 and 0.05 respectively and then combined together to generate a mixed matrix of mitotic and interphase samples. Similar procedure was performed for 10% and 20% simulation. The mixed matrices were then visualized using R software and compared to SMC2(-) mitotic samples.

### Insulation score profiling

Insulation scores were computed on 10kb binned, KR balanced contact matrices of each interphase and mitotic samples, using the “insulation” function of Cooltools (v0.4.0). The sliding window was set to be 12bin x 12bin as we previously practiced. (v3.6.1).

### Domain calling

We identified domains and associated boundaries through combining two independent algorithms. To begin with, domain boundaries were independently identified in each SMC2 deficient mitotic samples (0.5 hour, 1 hour, 4 hour and 8 hour auxin treated) as well as the asynchronous control samples, by searching for the “local-minima” of insulation scores. In detail, boundaries were identified in each sample through the “insulation” function of Cooltools with a 12bin x 12bin scanning window and threshold of 0.2. Boundaries from each sample were then merged together through Bedtools *merge* with a minimal merging distance of 50kb to create a non-redundant list of boundaries.

Domains were independently identified in all of the above mentioned samples using the “Arrowhead” function of juicertools. Domains were called at 50kb, 25kb and 10k resolutions respectively for each condition. All identified domains were collected together to generate an initial domain list. Domain calling has been shown to be highly algorithm dependent and often exhibit high false-positive results. To further remove spurious domains, we intersected our initial domain list and non-redundant boundary list acquired above. In brief, boundaries from each chromosome were randomly paired up so that the distance between each pair of boundaries were greater than 50kb and smaller than 10Mb. Each pair of boundaries construct a potential domain. To exclude false-positive boundary pairs, they were individually intersected with the initial domain list generated by “Arrowhead”. For a boundary pair to be considered as a real domain, it must overlap with a domain identified by “Arrowhead” with a wiggling window of 50kb for each boundary. Similar procedure was iterated through all SMC2(-) mitotic samples.

### Domain partition

To examine whether a domain displays punctate corner-dot signals at its summit, we intersected our domain and chromatin loop calling results. For a domain to be considered as a corner-dot positive domain, there must be at least 1 chromatin loop that fulfills the below criterion: (1) left anchor coordinate of the loop must wiggle within a predefined range of the up-stream boundary of the target domain and (2) the right anchor coordinate of the loop must wiggle within a predefined range of the down-stream boundary of the target domain. The predefined range is set as follows: for domains smaller than or equal to 200kb, the wiggle room is within [-10kb to 30kb] relative to the up-stream boundary and [-30kb to 10kb] relative to the down-stream boundary. For domains with a size between 200kb and 400kb, the wiggle room is within [- 20kb to 40kb] relative to the up-stream boundary and [-40kb to 20kb] relative to the down-stream boundary. For domains with a size larger than or equal to 400kb, the wiggle room is within [-30kb to 50kb] relative to the up-stream boundary and [-50kb to 30kb] relative to the down-stream boundary.

### Identification of mitotic specific compartments

EV1 values were computed for 25kb binned KR balanced contact matrices of asynchronous and SMC2(-) 4h mitotic samples of the G1E- ER4:SMC2-mAID-mCherry cells, using the “eigs-cis” function of Cooltools. Positive and negative EV1 values were assigned to genomic bins based on gene density. Z-scores of EV1 values for each 25kb genomic bin were computed across both replicates of the above two samples and subject to *k*-means clustering. We identified 3 clusters of genomic bins: (1) Group I bins with increased EV1 values in mitotic SMC2 depleted cells compared to interphase. These regions may represent enhanced A-to-A compartmental interactions, decreased B-to-B compartmental interactions or conversion from B to A in mitotic samples devoid of SMC2. (2) Group II bins with decreased EV1 values during mitosis. These regions may indicate a reduction of A-to-A compartmental interactions, increase of B-to-B compartmental interactions or conversion from A to B in mitosis. (3) Group III bins either showed mild EV1 change in mitosis or with inconsistent z-scores between biological replicates and thus cannot be meaningfully categorized. To dissect mitotic specific compartmental interactions, Group I and II genomic bins were further clustered based on the composition of histone modifications of each genomic bin. To perform this analysis, we computed the ChIP-seq signals of four active histone marks (H3K27ac, H3K36me3, H3K4me1 and H3K4me3), one constitutive heterochromatic histone mark (H3K9me3) and one facultative heterochromatic histone mark (H3K27me3) for each 25kb genomic bin within group I and II. We also included the Lamina-associated domain (LAD) signal generated from a CUT&Tag experiment using the anti-Lamin B1 antibody (ab16048). Histone modification and LAD signals were first normalized across the genome to enable comparisons across different ChIP or CUT&Tag experiments. Z-scores of the normalized histone modification and LAD signals were then computed and used as input to *k*-means clustering. This analysis resulted in four major types of compartments with distinct composition of histone marks and interaction properties during in SMC2 deficient mitotic cells. These domains are: (1) mA1 compartment which were enriched with active histone marks and strengthened during mitosis; (2) mA2 compartment which were enriched with active histone marks but weakened during mitosis; (3) mB4 compartment which were enriched with H3K9me3 heterochromatin histone marks and weakened during mitosis and (4) mB1 compartment with the repressive mark H3K27me3 and co-compartmentalize with mB4 in the condensin-deficient mitotic chromosomes.

### Attraction-segregation plots

Attraction-repulsion plots were generated by transforming the 25kb binned *cis*-observed/expected contact matrix. To begin with, each column of bins was re-ordered so that bins assigned with the same type of compartments were placed together. Within each type of domain, the original order of bins was maintained so that one may deduce position dependent contact information (e.g. pericentromeric region vs. telomeric region) from an attraction-repulsion plot. The same order of bins was then applied to each row to generate a symmetrical matrix. For an individual chromosome, aggregated bins of each compartment were then divided into 40 equal sections. In this way, compartments from different chromosomes may be averaged together to generate a genome-wide attraction-repulsion plot. In practice, we generated attraction-repulsion plots for mA1, mA2, mB4 and mB1-type compartments. Both homotypic intra-domain and heterotypic inter-domain interactions can be visualized through attraction-repulsion plots.

### Position dependent attraction-segregation plots

To investigate compartmentalization pattern at different regions of a chromosome, position dependent attraction-repulsion plots were generated. In brief, an on-diagonal square window of 15Mb was cropped from the Hi-C contact matrix at the pericentromeric region (starting from 3Mb in our case). The 15Mb window was then scanned through the chromosome toward telomere with a step-size of 1Mb. A small scale attraction-repulsion plot was generated within the window for each step as described above. Homotypic or heterotypic interactions within each window for each sample was subtracted by signals from the prometaphase samples from the parental cell line and were then plotted against chromosome position.

### CRE contact stratification

To acquire interphase CRE contacts, we adopted a previously characterized CRE list containing 22,294 CREs (enhancers and promoters defined by H3K27ac peaks) genome-wide. The CREs were then assigned to 10kb genomic bins. Bins containing CREs were randomly paired up with a selected range of separation (70kb to 300kb). The candidate CRE pairs were then subject to HICCUPS to obtain the observed and donut expected values in asynchronous and condensin-deficient mitotic samples. The candidate pairs were then subject to three additional filters: (1) candidate CRE pair must contain >0 reads in all tested samples (asynchronous and all mitotic time points); (2) the strength of the candidate CRE pair in at least one of the following samples (asynchronous sample and 30min, 1h, 4h and 8h condensin-deficient mitotic samples) must be at least 1.5 fold higher than the strength of this CRE pair in the 0h condensin replete mitotic samples; (3) the observed/expected value in the candidate CRE pair must be over 1.5 in the asynchronous sample. In the end, we obtained 6,825 CRE contacts.

### Contact probability decay curves (*Ps* curves)

The *Ps* curve of each individual chromosome was generated using the “expected-cis” function of Cooltools (v0.4.0). The genome-wide *Ps* curve was computed by averaging individual chromosomes together.

### Aggregated plots for loops and domains

Aggregated plots were generated using the python package Coolpup (v0.9.7). For unscaled aggregated peak analysis (APA), loops smaller than 70kb were removed from the plots to avoid influence from pixels close to the diagonal.

### Data availability

The Hi-C, Cut&Tag and ChIP-seq data generated in this study are deposited at GEO database with the accession numbers GSE227816 and GSE228402. External ChIP-seq data from previous studies are listed as below: H3K27ac (GSE61349) ^43^, H3K4me1 (GSM946535), H3K4me3 (GSM946533), H3K36me3 (GSM946529), H3K9me3 (GSM946542), H3K27me3 (GSM946531) ^44^ and RNA PolII (GSE168251) ^3^. External Hi-C data of CTCF depleted and complete G1E-ER4 cells are available at GSE168251 ^3^. External Hi-C data of prometaphase G1E- ER4 parental cell line is available at (GSE129997) ^2^.

### Code availability

Code will be available upon requests.

## FIGURE LEGENDS

**Extended Data Figure 1.**
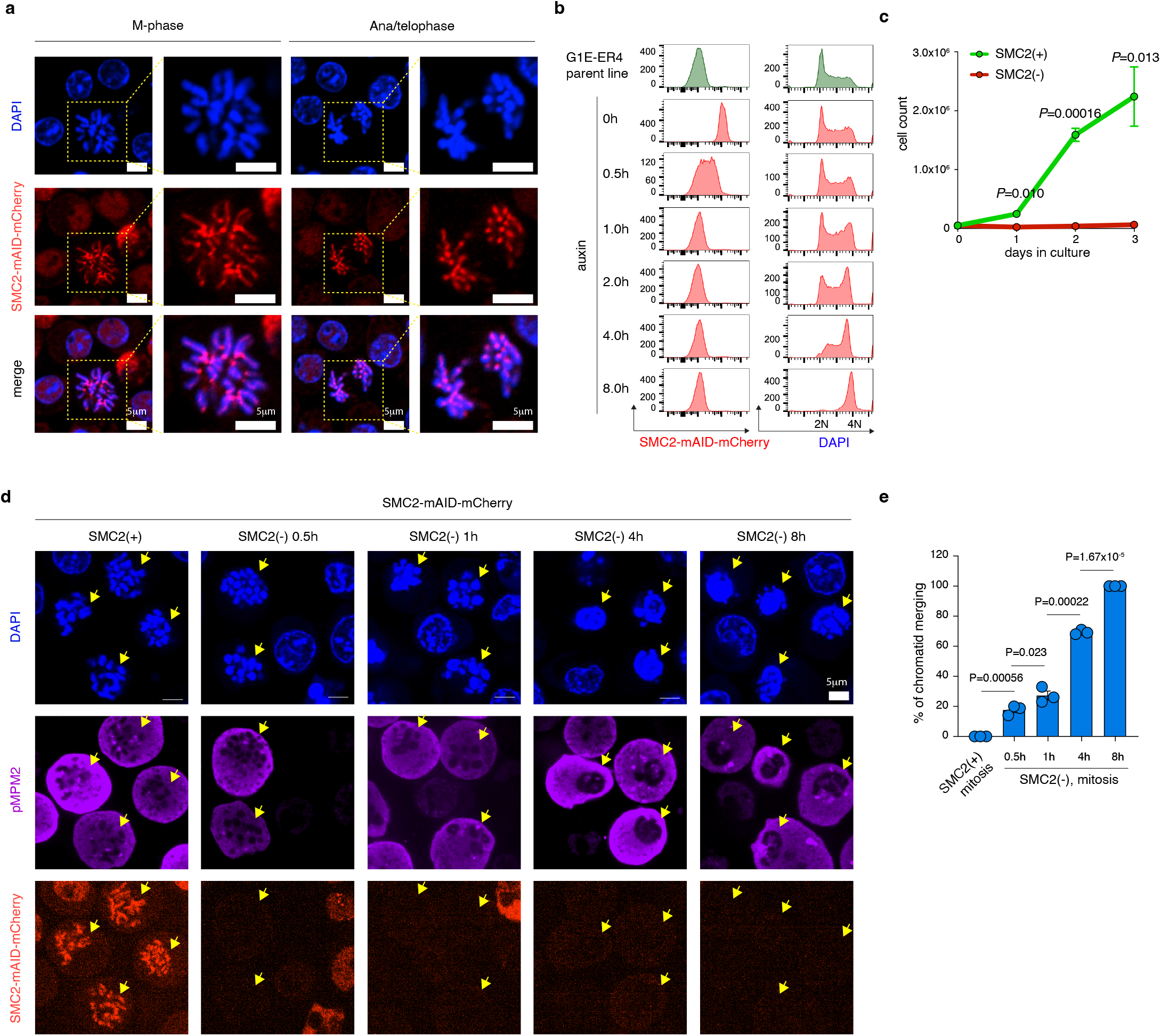
Characterization of the G1E-ER4:SMC2-AID-mCherry cell line. **a**, Left panel: representative image showing the condensation of mitotic chromosomes and correct nuclear localization of SMC2-mAID-mCherry fusion protein. Scale bar: 5*μ*m. Right panel: representative image showing the correct separation of chromatids during ana/telophase. Scale bar: 5*μ*m. Three independent experiments replicates were performed. **b**, Left panel: flow cytometry plots showing the rapid degradation of SMC2-mAID-mCherry fusion protein upon auxin treatment. Right panel: flow cytometry plot showing the shift of cell cycle distribution upon long-term auxin treatment. Two independent experiments were performed. **c**, Growth curve of G1E-ER4:SMC2- mAID-mCherry cells with or without auxin treatment. Error bar denotes SEM (n=3). *P* values were calculated using a two-sided student’s t test. **d**, Representative image showing the morphology of mitotic chromosomes after auxin treatment for indicated durations. Cells were stained with anti-pMPM2 antibody to indicate mitotic population. Scale bar: 5*μ*m. Three independent experiments were performed. **e**, Bar graph showing the percentage of mitotic cells with extensive chromatid entanglements after auxin treatment for indicated durations. Error bar denotes SEM (n=3). *P* values were calculated using a two-sided student’s t test.

**Extended Data Figure 2.**
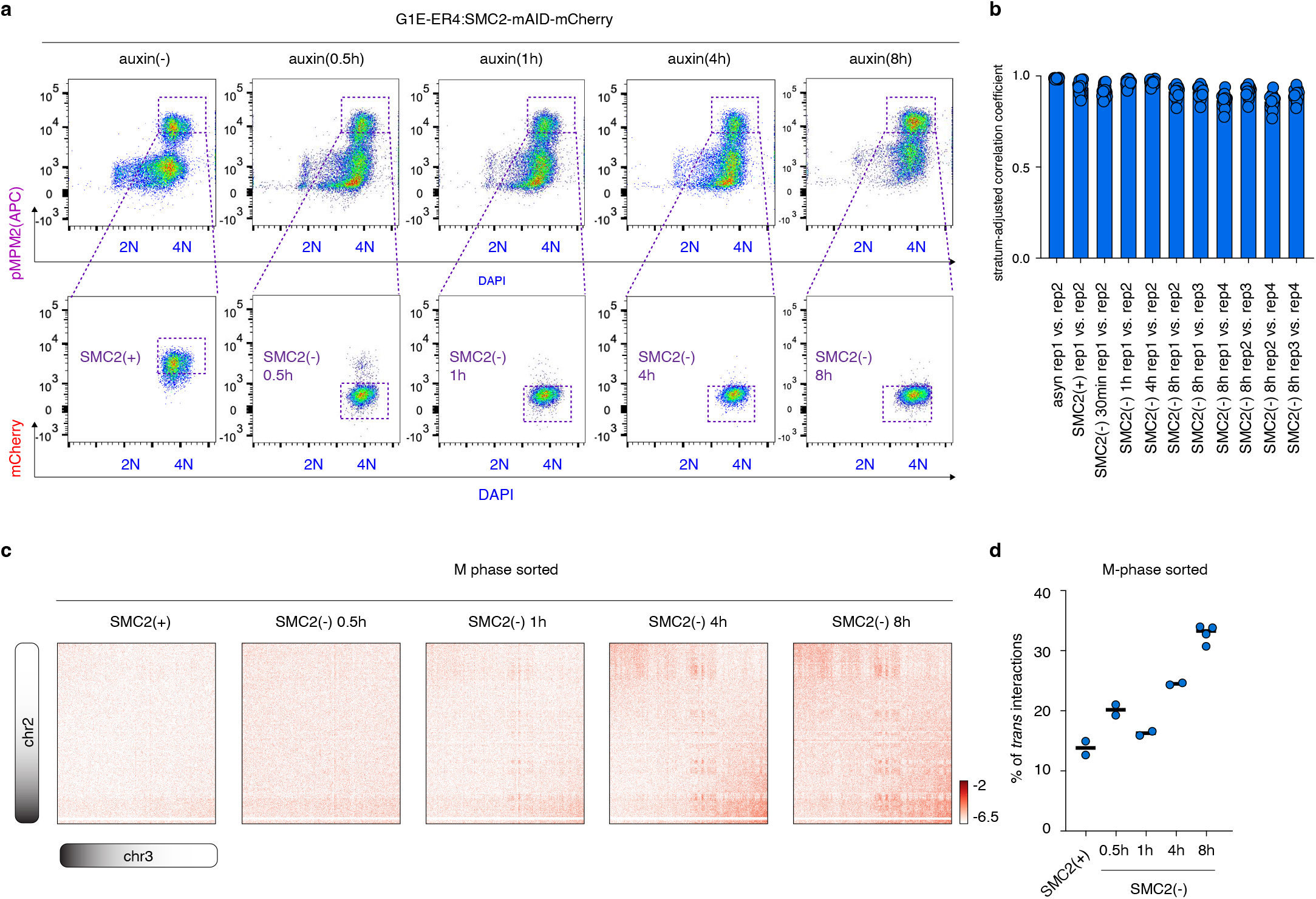
Purification of condensin-deficient mitotic cells. **a**, Flow cytometry plot showing the gating strategy to purify mitotic cells at different time points after addition of auxin. Cells were sorted by pMPM2, DAPI and mCherry signals. More than two independent experiments were performed. **b**, Bar graph showing the high stratum-adjusted correlation coefficient for each chromosome (n=19) between biological replicates for each condition. **c**, KR balanced Hi-C contact matrices (chr2 vs. chr3) showing the gradual increase of *trans*-chromosome interactions over time upon condensin loss. **d**, Dot plot showing the percentage of *tran*-chromosome interactions for independent biological replicates at each tested time point. n=2 for SMC2(+), 0.5h, 1h and 4h auxin treated samples. n=4 for 8h auxin treated samples.

**Extended Data Figure 3.**
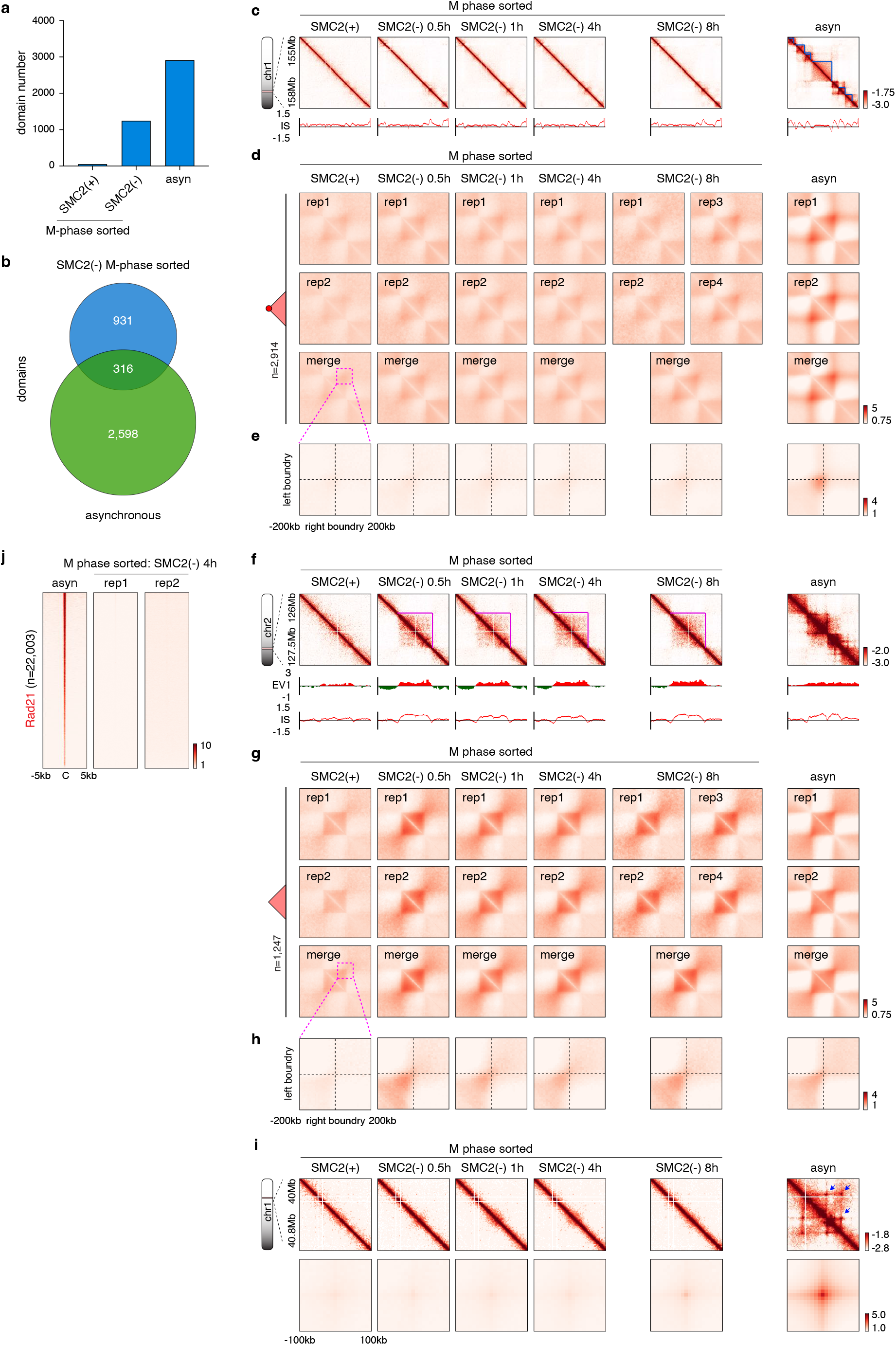
TADs and structural loops are not resumed in condensin-deficient mitotic cells. **a**, Bar graph showing the number of domains identified in control mitotic, SMC2 deficient mitotic and asynchronous cells. **b**, Venn-diagram showing the intersection results of domains for SMC2 deficient mitotic samples and asynchronous control samples. **c**, KR-balanced Hi-C contact maps showing representative domains (chr1:155-158Mb) identified in asynchronously growing cells. Bin size: 10kb. Browser tracks of insulation score profiles were shown. TAD was highlighted by blue lines. **d**, Composite contact plots of the rescaled 2,914 domains identified in asynchronous cells. Plots for independent biological replicates as well as replicate-merged samples were shown. **e**, Aggregated peak analysis showing the strong summit corner dots for domains identified in the asynchronous control cells. **f**, KR-balanced Hi-C contact maps showing the emergence of a representative domains (chr2:126-127.5Mb) in the SMC2 deficient mitotic cells. Bin size: 10kb. Browser tracks of EV1 values and insulation score profiles were shown. Domain was highlighted by pink lines. **g**, Composite contact plots of the rescaled 1,247 domains identified in SMC2 deficient mitotic cells. Plots for independent biological replicates as well as replicate-merged samples were shown. **h**, Aggregated peak analysis showing lack of summit corner dots for domains identified in the SMC2 deficient mitotic cells. **i**, Upper panel: KR-balanced Hi-C contact maps showing representative structural loops (chr1:40-40.8Mb) in condensin deficient mitotic cells and asynchronous control cells. Bin size: 10kb. Lower panel: Aggregated peak analysis for structural loop signals (n=4,837) in the SMC2 deficient mitotic cells. **j**, Density heatmaps showing loss of cohesin positioning in the condensin-deficient (4h) mitotic cells.

**Extended Data Figure 4.**
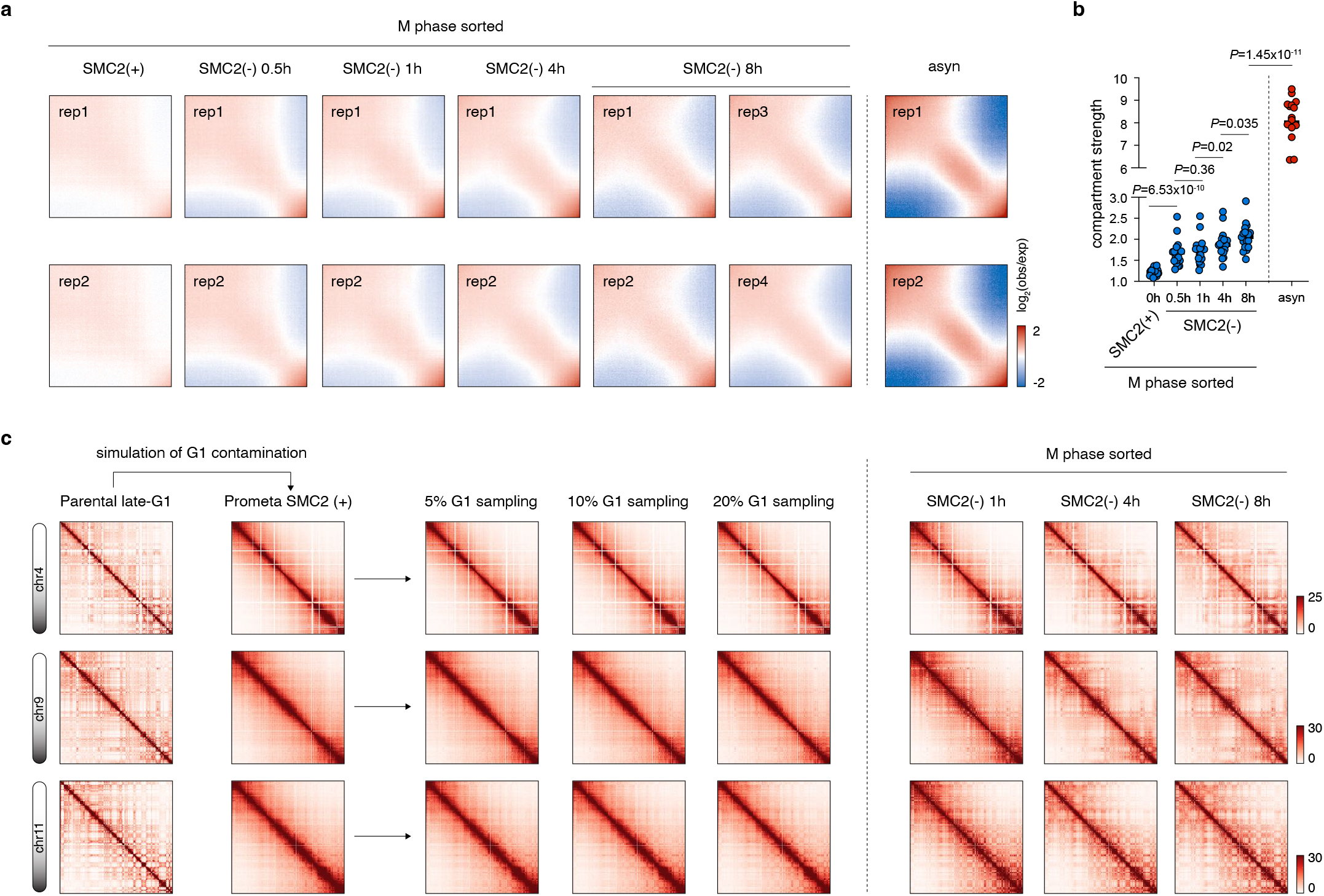
Progressively strengthened compartments in mitotic cells upon condensin loss. **a**, Saddle plot of independent biological replicates of asynchronous and condensin deficient mitotic cells. **b**, Dot plot showing progressive gain of compartmental strength of each individual chromosome (n=20). *P* values were calculated using a two-sided paired Wilcoxon signed-rank test. **c**, Raw Hi-C contact matrices showing the results of in-silicon simulation of various level of G1 contamination in mitotic control cells. Note that even 20% of G1 contamination, failed to show as strong compartmentalization as mitotic cells without condensin.

**Extended Data Figure 5.**
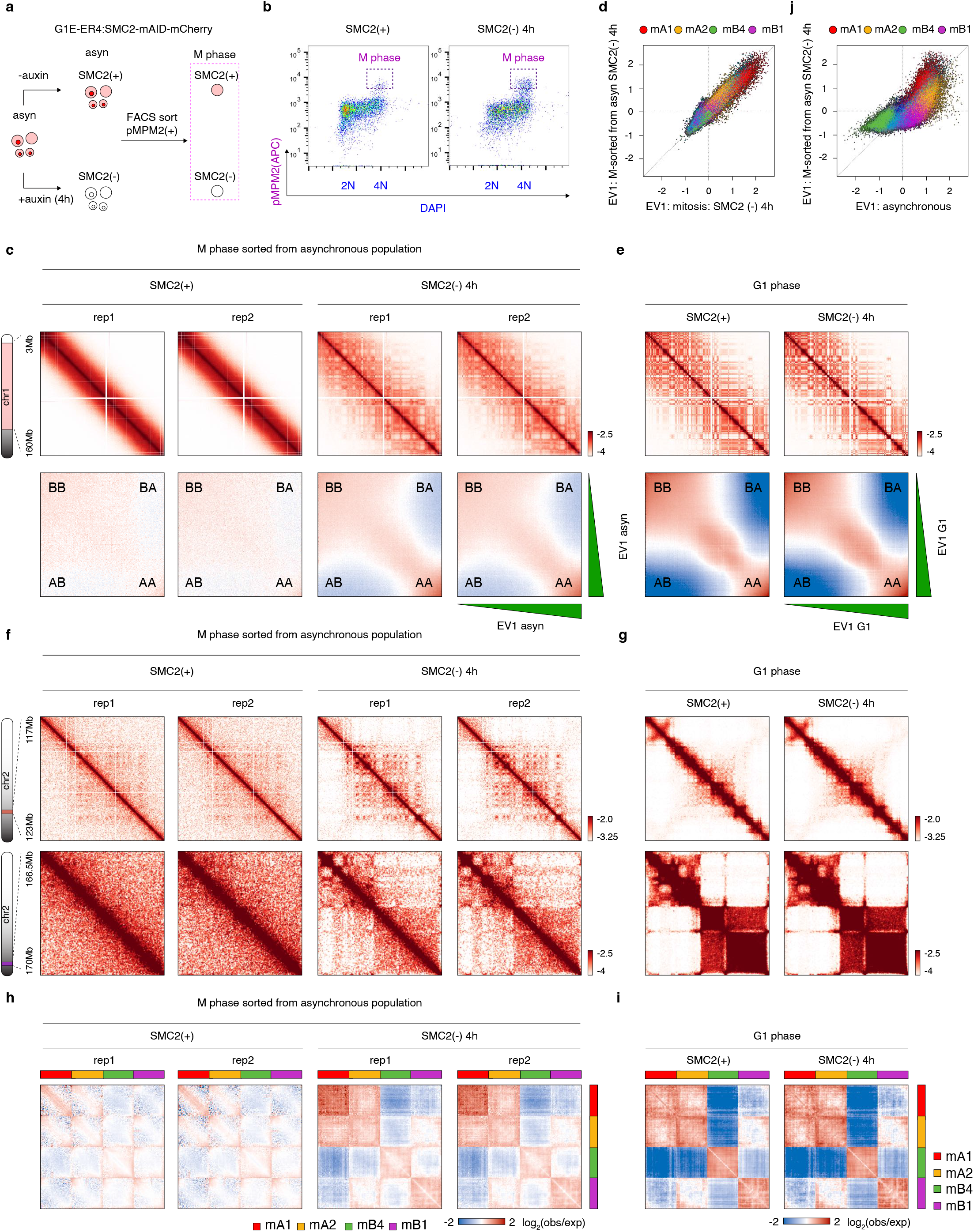
Compartmentalization of mitotic chromosomes from asynchronous population. **a**, Schematic showing the sorting strategy of condensin-deficient and control mitotic cells from asynchronous populations. **b**, Flow cytometry plot showing the gating strategy to purify mitotic cells from asynchronous population. Two independent experiments were performed. **c**, Upper panel: KR balanced Hi-C contact matrices (chr1:3-160Mb) of condensin-deficient (4h) and control mitotic cells from asynchronous population. Bin size: 100kb. Lower panel: Saddle-plots showing the compartment strength in the condensin-deficient (4h) and mitotic cells from asynchronous population. Each biological replicate is shown. **d**, Scatter plot showing the high correlation of EV1 values between condensin-deficient (4h) mitotic cells from nocodazole arrested (*x*-axis) vs. asynchronous populations (*y*-axis). **e**, Upper panel: KR balanced Hi-C contact matrices (chr1:3- 160Mb) of condensin-deficient (4h) and control G1 phase cells. Bin size: 100kb. Lower panel: Saddle-plots showing the compartment strength in the condensin-deficient (4h) and control G1 mitotic cells. **f**, Upper panel: KR balanced Hi-C contact maps (chr2:117-123Mb) showing representative mA1 homotypic interactions in condensin-deficient (4h) mitotic cells from asynchronous population. Lower panel: KR balanced Hi-C contact maps (chr2:156.5-170Mb) showing mB1 and mB4 co-compartmentalization in condensin-deficient (4h) mitotic cells from asynchronous population. Bin size: 25kb. **g**, KR balanced Hi-C contact maps showing the same regions in (**f**) in condensin-deficient (4h) and control G1 phase cells. Bin size: 25kb. **h**, Attraction-repulsion plots of condensin-deficient (4h) and control mitotic cells from asynchronous population. Each biological replicate is shown. **i**, Attraction-repulsion plots of condensin-deficient (4h) and control G1 cells. **j**, Scatter plot showing the EV1 values of 25kb genomic bins in asynchronous control cells (*x*-axis) against condensin-deficient (4h) mitotic cells from asynchronous population (*y*-axis). Bins were color coded based on their compartment assignment.

**Extended Data Figure 6.**
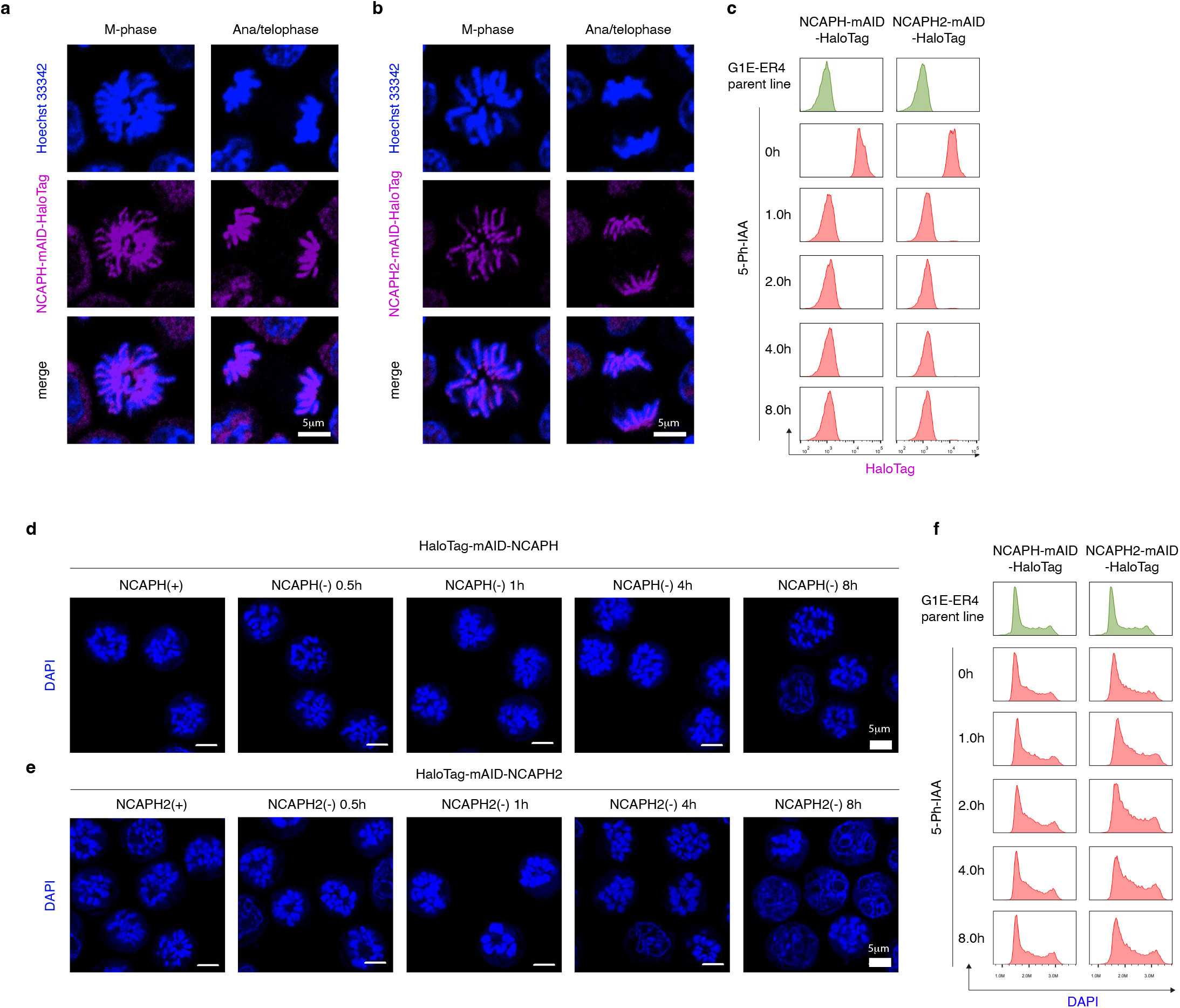
Characterization of G1E-ER4:NCAPH-mAID-HaloTag and G1E- ER4:NCAPH2-mAID-HaloTag cell lines. **a**, Representative image showing the correct localization of NCAPH-mAID-HaloTag fusion protein (Halo-646) in mitosis and ana/telophase. Scale Bar: 5*μ*m. Two independent experiments were performed. **b**, Representative image showing the correct localization of NCAPH2-mAID-HaloTag fusion protein (Halo-646) in mitosis and ana/telophase. Scale Bar: 5*μ*m. Two independent experiments were performed. **c**, Flow cytometry plots showing the rapid degradation of NCAPH-mAID-HaloTag and NCAPH2-mAID-HaloTag upon 5-Ph-IAA treatment. Two independent experiments were performed. **d**, Representative image showing the morphology of mitotic chromosomes after NCAPH degradation for indicated durations. Scale Bar: 5*μ*m. Two independent experiments were performed. **e**, Representative image showing the morphology of mitotic chromosomes after NCAPH2 degradation for indicated durations. Scale Bar: 5*μ*m. Two independent experiments were performed. **f**, Flow cytometry showing the cell cycle distribution after NCAPH or NCAPH2 are degraded for indicated durations. Two independent experiments were performed.

**Extended Data Figure 7.**
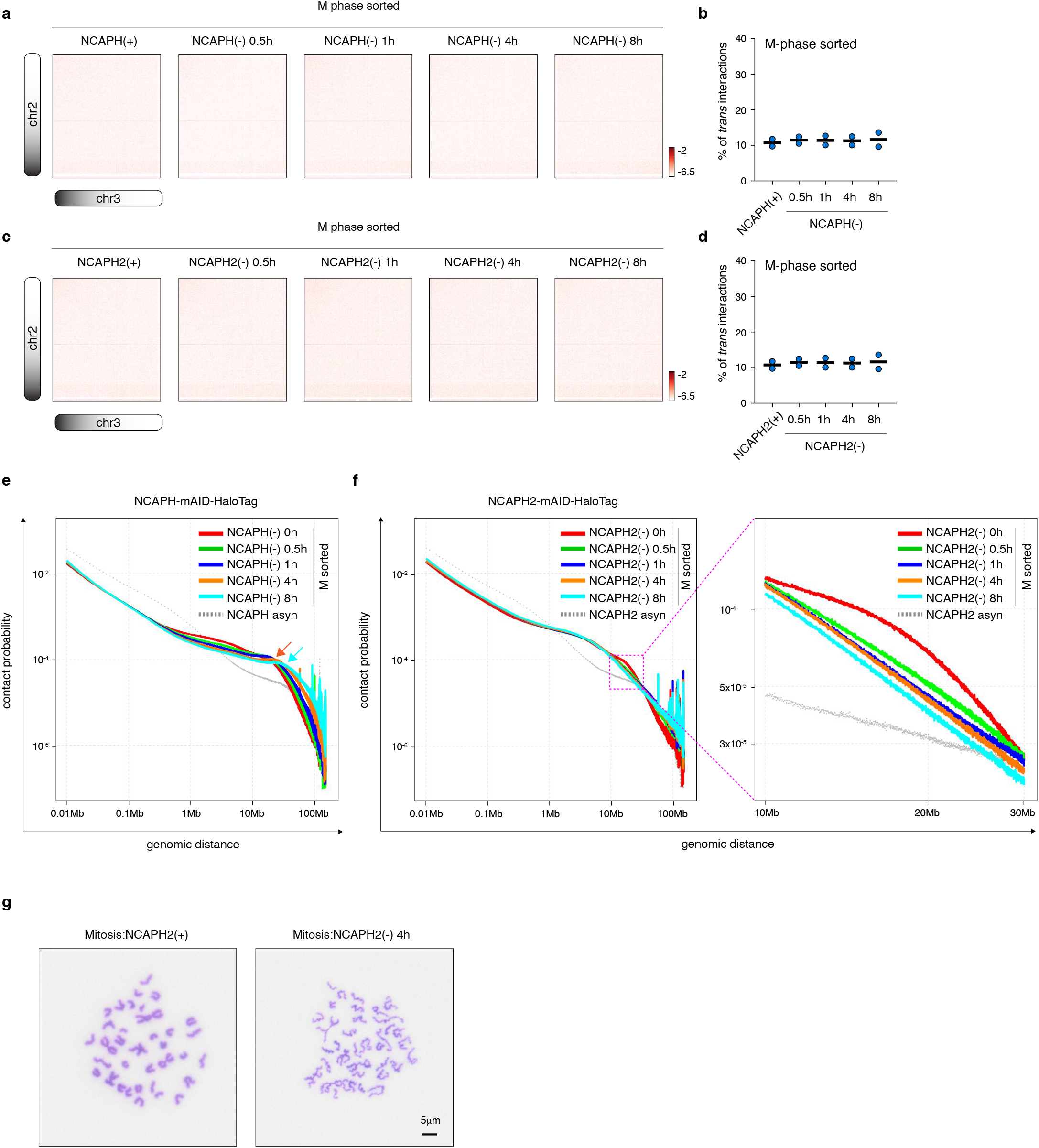
Characterization of NCAPH or NCAPH2 deficient mitotic chromosomes. **a**, KR balanced Hi-C contact matrices (chr2 vs. chr3) showing *trans*-chromosome interactions in mitotic cells upon NCAPH depletion. Bin size: 100kb. **b**, Dot plot showing the percentage of *tran*- chromosome interactions for independent biological replicates in NCAPH deficient mitotic cells. **c**, KR balanced Hi-C contact matrices (chr2 vs. chr3) showing *trans*-chromosome interactions in mitotic cells upon NCAPH2 depletion. Bin size: 100kb. **d**, Dot plot showing the percentage of *tran*-chromosome interactions for independent biological replicates in NCAPH2 deficient mitotic cells. **e**, *P(s)* curve showing progressive reconfiguration of mitotic chromosomes after NCAPH loss. **f**, Left panel: *P(s)* curve showing progressive reconfiguration of mitotic chromosomes after NCAPH2 loss. Right panel: enlarged plot of left panel showing genomic separations between 10- 30Mb. **g**, Chromatin spread analysis showing the mitotic chromosome morphology in control and NCAPH2 deficient (4h) cells. Scale Bar: 5*μ*m

**Extended Data Figure 8.**
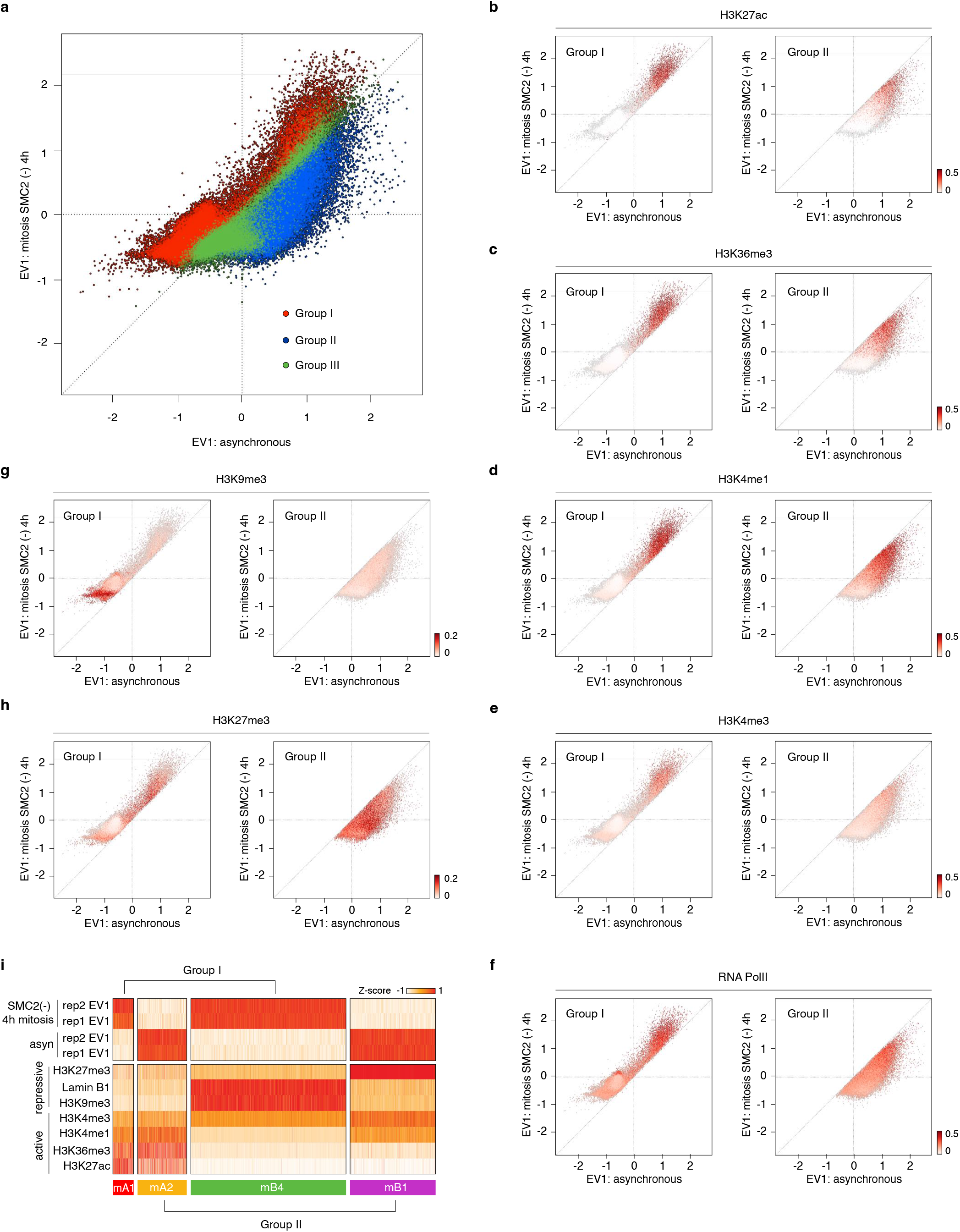
Initial clustering of mitotic specific compartments based on EV1 values. **a**, Scatter plot showing the EV1 values of 25kb genomic bins in asynchronous control cells (*x*-axis) against condensin-deficient (4h) mitotic cells (*y*-axis). Bins belong to group I, II and III were marked by red, blue and green colors respectively. **b**-**f**, Scatter plot showing the enrichment of indicated histone modification intensity or RNA Polymerase binding intensity for 25kb genomic bins (group I and II) in asynchronous control cells (*x*-axis) against condensin-deficient (4h) mitotic cells (*y*-axis). **i**, Heatmap showing the clustering result to group the genome into different mitotic specific compartments. The upper panel describes results from the initial clustering based on EV1 values. The lower panel illustrates the second step of clustering using chromatin-associating features.

**Extended Data Figure 9.**
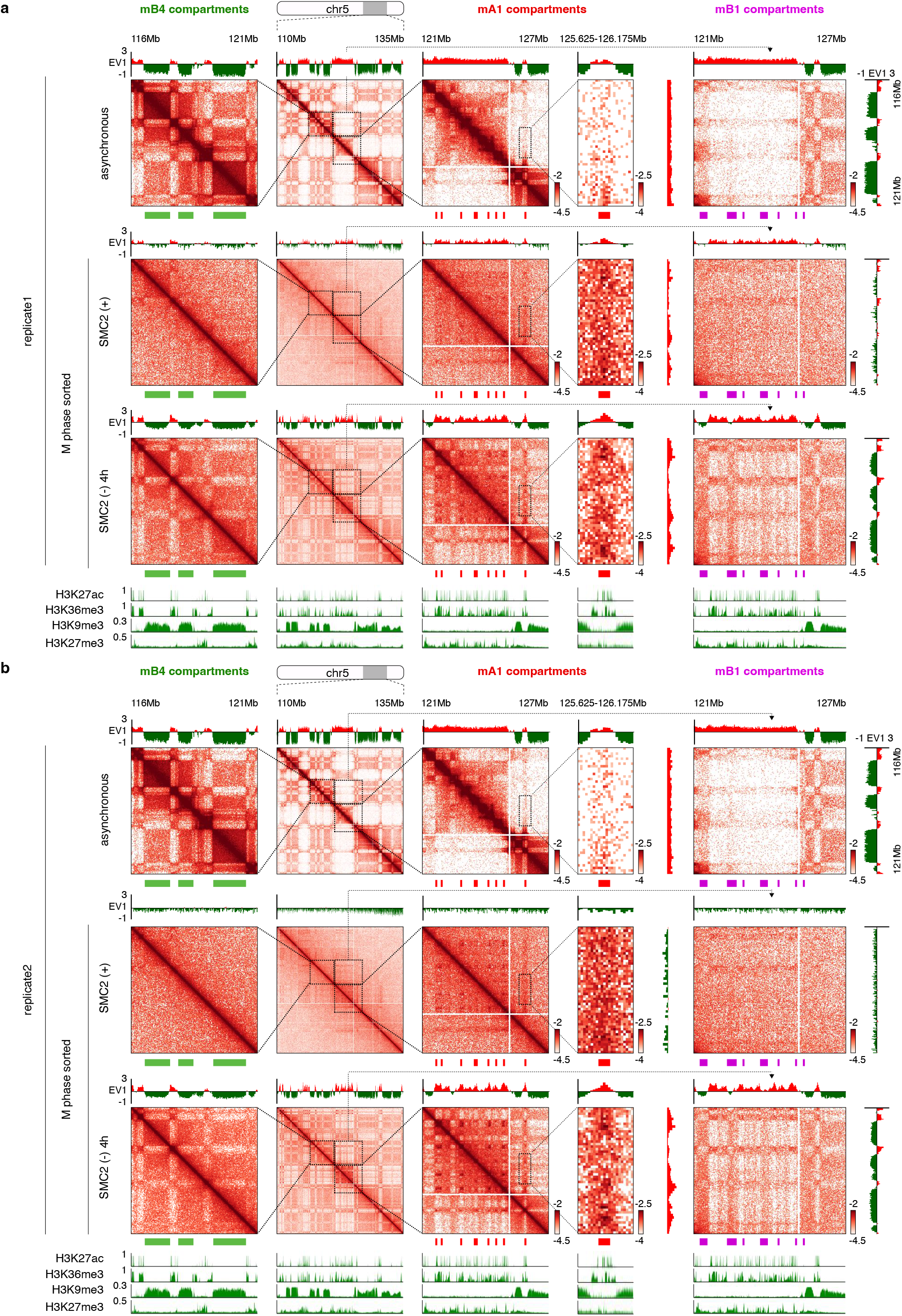
Additional examples of mA1-, mB4- and mB1-compartments in a single view of Hi-C contact map. **a**, KR balanced Hi-C contact matrices showing a genomic locus (chr5:110-135Mb) to illustrate homotypic and heterotypic interactions of compartment mA1, mB4 and mB1 in control mitosis, condensin-deficient (4h) mitosis and asynchronous control samples (replicate 1). Homotypic interactions of mA1 (chr5:121-127Mb) and mB1 (chr5:116-121Mb) compartments as well as heterotypic interactions between mB4 and mB1 (chr5:116-121Mb vs. chr5:121-127Mb) compartments were highlighted by dotted boxes. Bin size: 25kb. mA1, mB4 and mB1 compartments were indicated by red, green and purple bars respectively. Browser tracks of EV1 values for each condition as well as H3K27ac, H3K36me3, H3K9me3 and H3K27me3 from asynchronously growing cells were shown. **b**, Similar to (**a**) showing Hi-C maps of biological replicate 2.

**Extended Data Figure 10.**
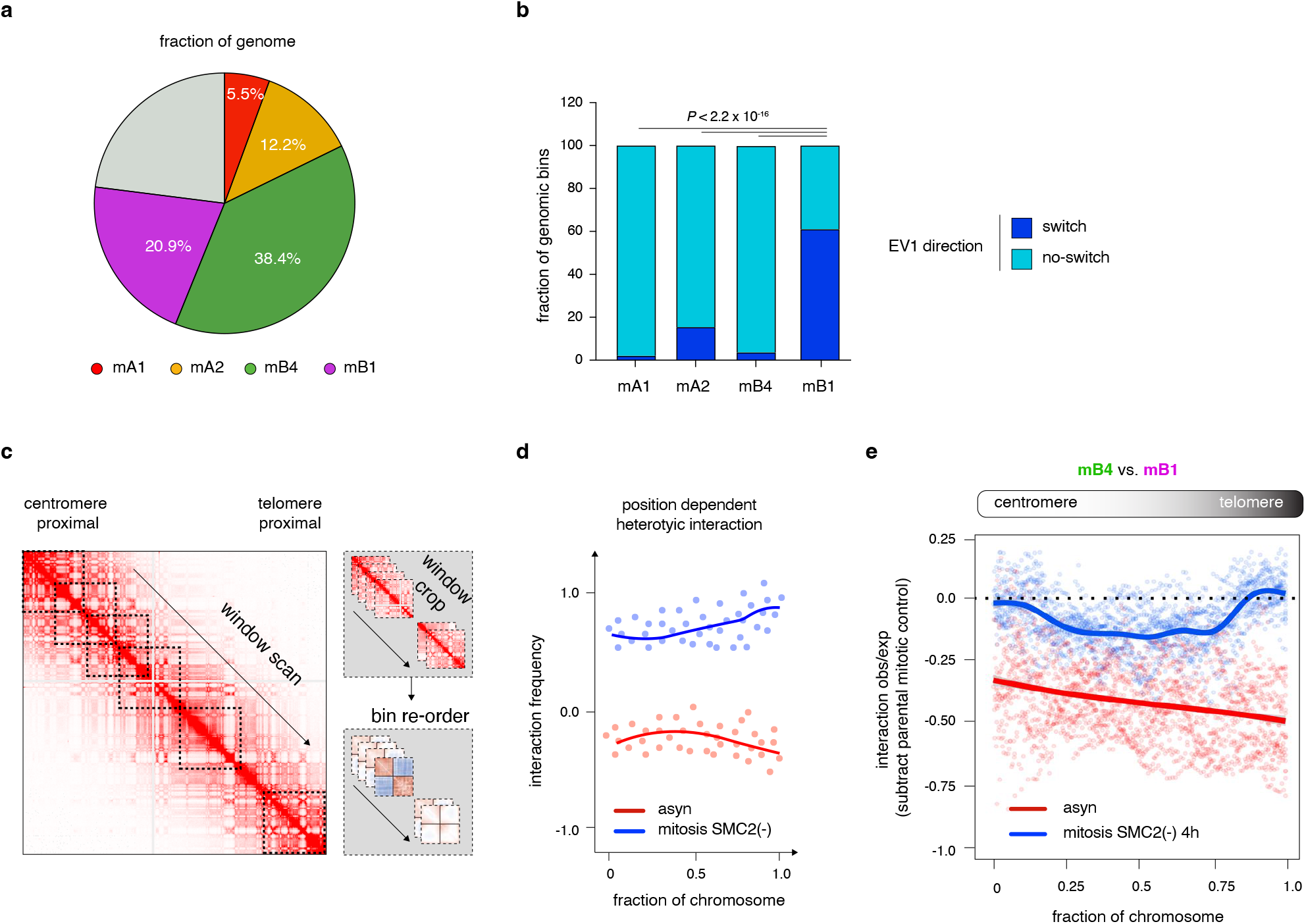
Characterization of repressive compartments in the condensin-deficient mitotic cells. **a**, Pie chart showing the fraction of different mitotic compartments in the genome. **b**, Bar graph showing the percentage of genomic bins with inverted EV1 values during mitosis in each type of compartments. *P* values were calculated using a two-sided Fisher’s exact test. **c**, Schematic illustration showing how to generate the chromosome-location dependent attraction-repulsion curve. **d**, Pseudo-data showing analytical results of (**c**). **e**, Chromosome-location dependent attraction-repulsion curve between mB1 and mB4 compartments for condensin-deficient (4h) mitotic and asynchronous cells.

**Extended Data Figure 11.**
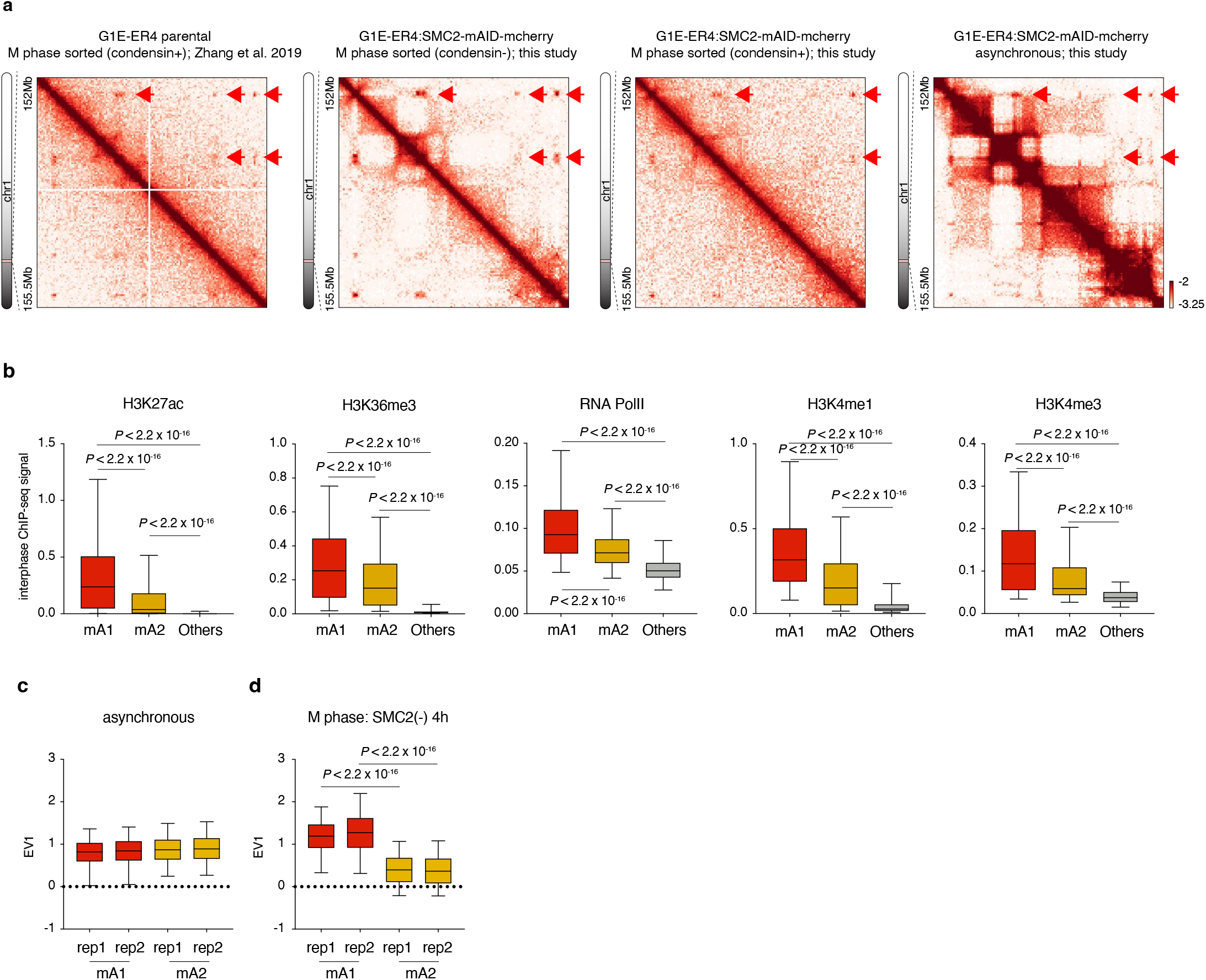
Characterization of active compartments in the condensin-deficient mitotic cells. **a**, KR balanced Hi-C contact matrices showing the prominent interactions among mA1 compartment (chr1:152-155.5Mb) in the condensin replete mitotic samples (from this study and a prior study by Zhang et al.^2^) as well as condensin-deficient (4h) mitotic samples and asynchronous samples. **b**, Box-plots showing the enrichment of indicated chromatin associating features (histone modification and transcription) for mA1 (n=5,299 genomic bins), mA2 (n=11,531 genomic bins) compartments and the rest of the genome (n=77,997 genomic bins). For all box plots, central lines denote medians; box limits denote 25th–75th percentile; whiskers denote 5th–95th percentile. *P* values were calculated using a two-sided Wilcoxon signed-rank test. **c**, Box-plots showing the asynchronous EV1 values for mA1 (n=5,299 genomic bins) and mA2 (n=11,531 genomic bins) compartments for both biological replicates. For all box plots, central lines denote medians; box limits denote 25th–75th percentile; whiskers denote 5th–95th percentile. **c**, Box-plots showing the EV1 values for mA1 (n=5,299 genomic bins) and mA2 (n=11,531 genomic bins) compartments in the condensin-deficient (4h) mitotic cells for both biological replicates. For all box plots, central lines denote medians; box limits denote 25th–75th percentile; whiskers denote 5th–95th percentile. *P* values were calculated using a two-sided Wilcoxon signed-rank test.

**Extended Data Figure 12.**
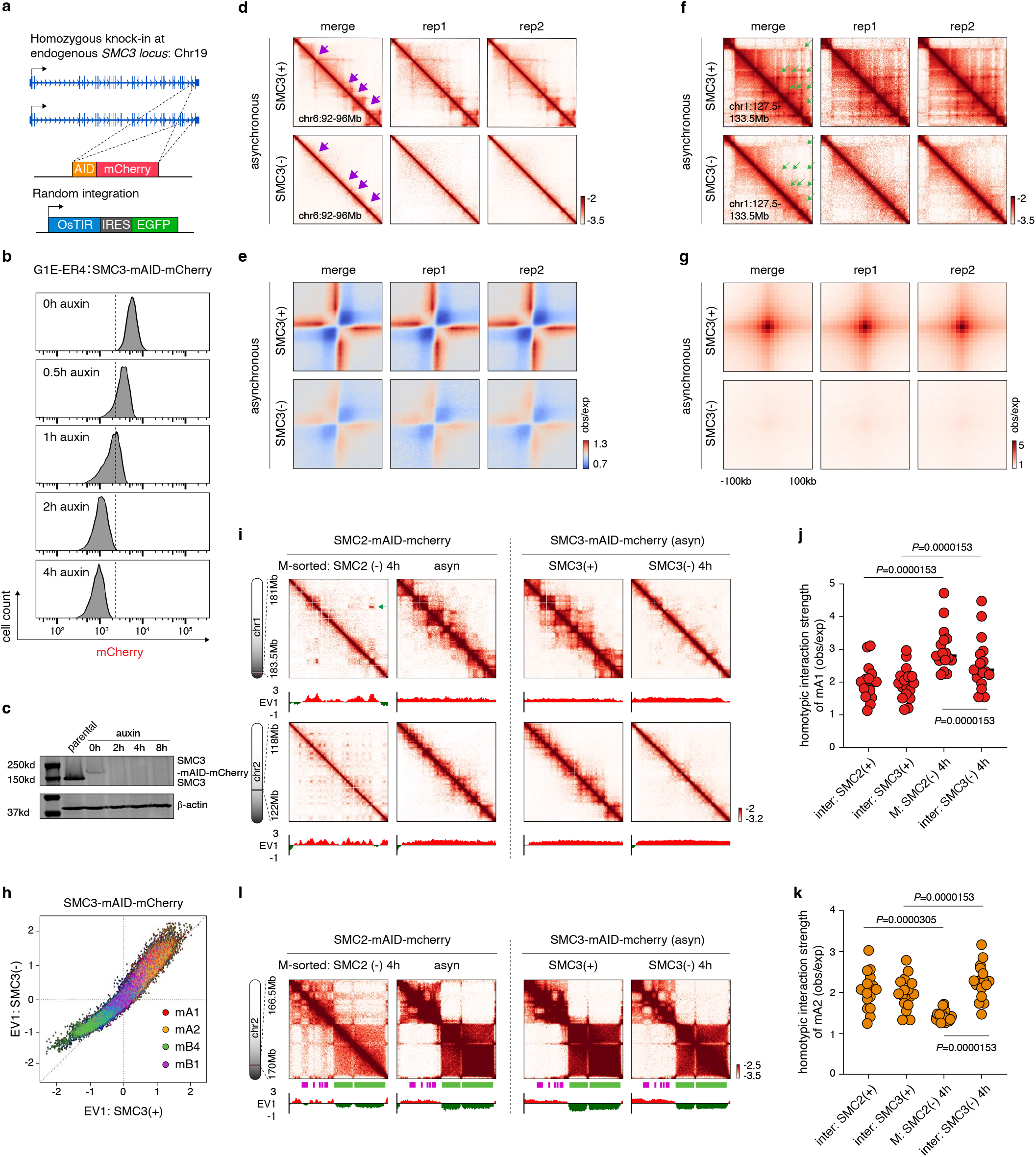
Comparison of the “extrusion-free” mitotic and interphase chromatin. **a**, Schematic illustration showing homozygous insertion of mAID-mCherry tag at the C terminus of endogenous SMC3. **b**, Flow-cytometry showing the rapid degradation of SMC3 upon auxin treatment. Two independent experiments were performed. **c**, Western blot showing the degradation of SMC3 upon auxin treatment. One experiment was performed. **d**, KR balanced Hi-C contact matrices (chr6:92-96Mb) showing the loss of TADs boundaries upon SMC3 depletion. Contact maps of both biological replicates and replicate-merged samples were shown. Bin size: 10kb. TAD boundaries were indicated by purple arrows. **e**, Composite contact map of TAD boundary showing reduced insulation after SMC3 depletion. **f**, KR balanced Hi-C contact matrices (chr1:127.5- 133.5Mb) showing elimination of chromatin loops upon SMC3 depletion. Contact maps of both biological replicates and replicate-merged samples were shown. Bin size: 10kb. Chromatin loops were indicated by green arrows. **g**, APA plots showing disappearance of structural loops upon SMC3 loss. **h**, Scatter plot showing the EV1 values of 25kb genomic bins in SMC3 (+) (*x*-axis) against SMC3 (-) G1 phase cells (*y*-axis). Bins were color coded by their compartment assignment. **i**, KR balanced Hi-C contact matrices (chr1:181-183.5Mb and chr2:118-112Mb) showing mild increase of mA1 homotypic interactions after SMC3 loss. Bin size: 25kb. Browser tracks of EV1 values were shown. **j**, Dot plot showing the strength of mA1 homotypic interactions in indicated samples. Each dot represents an individual chromosome (n=17). *P* values were calculated using a two-sided paired Wilcoxon signed-rank test. **k**, Dot plot showing the strength of mA2 homotypic interactions in indicated samples. Each dot represents an individual chromosome (n=17). *P* values were calculated using a two-sided paired Wilcoxon signed-rank test. **l**, KR-balanced Hi-C contact matrices (chr2:166.5-170Mb and chr2:118-112Mb) showing the lack of mB1-to-mB4 interactions in interphase cells without SMC3. mB1 and mB4 compartments were highlighted by purple and green bars respectively. Browser tracks of EV1 values were shown. Bin size: 25kb.

**Extended Data Figure 13.**
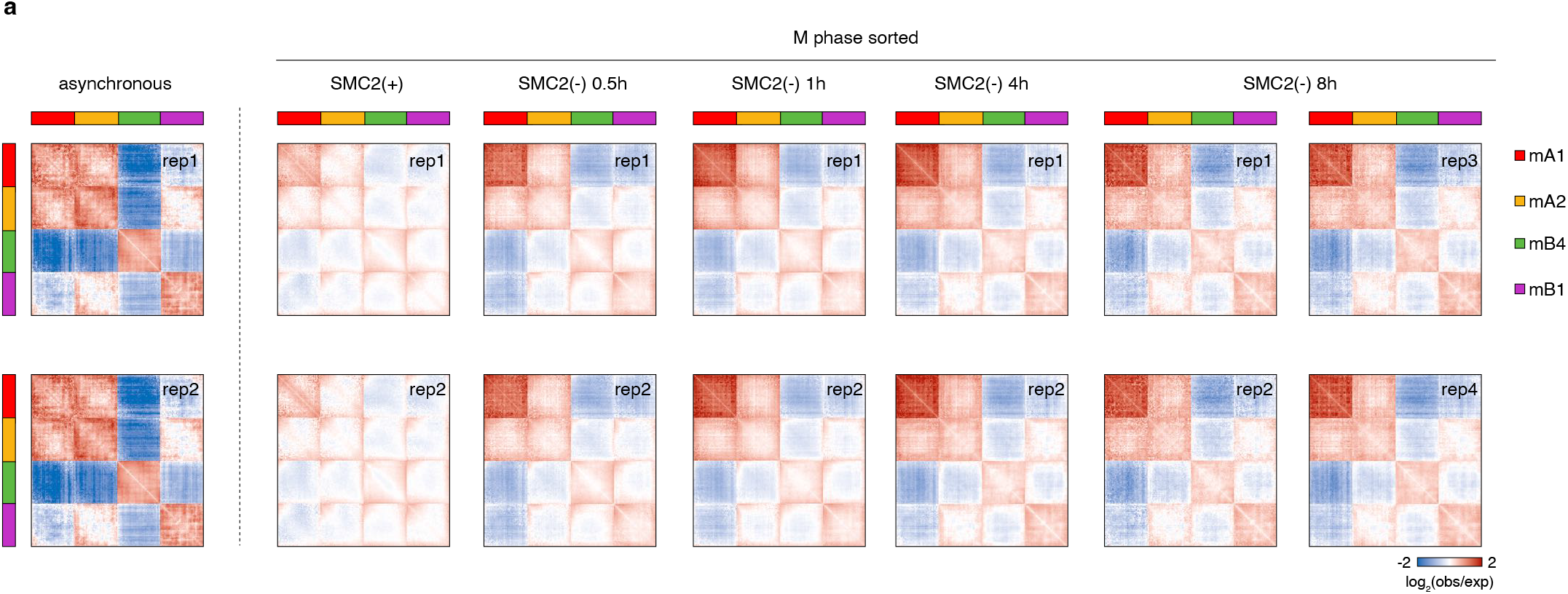
Dynamics of mitotic compartmentalization upon condensin loss. **a**, Attraction-repulsion plots showing the homotypic and heterotypic interactions among four different compartments: mA1, mA2, mB4 and mB1. Plots for biological replicates of control mitotic, condensin-deficient mitotic (all tested time points) and asynchronous cells were shown.

**Extended Data Figure 14.**
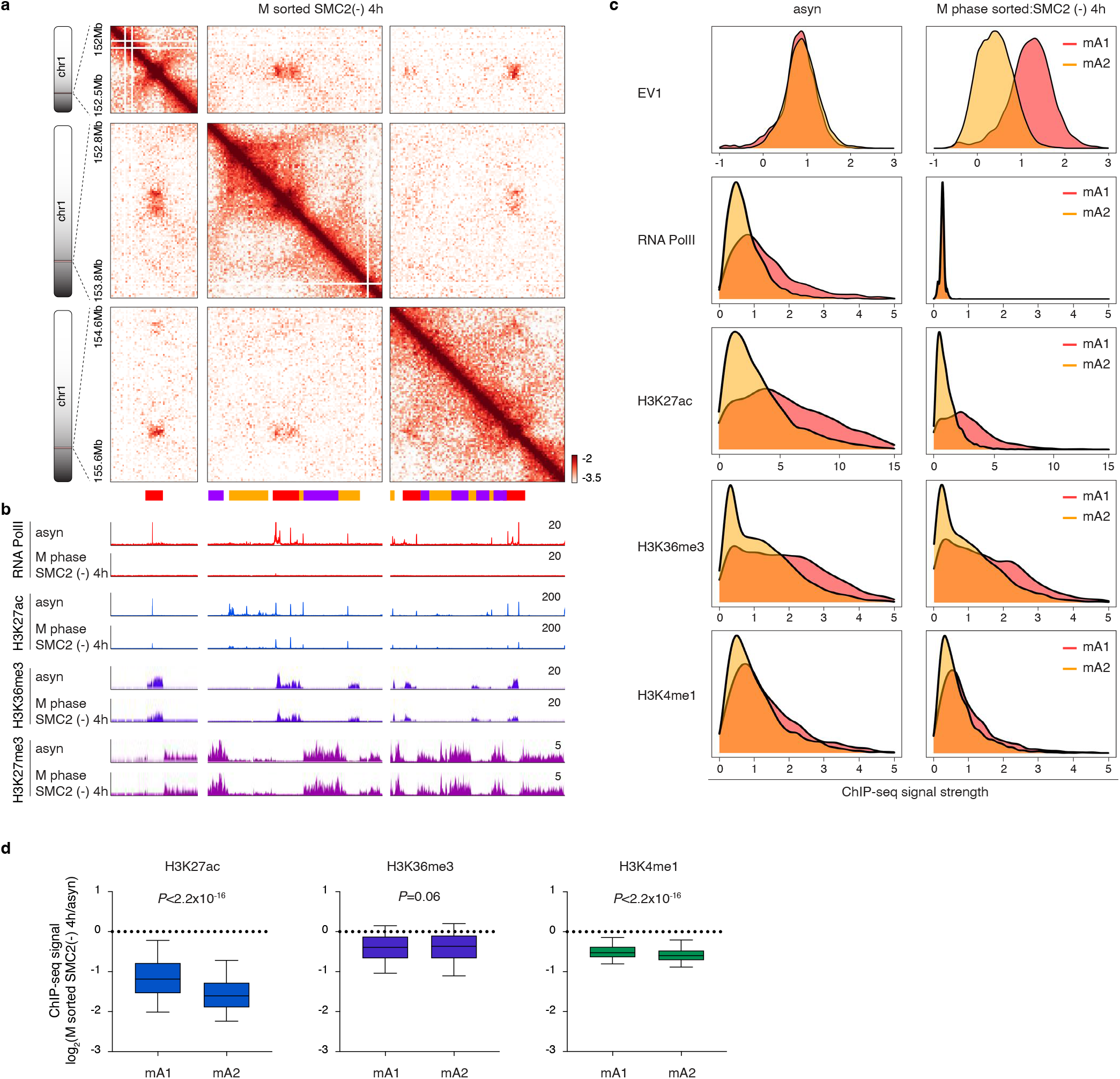
Epigenetic landscape of mitotic chromosomes in the absence of condensin. **a**, KR balanced Hi-C contact matrices (chr1:152-152.5Mb and chr1:152.8-153.8Mb and chr1:154.6-155.6Mb) showing mitotic compartments. Bin size: 25kb. mA1, mA2 and mB1 compartments are labeled by red, yellow and purple bars respectively. **b**, Representative tracks corresponding to the genomic regions in (**a**), showing the ChIP-seq profiles of RNA PolII, H3K27ac, H3K36me3 and H3K27me3 in asynchronous as well as condensin-deficient (4h) mitotic cells. **c**, Density plots showing the EV1 values as well as ChIP-seq signal strength of indicated marks in mA1 and mA2 compartments in the asynchronous and condensin-deficient mitotic cells. **d**, Box-plots showing the log_2_ fold change of indicated marks between condensin-deficient mitotic and asynchronous samples. For all box plots, central lines denote medians; box limits denote 25th– 75th percentile; whiskers denote 5th–95th percentile. *P* values were calculated using a two-sided Wilcoxon signed-rank test.

**Extended Data Figure 15.**
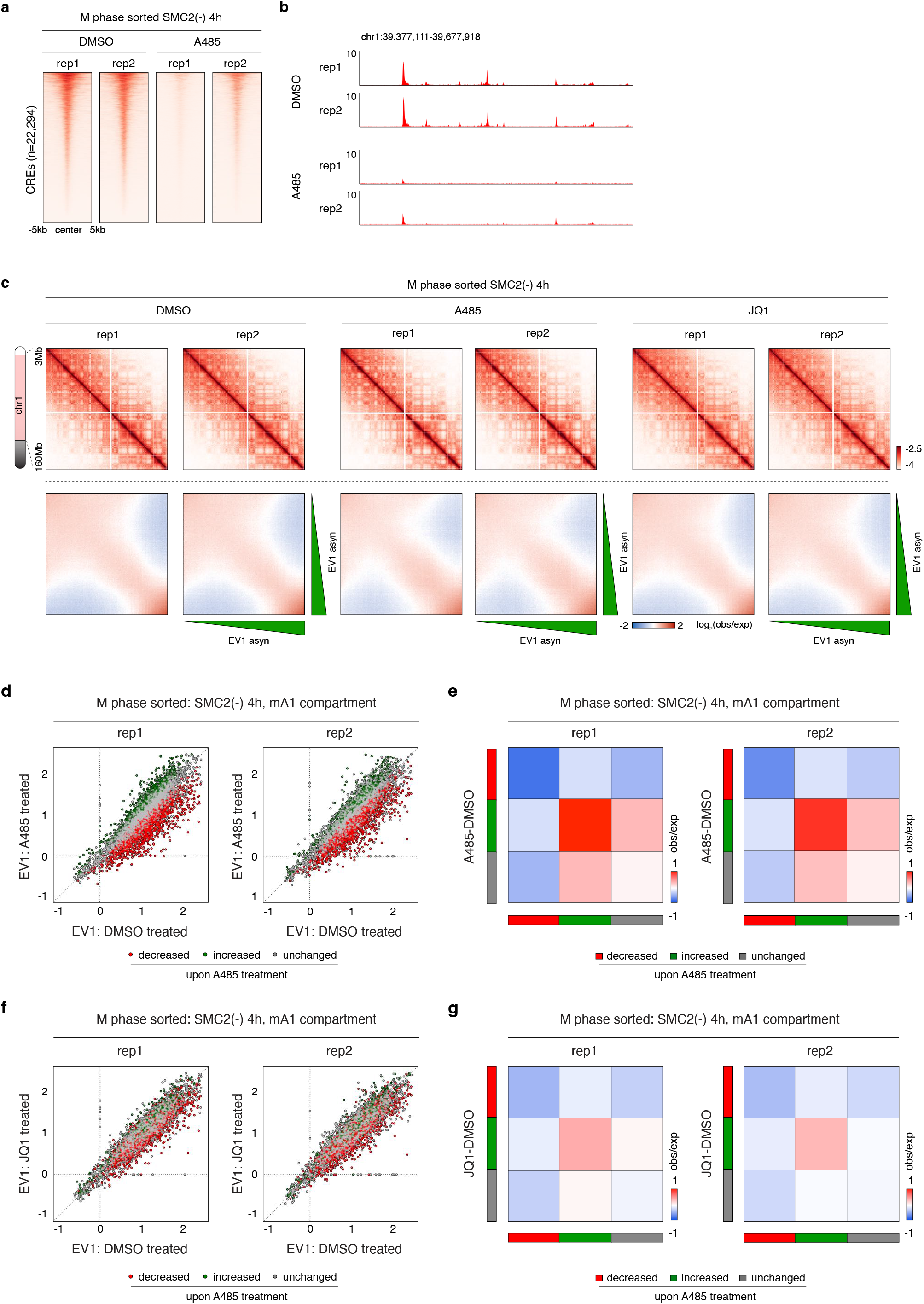
Effects of A485 and JQ1 treatment on condensin-deficient mitotic chromosome architecture. **a**, Density heatmaps showing loss of H3K27ac signals in the condensin-deficient (4h) mitotic chromosomes after A485 treatment. Individual biological replicates are shown. **b**, Genomic tracks of a representative region showing loss of H3K27ac signals in the condensin-deficient (4h) mitotic chromosomes after A485 treatment. Individual biological replicates are shown. **c**, Upper panel: KR balanced Hi-C contact matrices (chr1:3-160Mb) of condensin-deficient (4h) mitotic cells treated with DMSO, A485 or JQ1. Bin size: 100kb. Lower panel: Saddle-plots showing the compartment strength in the condensin-deficient (4h) mitotic cells treated with DMSO, A485 or JQ1. Independent biological replicates were shown. **d**, Scatter plot showing the EV1 values of mA1 25kb genomic bins in DMSO treated (*x*-axis) against A485 treated (*y*-axis) condensin-deficient (4h) mitotic cells. Bins were color coded based on their response to A485 treatment. Each biological replicate was shown. **e**, Heatmap showing the differential interaction strengths among mA1-D, mA1-I and mA1-U compartments upon A485 treatment compared to DMSO treated control. Each biological replicate was shown. **f**, Scatter plot showing the EV1 values of mA1 25kb genomic bins in DMSO treated (*x*-axis) against JQ1 treated (*y*-axis) condensin-deficient (4h) mitotic cells. Bins were color coded based on their response to A485 treatment. Each biological replicate was shown. **e**, Heatmap showing the differential interaction strengths among mA1-D, mA1-I and mA1-U compartments upon JQ1 treatment. Each biological replicate was shown.

**Extended Data Figure 16.**
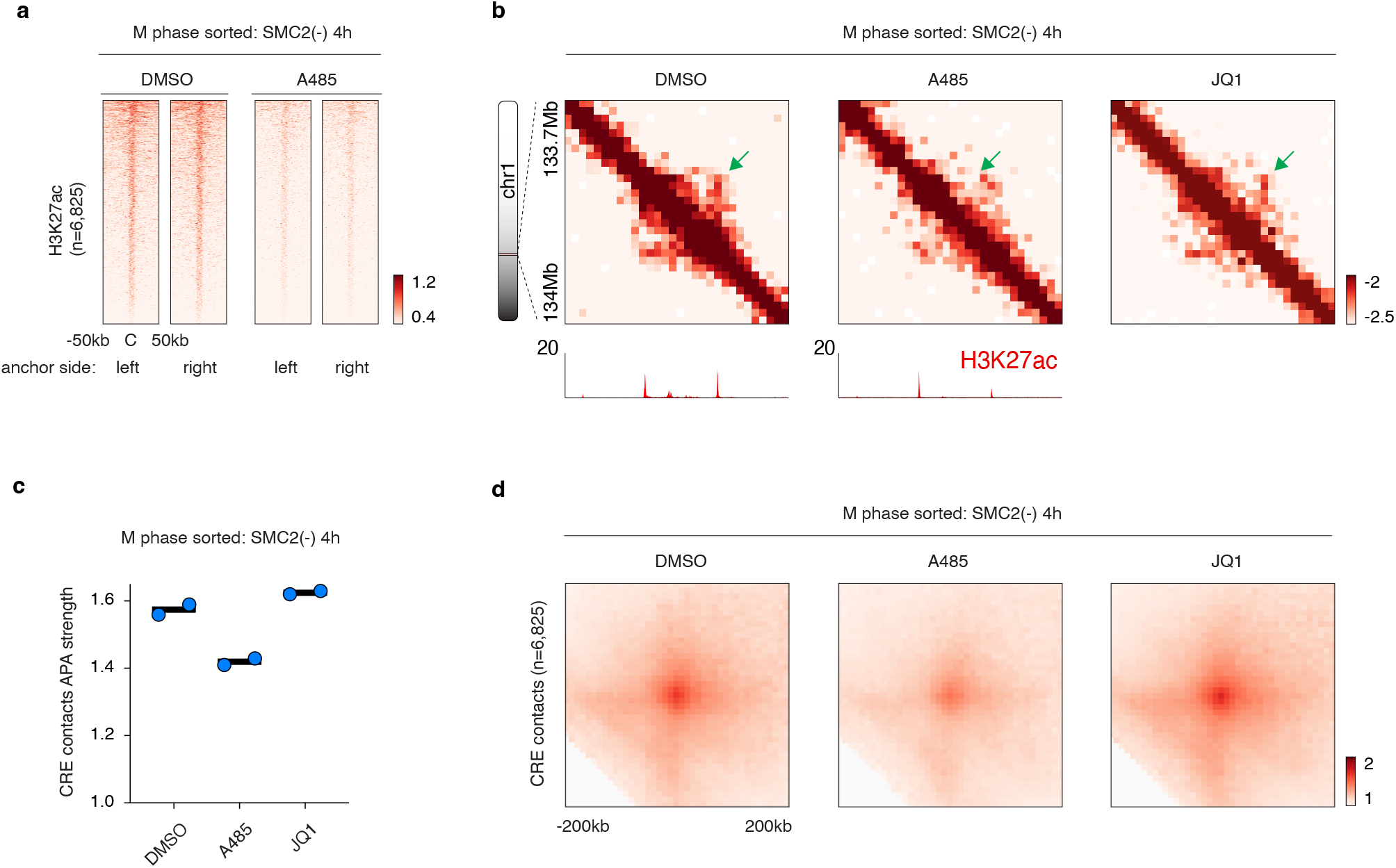
Effects of A485 and JQ1 treatment on mitotic CRE contacts. **a**, Density heatmaps showing mitotic H3K27ac ChIP-seq signals with or without A485 treatment flanking the center of the up-stream or down-stream anchors of CRE contacts. **b**, KR balanced Hi-C contact maps (chr1:133.7-134Mb) for DMSO and A485 treated condensin-deficient (4h) mitotic cells showing a representative mitotic CRE contact. Bin size: 10kb. Tracks of corresponding H3K27ac ChIP-seq results were coupled. **c**, Dot plot showing the APA signals of mitotic CRE contacts upon DMSO, A485 or JQ1 treatment. Each dot represents a biological replicate. **d**, APA plots showing a mild reduction of CRE contact strength in condensin-deficient (4h) mitotic cells after A485 treatment but not after JQ1 treatment.

**Extended Data Figure 17.**
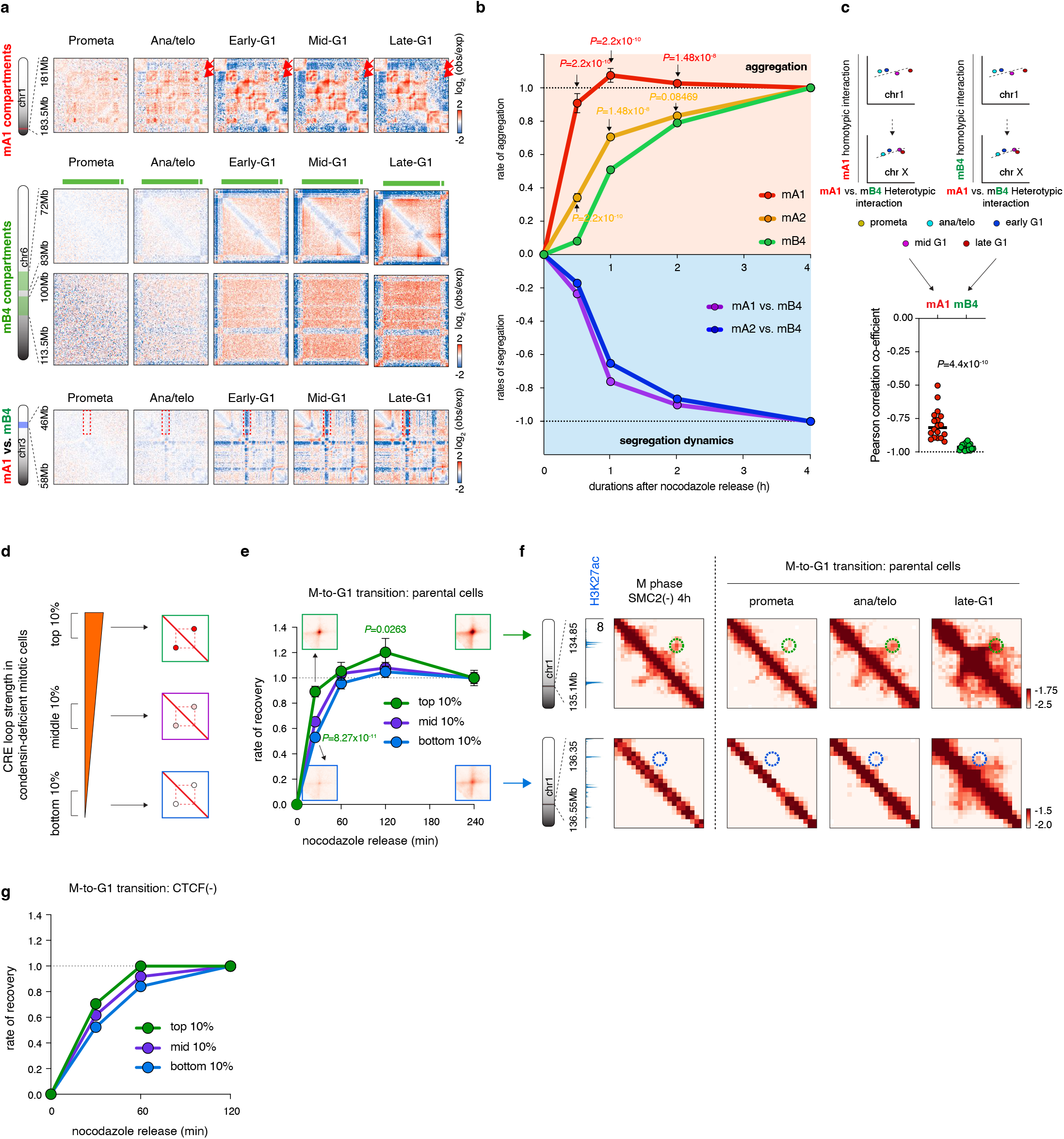
Mitotic mA1 compartmentalization and CRE contacts emerge swiftly in the unperturbed post-mitotic cells. **a**, Upper panel: KR balanced Hi-C contact matrices (chr1:181-183.5Mb) showing rapid emergence of mA1 homotypic interactions in parental G1E-ER4 cells during mitotic exit. Bin size: 25kb. Middle panel: KR balanced Hi-C contact matrices showing the representative (chr6:72-83Mb vs. chr6:100-113.5Mb) delay of the formation of mB4 homotypic interactions. Bin size: 100kb. Lower panel: KR balanced Hi-C contact matrices (chr3:46-58Mb) showing segregation of mA1 from mB4 compartments during mitotic exit. Bin size: 25kb. **b**, Line plots showing the differential reformation kinetics for mA1, mA2 and mB4 as well as the repulsion kinetics between mA1 vs. mB4 and mA2 vs. mB4 in the parental G1E-ER4 cells during mitotic exit. Error-bar denotes SEM (n=18 chromosomes). Statistic tests were performed for comparison of the aggregation dynamics between mA1 vs. mB4 compartments (red) and mA2 vs. mB4 compartments (yellow). *P* values were calculated using a two-sided Wilcoxon signed-rank test. **c**, Left and right: schematics showing how correlations are computed between mA1 or mB4 aggregation strength and mA1 vs mB4 segregation over time. Middle: Dot plot showing the Pearson correlation coefficients between mA1 or mB4 compartment strength and mA1 vs. mB4 segregation in parental cells during mitotic exit. Each dot represents an individual chromosome (n=18). *P* values were calculated using a two-sided Wilcoxon signed-rank test. **d**, Schematic showing the ranking of CRE contacts based on their strength in condensin-deficient mitotic cells. **e**, Line plots showing the faster reformation of the top10% CRE contacts in the parental cells during mitotic exit. APA plots of top and bottom10% CRE contacts in ana/telophase and late-G1 phase were shown. Error bar denotes SEM (n=641). Statistic tests were performed for comparison of the reformation dynamics between top10% and bottom10% CRE contacts. *P* values were calculated using a two-sided Wilcoxon signed-rank test. **f**, Representative KR balanced Hi-C interaction matrices showing the fast reformation of a top10% CRE contact (chr1:134.85-135.1Mb) and the slow reformation of a bottom10% CRE contact (chr1:136.35-136.55Mb) in the parental cells after mitosis. Bin size: 10kb. **g**, Line plots showing the faster reformation of the top10% CRE contacts in the CTCF deficient cells during mitotic exit.

**Extended Data Figure 18.**
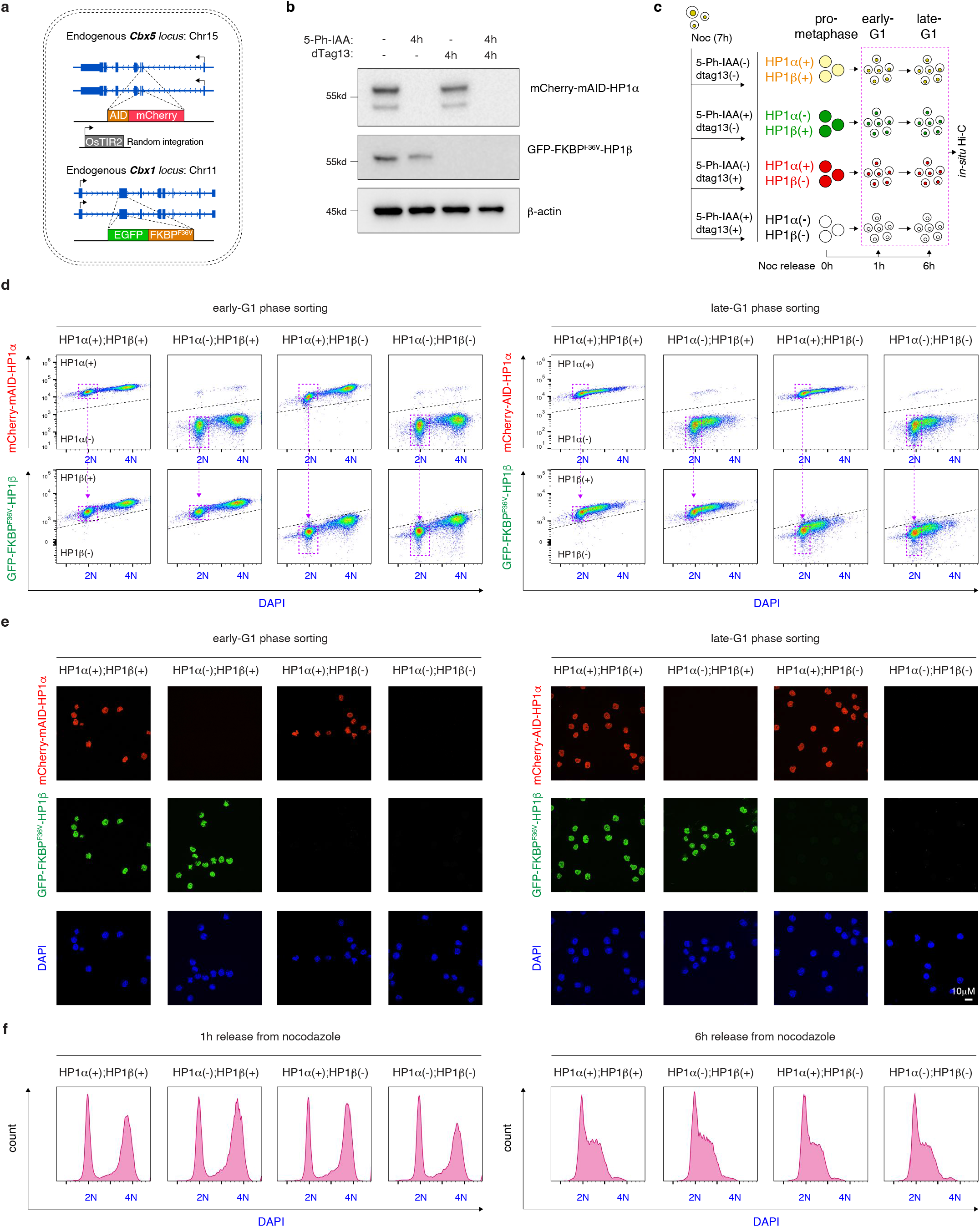
Characterization of the G1E-ER4:mCherry-mAID-HP1α;GFP- FKBP^F36V^-HP1β cell line. **a**, Schematic illustration showing the homozygous insertion of mCherry-mAID tag and GFP-FKBP^F36V^ tag to the N terminus of endogenous *Cbx5* and *Cbx1* locus respectively. **b**, Western blot showing the degradation of HP1α and HP1β upon 5-Ph-IAA or/and dTag13 treatment. One experiment was performed. **c**, Schematic illustration showing the strategy of nocodazole based arrest/release in conjunction with 5-Ph-IAA or/and dTag13 treatment. Early and late-G1 phase cells with four distinct HP1 protein configurations were collected. **d**, Flow cytometry plot showing the sorting strategy of early-G1 and late-G1 phase cells with distinct HP1 protein configurations. Two biological replicates were performed. **e**, Representative confocal images showing the successful depletion of HP1α or HP1β or both in the sorted G1 phase cells. Scale bar: 10*μ*m. Two biological replicates were performed. **f**, Flow cytometry plot showing the mitotic progression of cells under distinct configurations of HP1 proteins after nocodazole release. Two biological replicates were performed.

**Extended Data Figure 19.**
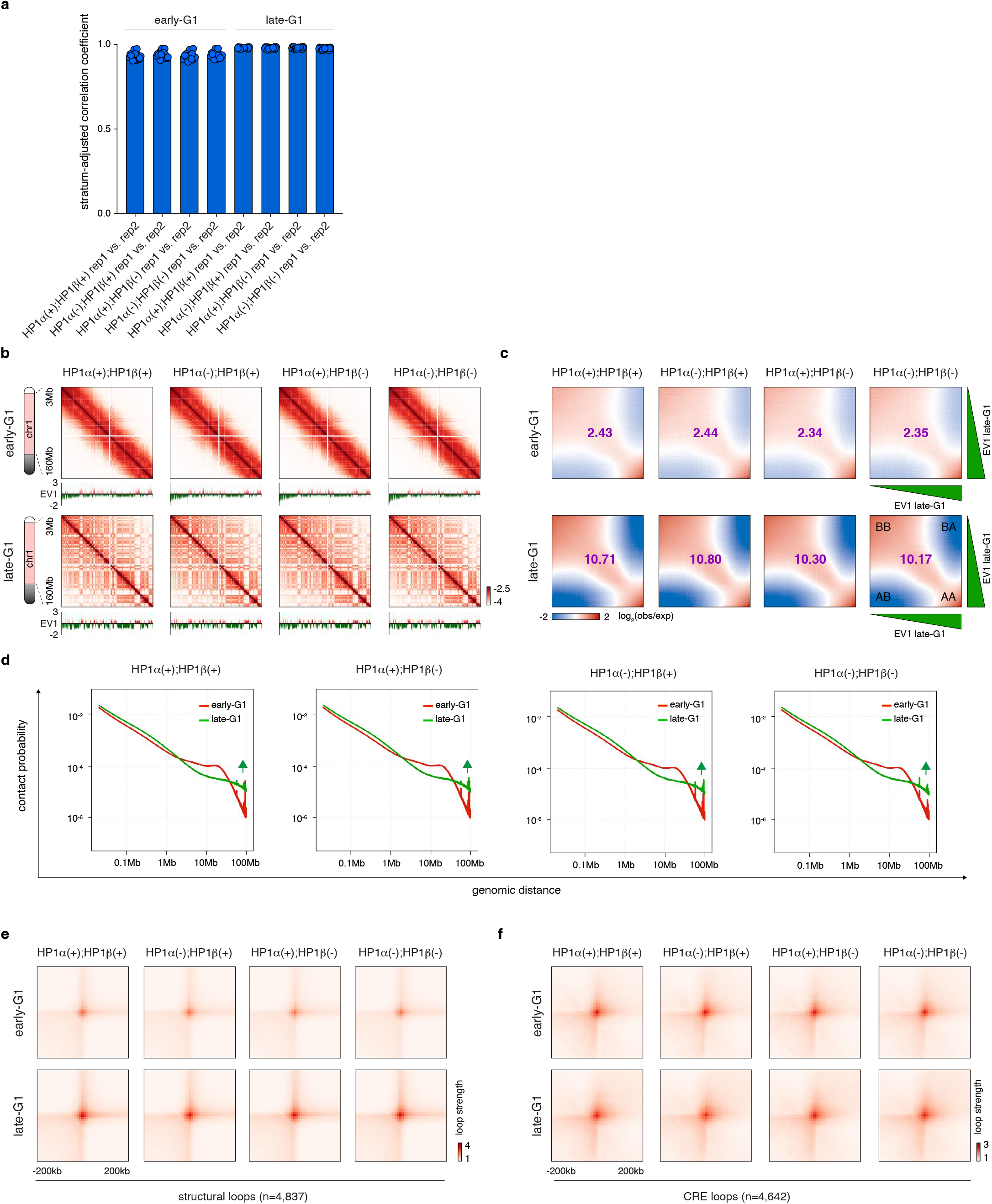
HP1α and HP1β are dispensable for post-mitotic genome refolding. **a**, Bar graph showing the high stratum-adjusted correlation coefficient for each chromosome (n=19) between biological replicates for each condition. **b**, KR balanced Hi-C contact matrices (chr1:3- 160Mb) showing chromatin the compartmentalization in early-G1 and late-G1 cells. Samples with four distinct HP1 protein configurations were shown. Bin size: 100kb. Browser tracks of EV1 values were shown for each contact map. **c**, Saddle-plots showing the progressive compartmentalization of chromatin from early-G1 to late-G1. Samples with four distinct HP1 protein configurations were shown. Compartmental strength for sample were labeled for each plot. **d**, *P(s)* curves for early-G1 and late-G1 phase samples under distinct HP1 protein configurations. **e**, APA plots for structural loop (n=4,837) signals in early-G1 and late-G1 phase samples under distinct HP1 protein configurations. **f**, APA plots for CRE loop (n=4,642) signals in early-G1 and late-G1 phase samples under distinct HP1 protein configurations.

**Extended Data Figure 20.**
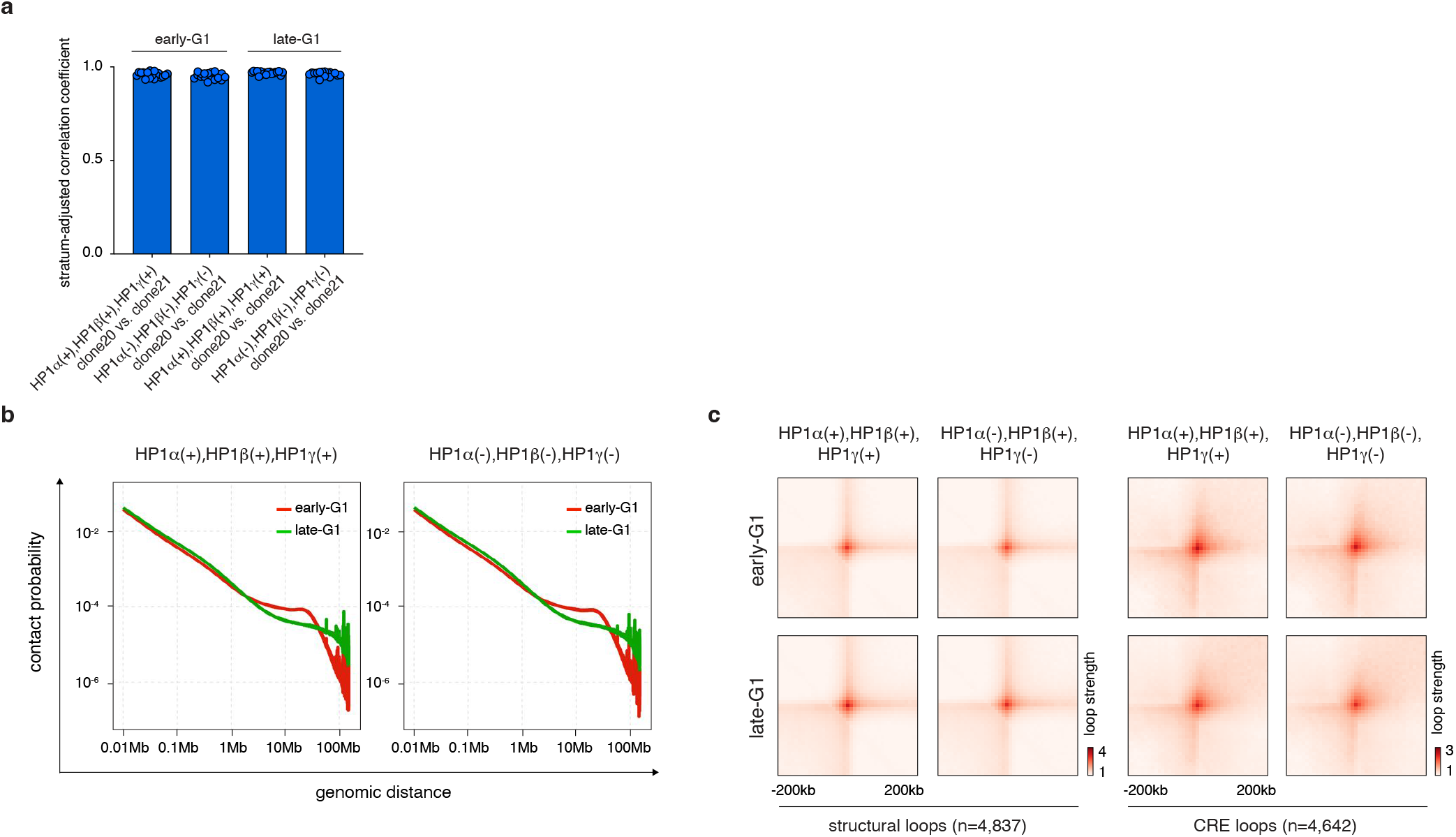
HP1α, HP1β and HP1γ are dispensable for post-mitotic genome refolding. **a**, Bar graph showing the high stratum-adjusted correlation coefficient for each chromosome (n=19) between biological replicates for each condition. **b**, *P(s)* curves for early-G1 and late-G1 phase samples with or without three HP1 proteins. **c**, APA plots for structural loop (n=4,837) and CRE loop (n=4,642) signals in early-G1 and late-G1 phase samples with or without three HP1 proteins.

Supplementary table 1: Hi-C data processing statistics

Supplementary table 2: Arrowhead domain calls

Supplementary table 3: Compartment list

Supplementary table 4: Oligo list

## REFERENCE

1 Nichols, M. H. & Corces, V. G. Principles of 3D compartmentalization of the human genome. Cell reports 35, 109330, doi:10.1016/j.celrep.2021.109330 (2021).

2 Zhang, H. et al. Chromatin structure dynamics during the mitosis-to-G1 phase transition. Nature 576, 158–162, doi:10.1038/s41586-019-1778-y (2019).

3 Zhang, H. et al. CTCF and transcription influence chromatin structure re-configuration after mitosis. Nature communications 12, 5157, doi:10.1038/s41467-021-25418-5 (2021).

4 Abramo, K. et al. A chromosome folding intermediate at the condensin-to-cohesin transition during telophase. Nature cell biology 21, 1393–1402, doi:10.1038/s41556-019-0406-2 (2019).

5 Pelham-Webb, B. et al. H3K27ac bookmarking promotes rapid post-mitotic activation of the pluripotent stem cell program without impacting 3D chromatin reorganization. Molecular cell 81, 1732–1748 e1738, doi:10.1016/j.molcel.2021.02.032 (2021).

6 Zhang, H. & Blobel, G. A. Genome folding dynamics during the M-to-G1-phase transition. Current opinion in genetics & development 80, 102036, doi:10.1016/j.gde.2023.102036 (2023).

7 Gibcus, J. H. et al. A pathway for mitotic chromosome formation. Science 359, doi:10.1126/science.aao6135 (2018).

8 Samejima, K. et al. Functional analysis after rapid degradation of condensins and 3D-EM reveals chromatin volume is uncoupled from chromosome architecture in mitosis. Journal of cell science 131, doi:10.1242/jcs.210187 (2018).

9 Ito, K. & Zaret, K. S. Maintaining Transcriptional Specificity Through Mitosis. Annu Rev Genomics Hum Genet, doi:10.1146/annurev-genom-121321-094603 (2022).

10 Hsiung, C. C. et al. A hyperactive transcriptional state marks genome reactivation at the mitosis-G1 transition. Genes & development 30, 1423–1439, doi:10.1101/gad.280859.116 (2016).

11 Walther, N. et al. A quantitative map of human Condensins provides new insights into mitotic chromosome architecture. The Journal of cell biology 217, 2309–2328, doi:10.1083/jcb.201801048 (2018).

12 Nishimura, K., Fukagawa, T., Takisawa, H., Kakimoto, T. & Kanemaki, M. An auxin-based degron system for the rapid depletion of proteins in nonplant cells. Nat Methods 6, 917–922, doi:10.1038/nmeth.1401 (2009).

13 Campbell, A. E., Hsiung, C. C. & Blobel, G. A. Comparative analysis of mitosis-specific antibodies for bulk purification of mitotic populations by fluorescence-activated cell sorting. Biotechniques 56, 90–94, doi:10.2144/000114137 (2014).

14 Owens, N. et al. CTCF confers local nucleosome resiliency after DNA replication and during mitosis. Elife 8, doi:10.7554/eLife.47898 (2019).

15 Hoencamp, C. et al. 3D genomics across the tree of life reveals condensin II as a determinant of architecture type. *Science (New York*, N.Y.) 372, 984–989, doi:10.1126/science.abe2218 (2021).

16 Spracklin, G. et al. Diverse silent chromatin states modulate genome compartmentalization and loop extrusion barriers. Nature structural & molecular biology, doi:10.1038/s41594-022-00892-7 (2022).

17 Rao, S. S. et al. A 3D map of the human genome at kilobase resolution reveals principles of chromatin looping. Cell 159, 1665–1680, doi:10.1016/j.cell.2014.11.021 (2014).

18 Schwarzer, W. et al. Two independent modes of chromatin organization revealed by cohesin removal. Nature 551, 51–56, doi:10.1038/nature24281 (2017).

19 Rao, S. S. P. et al. Cohesin Loss Eliminates All Loop Domains. Cell 171, 305–320 e324, doi:10.1016/j.cell.2017.09.026 (2017).

20 Behera, V. et al. Interrogating Histone Acetylation and BRD4 as Mitotic Bookmarks of Transcription. Cell reports 27, 400–415 e405, doi:10.1016/j.celrep.2019.03.057 (2019).

21 Xie, L. et al. BRD2 compartmentalizes the accessible genome. Nat Genet 54, 481–491, doi:10.1038/s41588-022-01044-9 (2022).

22 Crump, N. T. et al. BET inhibition disrupts transcription but retains enhancer-promoter contact. Nature communications 12, 223, doi:10.1038/s41467-020-20400-z (2021).

23 Larson, A. G. et al. Liquid droplet formation by HP1alpha suggests a role for phase separation in heterochromatin. Nature 547, 236–240, doi:10.1038/nature22822 (2017).

24 Zenk, F. et al. HP1 drives de novo 3D genome reorganization in early Drosophila embryos. Nature 593, 289–293, doi:10.1038/s41586-021-03460-z (2021).

25 Sanulli, S. et al. HP1 reshapes nucleosome core to promote phase separation of heterochromatin. Nature 575, 390–394, doi:10.1038/s41586-019-1669-2 (2019).

26 Keenen, M. M. et al. HP1 proteins compact DNA into mechanically and positionally stable phase separated domains. Elife 10, doi:10.7554/eLife.64563 (2021).

27 Qin, W. et al. HP1beta carries an acidic linker domain and requires H3K9me3 for phase separation. *Nucleus (Austin*, Tex.) 12, 44–57, doi:10.1080/19491034.2021.1889858 (2021).

28 Vakoc, C. R., Mandat, S. A., Olenchock, B. A. & Blobel, G. A. Histone H3 lysine 9 methylation and HP1gamma are associated with transcription elongation through mammalian chromatin. Molecular cell 19, 381–391, doi:10.1016/j.molcel.2005.06.011 (2005).

29 Spracklin, G. et al. Diverse silent chromatin states modulate genome compartmentalization and loop extrusion barriers. Nature structural & molecular biology 30, 38–51, doi:10.1038/s41594-022-00892-7 (2023).

30 Falk, M. et al. Heterochromatin drives compartmentalization of inverted and conventional nuclei. Nature 570, 395–399, doi:10.1038/s41586-019-1275-3 (2019).

31 Schneider, M. W. G. et al. A mitotic chromatin phase transition prevents perforation by microtubules. Nature 609, 183–190, doi:10.1038/s41586-022-05027-y (2022).

32 Barutcu, A. R., Blencowe, B. J. & Rinn, J. L. Differential contribution of steady-state RNA and active transcription in chromatin organization. EMBO reports 20, e48068, doi:10.15252/embr.201948068 (2019).

33 Lu, J. Y. et al. Homotypic clustering of L1 and B1/Alu repeats compartmentalizes the 3D genome. Cell research 31, 613–630, doi:10.1038/s41422-020-00466-6 (2021).

34 Ma, K. et al. Ribosomal RNA regulates chromosome clustering during mitosis. Cell Discov 8, 51, doi:10.1038/s41421-022-00400-7 (2022).

35 K, L. B., et al. Chromatin proteins and RNA are associated with DNA during all phases of mitosis. Cell Discov 2, 16038, doi:10.1038/celldisc.2016.38 (2016).

36 Meng, Y. et al. The non-coding RNA composition of the mitotic chromosome by 5’-tag sequencing. Nucleic acids research 44, 4934–4946, doi:10.1093/nar/gkw195 (2016).

37 Ginno, P. A., Burger, L., Seebacher, J., Iesmantavicius, V. & Schubeler, D. Cell cycle-resolved chromatin proteomics reveals the extent of mitotic preservation of the genomic regulatory landscape. Nature communications 9, 4048, doi:10.1038/s41467-018-06007-5 (2018).

38 van Steensel, B. & Belmont, A. S. Lamina-Associated Domains: Links with Chromosome Architecture, Heterochromatin, and Gene Repression. Cell 169, 780–791, doi:10.1016/j.cell.2017.04.022 (2017).

39 Erdel, F. et al. Mouse Heterochromatin Adopts Digital Compaction States without Showing Hallmarks of HP1-Driven Liquid-Liquid Phase Separation. Molecular cell 78, 236–249 e237, doi:10.1016/j.molcel.2020.02.005 (2020).

40 Saksouk, N. et al. The mouse HP1 proteins are essential for preventing liver tumorigenesis. Oncogene 39, 2676–2691, doi:10.1038/s41388-020-1177-8 (2020).

41 Feng, Y. et al. Simultaneous epigenetic perturbation and genome imaging reveal distinct roles of H3K9me3 in chromatin architecture and transcription. Genome Biol 21, 296, doi:10.1186/s13059-020-02201-1 (2020).

42 Yang, T. et al. HiCRep: assessing the reproducibility of Hi-C data using a stratum-adjusted correlation coefficient. Genome research 27, 1939–1949, doi:10.1101/gr.220640.117 (2017).

43 Dogan, N. et al. Occupancy by key transcription factors is a more accurate predictor of enhancer activity than histone modifications or chromatin accessibility. Epigenetics Chromatin 8, 16, doi:10.1186/s13072-015-0009-5 (2015).

44 Wu, W. et al. Dynamic shifts in occupancy by TAL1 are guided by GATA factors and drive large-scale reprogramming of gene expression during hematopoiesis. Genome research 24, 1945–1962, doi:10.1101/gr.164830.113 (2014).

